# How Well Can We Detect Shifts in Rates of Lineage Diversification? A Simulation Study of Sequential AIC Methods

**DOI:** 10.1101/011452

**Authors:** Michael R. May, Brian R. Moore

## Abstract

Evolutionary biologists have long been fascinated by the extreme differences in species numbers across branches of the Tree of Life. This has motivated the development of statistical phylogenetic methods for detecting shifts in the rate of lineage diversification (speciation – extinction). One of the most frequently used methods—implemented in the program `MEDUSA`—explores a set of diversification-rate models, where each model uniquely assigns branches of the phylogeny to a set of one or more diversification-rate categories. Each candidate model is first fit to the data, and the Akaike Information Criterion (AIC) is then used to identify the optimal diversification model. Surprisingly, the statistical behavior of this popular method is completely unknown, which is a concern in light of the poor performance of the AIC as a means of choosing among models in other phylogenetic comparative contexts, and also because of the *ad hoc* algorithm used to visit models. Here, we perform an extensive simulation study demonstrating that, as implemented, `MEDUSA` (1) has an extremely high Type I error rate (on average, spurious diversification-rate shifts are identified 42% of the time), and (2) provides severely biased parameter estimates (on average, estimated net-diversification and relative-extinction rates are 183% and 20% of their true values, respectively). We performed simulation experiments to reveal the source(s) of these pathologies, which include (1) the use of incorrect critical thresholds for model selection, and (2) errors in the likelihood function. Understanding the statistical behavior of `MEDUSA` is critical both to empirical researchers—in order to clarify whether these methods can reliably be applied to empirical datasets—and to theoretical biologists—in order to clarify whether new methods are required, and to reveal the specific problems that need to be solved in order to develop more reliable approaches for detecting shifts in the rate of lineage diversification.

---

Many evolutionary phenomena entail differential rates of diversification (speciation – extinction); *e.g.*, adaptive radiation, diversity-dependent diversification, key innovations, and mass extinction. Phylogeny-based statistical methods have been developed to detect shifts in diversification rate *through time*, such as tree-wide shifts in diversification rate associated with episodes of mass extinction or adaptive radiation (Stadler 2010; 2011; Morlon et al. 2011), or diversity-dependent decreases in diversification rate associated with ecological limits on speciation (Rabosky 2006; Etienne et al. 2012). Other methods seek to identify *correlations* between rates of diversification and some other variable, such as the evolution of discrete or continuous traits (Maddison et al. 2007; FitzJohn 2010) or episodes of biogeographic or climatic change (Moore and Donoghue 2009; Goldberg et al. 2011). Here, we focus on a third class of methods that seek to detect shifts in diversification rate *along lineages* of a phylogenetic tree (Moore et al. 2004; Chan and Moore 2005; Rabosky et al. 2007; Alfaro et al. 2009).

The study of diversification-rate shifts along lineages is most often pursued using the approach proposed by Alfaro et al. (2009). In fact, this approach—Modeling Evolutionary Diversification Using Stepwise AIC (`MEDUSA`)—is used more frequently than *any* other statistical phylogenetic method associated with lineage diversification (S.3). This approach assumes that phylogenies are generated under a birth-death stochastic-branching process model with one or more diversification-rate categories. The `MEDUSA` algorithm proceeds by first fitting a series of increasingly complex models with 1, 2,…, *j* diversification-rate categories; each with unique speciation, *λ*, and extinction, *µ* rate parameters. Each diversification model uniquely specifies both the number of rate categories and the assignment of those rates to branches of the phylogeny. After identifying the best diversification model for each category, `MEDUSA` then selects among the *j* diversification models using standard model-selection methods (the Akaike Information Criterion, AIC; Akaike 1974).

The popularity of `MEDUSA` stems from several advantages it holds over alternative approaches: (1) rather than requiring complete species-level phylogenies, `MEDUSA` allows unsampled species to be included within unresolved terminal subclades; (2) rather than requiring the location of diversification-rate shifts to be specified *a priori*, `MEDUSA` agnostically evaluates diversification-rate shifts along all branches of the tree, allowing this problem to be studied within an exploratory-data analysis framework; (3) rather than assuming a pure-birth (Yule) stochastic-branching process model, `MEDUSA` is based on a more realistic birth-death model that accommodates extinction, and; (4) in addition to inferring the location(s) of diversification-rate shifts, `MEDUSA` also provides estimates of the diversification-rate parameters for each branch of the tree.

Surprisingly, the statistical behavior of this popular method is completely unknown, and, in fact, the algorithm has never been formally described in the literature. This is particularly troubling, as the reliability of the AIC for model selection is known to be problematic in other phylogenetic contexts (Alfaro and Huelsenbeck 2006; Boettiger et al. 2012), and because three separate variants of the algorithm are now available to users. Simulation is a critical tool for validating an inference method, as it can uniquely assess the ability of an approach to recover the true/known parameter values that were used to generate the data. These controlled experiments allow us to characterize the statistical behavior of a given method. Simulation can either confirm that a method will provide reliable inferences when applied to empirical data (where the actual parameter values are unknown), or if it is found to be unreliable, simulation can be used to understand why the method fails.

With these considerations in mind, we first provide a formal description of the `MEDUSA` algorithm, and then perform an extensive simulation study to characterize the statistical behavior of the stepwise AIC method on simulated phylogenies. Our primary goals are (1) to reveal whether `MEDUSA` provides reliable estimates of diversification-rate shifts, and, if not (2) to understand *why* these methods are unreliable. Therefore, our simulation study is presented in the manner in which it was conducted: as an autopsy where (after determining that `MEDUSA` has fatal issues), we perform a series of simulation experiments to understand the nature of the methodological pathologies.

## Inferring Diversification-Rate Shifts with Medusa

### Data

We treat the inferred phylogeny—including the tree topology and divergence times—as observations. Phylogenies may be *complete*, where each tip in the tree represents a single species. Alternatively, the phylogeny may be *incomplete*, in which case the tips represent terminally unresolved lineages. The latter *species-diversity trees* entail additional assumptions: the monophyly, age, and species diversity of all terminal lineages are assumed to be precisely known without error. More formally, the data comprise a rooted ultrametric tree with *k* terminal lineages denoted as Ψ = {*τ*, **t**, **n**}, where *τ* is the tree topology, **t** is a vector of divergence times starting from the root, and **n** is a vector of species-richness values for each of the *k* terminal lineages.

### Likelihood Function and Diversification-Rate Models

We assume the phylogeny, Ψ, was generated by a birth-death stochastic-branching process with a vector of branch-specific speciation rates, ***λ*** = {*λ*_1_, *λ*_2_,…, *λ*_2*k*−2_}, and a vector of branch-specific extinction rates, ***µ*** = {*µ*_1_, *µ*_2_,…, *µ*_2*k*−2_}, where *λ_i_* and *µ_i_* are the speciation and extinction rates for branch *i*, respectively.

#### Likelihood function

The likelihood of observing the data under the stochastic-branching process model is calculated piecewise (*c.f.*, Rabosky et al. 2007). We first compute the likelihood of the internal branches, then compute the likelihood of the terminal lineages, and finally combine these two partial likelihoods. The probability of observing internal branch of length *t*_*i*_, conditional on it having descendants at the present, is

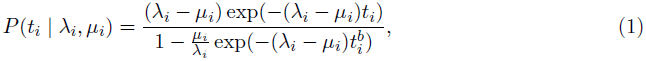

where 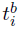 is the birth-time of the branch (Nee et al. 1994; equation 17). The probability of observing terminal lineage *i* with *n*_*i*_ species, conditional on it having descendants at the present (*n*_*i*_ > 0) is

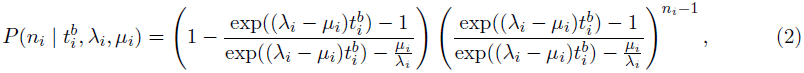

which follows from (Raup 1985; equation A17). Parsing the tree into *𝒯* terminal lineages and *𝓘* internal branches, the likelihood for the phylogeny and species-richness data is calculated as

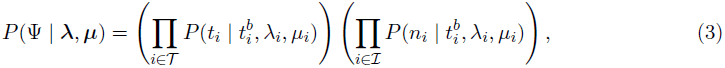

as proposed by Rabosky et al. (2007).

#### Diversification-rate models

The `MEDUSA` algorithm assumes that diversification-rate parameters are inherited identically over the tree unless a shift in diversification rate occurs along a branch. Every tree will have a “background” diversification rate (with a pair of rate parameters, *λ*_0_ and *µ*_0_). Each of the 2*k* − 2 branches on the tree can experience one or zero shifts in diversification rate. In principle, the diversification rate may change at any point(s) along a given branch. In practice, however, the `MEDUSA` algorithms assume that a single shift occurs either at the very beginning of a branch (*i.e.*, immediately after the speciation event that gave rise to the branch), or at the very end of a branch (*i.e.*, immediately before the branch experiences speciation). In the former ‘*stem*’ rate-shift scenario, the branch inherits the new diversification-rate parameters; in the latter ‘*crown*’ rate-shift scenario, the branch retains the ancestral diversification-rate parameters. Each of the *i* rate shifts adds a new diversification-rate category (with rate parameters *λ*_*i*_ and *µ*_*i*_). Models that share the same number of diversification-rate categories, *i*, belong to the same model index, which we denote *M*_*i*_. Additionally, we use the notation 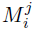 to indicate the set of *j* diversification-rate models that are members of model index *i*.

To illustrate our notation, consider the simple tree in Figure 1: the rooted, binary species-diversity tree has *k* = 4 terminal lineages and (2*k* − 2) = 6 branches. Accordingly, this tree can experience *i* = {0, 1,…, 4} crown diversification-rate shifts, each with a corresponding model index *M_i_*. Within a given model index, *M_i_*, each of the *j* distinct locations of the *i* rate shifts on branches represents a unique diversification-rate model, 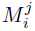. For example, there is only a single model, 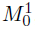, where all branches diversify under the background diversification rate (with parameters *λ*_0_ and *µ*_0_). By contrast, there are six distinct diversification-rate models (*i.e.*, 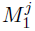, where *j* = {1, 2,…, 6}) that variously assign the single diversification-rate shift to one of the six branches in the tree.

The state space of possible diversification-rate models for a tree with *b* = (2*k* − 2) branches is described by the corresponding Bell number (Bell 1934). The Bell number for *b* branches is the sum of the Stirling numbers of the second kind:

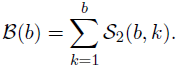

The Stirling number of the second kind, *𝒮*_2_(*b*, *k*), for *b* elements and *k* subsets (corresponding here to the number of branches and diversification-rate categories, respectively) is given by the following equation:

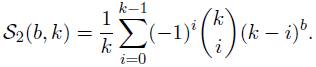

The state space of possible diversification-rate models quickly becomes large, even for small trees. For example, the simple tree in Figure 1 has *𝓑*(6) = 203 possible diversification-rate models, a species-diversity tree with six lineages has *𝓑*(10) = 115, 975 possible diversification-rate models, a tree with ten lineages has *𝓑*(18) = 682, 076, 806, 174 possible models, and a tree with 50 lineages has *𝓑*(98) = 3.12 × 10^112^ possible models. Clearly, the space of candidate diversification models quickly becomes vast, which motivates the consideration of heuristic algorithms that might allow us to efficiently explore this space.

### The MEDUSA Algorithm

The essence of the `MEDUSA` algorithm is straightforward: it explores the space of diversification-rate models, estimates the maximum likelihood for each candidate model, and then uses AIC to select the optimal diversification-rate model. The potentially vast solution space, however, requires heuristic techniques to efficiently search a small subspace of all possible diversification-rate models. An exhaustive algorithm (one that systematically evaluates every possible diversification-rate model) is prohibitive even for simple problems. Consider, for example, a species-diversity tree with ten lineages: even if we could compute the maximum-likelihood estimate for each candidate model at a rate of one per second, it would require ∼ 21, 628 years to evaluate every diversification-rate model. Although critical, heuristic shortcuts introduce considerable complexity into the `MEDUSA` algorithm. We provide a relatively detailed description of the `MEDUSA` algorithm in the Supplemental Material, and illustrate the basic procedure here with a simple example.

Consider the hypothetical species-diversity tree with four terminal lineages depicted in Figure 1. `MEDUSA` evaluates diversification-rate models in order of increasing complexity; *M*_0_, *M*_1_…, *M_max_*. Accordingly, the algorithm begins by fitting a constant-rate model (*i.e.*, *M*_0_, with zero rate shifts) to the tree and species-richness data: numerical optimization is used to identify the rate-parameter values, 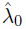 and 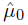, that maximize the likelihood function (equation 3). The resulting maximum-likelihood estimate, *𝓛*, under the one-rate model is then used to compute its AIC score: AIC = 2*p* − 2 ln *𝓛*, where *p* is the number of free parameters in the model (Akaike 1974).

The `MEDUSA` algorithm then evaluates each of the diversification models with one rate-shift, 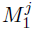 (where *j* = 6 diversification-rate models with two pairs of rate parameters, *λ*_0_, *µ*_0_, and *λ*_1_, *µ*_1_). The two-rate model that best fits the data (*i.e.*, with the highest log likelihood) is selected, and its AIC score is computed. We then assess the relative improvement in the fit of the two-rate model over the one-rate model as the difference in their respective AIC scores: ∆AIC=AIC_one-rate_−AIC_two-rate_. If the computed ∆AIC is greater than some pre-specified threshold, ∆AIC_crit_, `MEDUSA` accepts the best two-rate model and the corresponding location of the diversification-rate shift specified by that two-rate model.

The algorithm then identifies the best three-rate model that incorporates the previously inferred rate shift, and accepts or rejects it over the two-rate model based on the same ∆AIC_crit_ threshold. The algorithm continues, incrementing the model index until either (1) the most complex model is accepted or (2) the improvement in the AIC score is too small to exceed the ∆AIC_crit_ threshold. In principle, the most complex model index is a function of the number of lineages, *M_max_* = 2*k* − 2. In practice, however, *M_max_* is set to an arbitrary value that limits the evaluation to a subset of all possible models (20 shifts in the original algorithm, but the number varies by implementation). Note that the `MEDUSA` algorithm tremendously reduces the scope of candidate diversification-rate models: in the present case—a species-diversity tree with four lineages—the space collapses from 203 to 16 candidate models.

Different implementations of `MEDUSA` specify the ∆AIC_crit_ differently. The original implementation (Alfaro et al. 2009) uses the conventional value for the ∆AIC_crit_ threshold, which is fixed at 4. More recent versions of the algorithm specify a critical threshold that is a function of the number of terminal lineages. Specifically, both `turboMEDUSA` (implemented both as a stand-alone R package and in `GEIGER >1.99`) and a newer version of `MEDUSA` (available at http://github.com/josephwb/turboMEDUSA) compute the ∆AIC_crit_ threshold based on the number of terminal lineages (the latest versions prohibit negative ∆AIC_crit_ values). An example of the three different ∆AIC_crit_ thresholds (corresponding to the parameter space that we explore in one of our simulations) is depicted in Figure 3. We refer to the original `MEDUSA`, `turboMEDUSA` and the newest version of `MEDUSA` algorithms as `oMEDUSA`, `tMEDUSA` and `nMEDUSA`, respectively.

## Simulation Study

Our motivation for exploring the statistical behavior of the `MEDUSA` algorithm stems from its reliance on the Akaike Information Criterion (AIC) to choose among candidate diversification-rate models. Use of the AIC to choose among models in other phylogenetic settings is known to be problematic (*e.g.*, Alfaro and Huelsenbeck 2006; Boettiger et al. 2012). Specifically, there is little theory to guide the specification of an appropriate critical threshold for preferring one model to another (*i.e.*, the difference in AIC scores between the two competing models, ∆AIC_crit_), and arbitrarily specified thresholds may strongly bias the model-selection procedure. Moreover, the AIC assumes that the sample size (in this case, the number of species/terminal lineages) is large. In practice, however, `MEDUSA` is generally applied to trees with a small number of incompletely sampled terminal lineages. Accordingly, the large-sample size assumption of the AIC is apt to be violated, which may compromise the ability of `MEDUSA` to reliably choose the correct model. With these considerations in mind, we performed a series of simulations to determine how well `MEDUSA` recovers the correct diversification-rate model under a variety of circumstances. All simulations were performed in `R` (R Core Team 2013) using the packages `ape` and `TreeSim` (Paradis et al. 2004; Stadler 2013); trees were subsequently manipulated with custom `R` scripts. All simulated data as well as the `R` scripts used to generate and analyze the simulated data are available from the Dryad Digital Repository: http://dx.doi.org/10.5061/dryad.261v1.

### Type I Error Rate

We explored the Type I error rate of `MEDUSA` by simulating trees under a constant-rate birth-death process (*i.e.*, where the diversification rate is constant across lineages), and then analyzed these trees with `MEDUSA` to assess the frequency with which it incorrectly inferred a diversification-rate shift (*i.e.*, the false-positive rate). We simulated constant-rate trees over a wide range of diversification parameters (net diversification, *r* = *λ* − *µ*, and relative extinction, *∈* = *µ* ÷ *λ*), tree sizes, *N*, and number of unresolved terminal lineages, *k*. The set of relative-extinction rates, *∈*, that we explored span the possible range of values (from a pure-birth process with no extinction, to scenarios with an extreme extinction load). Our choice of parameters for tree size, *N*, and the number of terminally unresolved clades, *k*, reflects the dimensions of empirical datasets to which `MEDUSA` has been applied (Table S.1). We note that the algorithm as implemented in the most recent version `GEIGER > 1.99`, which most closely resembles `tMEDUSA`, is both considerably faster than `oMEDUSA` and is also the version recommended by the developers of these methods (Luke Harmon, personal communication). We have nevertheless opted to examine the behavior of all three iterations of the `MEDUSA` algorithm, as most of the empirical applications have used earlier versions of the method (51.7%, 46.6%, and 1.7% of empirical studies have used `oMEDUSA`, `tMEDUSA` and `nMEDUSA`, respectively; Table S.1).

#### The effect of relative-extinction rate, tree size, and phylogenetic resolution

In the first set of simulations, the absolute speciation rate was *λ* = 0.01 and the extinction rate was incremented over a range of values, *µ* = {0.000, 0.001, 0.002,…, 0.009} (see Table S.2 for corresponding composite parameters, *r* and *∈*). For each value of *µ*, we simulated 1, 000 trees of three sizes, *N* = {100, 1, 000, 10, 000}. We then created terminally unresolved clades in each tree by identifying the time when there were *k* lineages, for *k* = {10, 15, 20, 25, 30, 40, 100}, pruning out all subsequent speciation events, and then assigning the corresponding species-richness values to each terminal lineage. In total, we generated 210, 000 trees with species-richness data (1, 000 replicates for each combination of *µ*, *N*, and *k*). We then analyzed each simulated tree using the three variants of the `MEDUSA` algorithm. Note that the original algorithm, `oMEDUSA`, assumes that diversification-rate shifts occur along stem branches, whereas the two more recent algorithms, `tMEDUSA` and `nMEDUSA`, allow the user to specify whether rate shifts are assumed to occur along crown or stem branches. To enhance comparability, we focus on results where we assumed that diversification-rate shifts occur along stem branches. However, we have replicated all of the analyses under the alternative option, where diversification-rate shifts were assumed to occur along crown branches (these results are presented in the Supplementary Material).

All three of the `MEDUSA` algorithms use the conventional `optim` algorithm (Nelder and Mead 1965) to estimate the vector of parameter values (***λ*** and ***µ***) that collectively maximize the likelihood of the phylogenetic observations, Ψ, under the corresponding diversification-rate model, 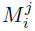. In general, these numerical algorithms begin by first specifying an initial value for each parameter, and then calculate the log-likelihood for the vector of initial parameter values (by evaluating equation 3). New parameter values are iteratively proposed and evaluated until any possible change to parameter values decreases the log-likelihood, at which point the algorithm has either converged to a local or the global maximum-likelihood estimate, and the procedure terminates.

Our preliminary analyses using default initial parameter values revealed that the numerical algorithms used for maximum-likelihood estimation in `MEDUSA` frequently failed in one of two ways. Specifically, 25% of the analyses experienced ‘*hard*’ failures, where the program terminated with an error message indicating that the maximum log-likelihood could not be calculated. More troubling was the incidence of ‘*soft*’ failures, where analyses that appeared to terminate successfully actually failed to converge to the maximum-likelihood estimate. To control for the potentially confounding effect of this numerical instability on our evaluation of the statistical behavior of `MEDUSA`, we initialized analyses with parameters set to their true values (the parameter values used to simulate the data). Because the true parameter values are unknown for empirical datasets, our analyses of the simulated data will provide a relatively favorable assessment of the performance of the methods.

Results of the first simulation are summarized in Figure 2 (see also Table S.2). We can view Figure 2 as a graphical table with nine panels: the three panels in each row summarize results for one algorithm (`oMEDUSA`, `tMEDUSA`, and `nMEDUSA`, from top to bottom), and the three panels in each column summarize results for one tree size (with 100, 1, 000, and 10, 000 species, from left to right). The overall Type I error rate—calculated as the unweighted average over all of the cells within each of the nine panels—is 40.2%, which is ∼ 8 times higher than the nominal significance level, *α* = 0.05. The average Type I error rate varies over rows of Figure 2: 51%, 29%, 41% for the progressively newer `oMEDUSA`, `tMEDUSA`, and `nMEDUSA` algorithms, respectively. Additionally, the overall Type I error rate tends to increase across each column of Figure 2: 26%, 47%, 48% for increasingly larger trees.

The relatively clear patterns manifest among columns/rows of panels in Figure 2 contrasts with the complex patterns of Type I error rate observed *within* each of the panels. That is, trends in the Type I error rate within panels differed qualitatively between different versions of the algorithm. For example, Type I error rates for `tMEDUSA` tended to increase with the relative-extinction rate, *∈*, and decrease with the number of terminal lineages, *k*, (increasing from the lower right to the upper left of each panel). By contrast, Type I error rates for `nMEDUSA` tended to increase with the relative-extinction rate and the number of terminal lineages (increasing from the lower left to the upper right of each panel). The source of the disparate behavior of the `tMEDUSA` and `nMEDUSA` algorithms is unclear, as they are said to be identical (Harmon et al. 2008), except that `nMEDUSA` does not allow negative ∆AIC_crit_ values (*c.f.*, Figure 3). Finally, patterns of Type I error rate within panels for `oMEDUSA` did not exhibit any clear correlation with the relative-extinction rate or the number of lineages, but appear to be some complex function of these two variables.

We first considered the possibility that the complex patterns of Type I error rate might be an artifact of Monte Carlo error (*i.e.*, error variance caused by simulating insufficient replicates). However, our experiments using an increased number of replicates exclude this explanation; the high level and complex patterns of Type I error are estimated precisely (Figure S.4, Table S.3). Below, we describe our efforts to first rule out the possibility that the high Type I error rates are artifactual, and then to understand the possible source(s) of the high Type I error rates.

#### The effect of absolute diversification rates

Could the high the Type I error rates inferred in the initial simulation be an artifact of the absolute parameter values that we used? Although our choice of diversification rates was based on a survey of ‘normal’ empirical values, it is possible that the `MEDUSA` algorithm might commonly be applied to groups with anomalously high diversification rates. To explore this possibility, we performed a second series of constant-rate simulations to assess the effect of increasing the absolute speciation rate, *λ*, on the statistical behavior of `MEDUSA`. Specifically, we simulated trees under a five-and ten-fold higher speciation rate, *λ* = {0.05, 0.1}. For each speciation rate, we first simulated a set of trees under a range of extinction rates, *µ*, such that the corresponding relative-extinction rates, *∈*, were identical to those of the initial simulation; *∈* = {0.0, 0.1,…, 0.9} (Table S.4). For each absolute speciation rate, we also simulated a second set of trees under a range of extinction rates, *µ*, such that the corresponding net-diversification rates, *r*, were identical to those used in the initial simulation, *r* = {0.001, 0.002,…, 0.01} (Table S.5). We simulated 1, 000 trees with *N* = 10, 000 species for each combination of parameters. As in the initial simulation, we created a set of *k* = {10, 15, 20, 25, 30, 40, 100} terminally unresolved lineages in each tree, resulting in 7 × 4 × 1, 000 = 28, 000 trees with species-richness data. We then analyzed each tree using all three implementations of the `MEDUSA` algorithm. Again, we repeated the entire series of analyses evaluating ‘stem’ and ‘crown’ diversification-rate shifts.

Results of these simulations are summarized in Figures S.5 and S.6 (see also Tables S.4–S.5). Overall Type I error rates were slightly higher than those of the initial simulations: 40.2%, 44.7%, and 44.7% for simulations based on the initial, five-and ten-fold higher absolute rates, respectively. The relative performance of the three `MEDUSA` algorithms was also similar to the initial simulation: `oMEDUSA`, `tMEDUSA`, and `nMEDUSA` exhibited Type I error rates of 44%, 40%, and 51%, respectively. Moreover, these simulations cast light on the impact of relative extinction. For the first set of higher rate simulations—where the relative-extinction rates matched those of the initial simulation—the overall rate and patterns of Type I error were similar to those based on the initial lower-rate simulations (37.4% *vs.* 40.2%, respectively; Figure S.5). By contrast, the second set of higher-rate simulations—where the net-diversification rates matched those of the initial simulation—imposed a narrow range of relatively high relative-extinction rates, *∈* = {0.80, 0.82, 0.84…, 0.98}. For these simulations, the overall Type I error rate increased substantially compared to the initial lower-rate simulation (51.5% *vs.* 40.2%, respectively; Figure S.6).

In summary, the high Type I error rates identified in the first simulation do not appear to be an artifact of the chosen parameter values, and appear to be strongly influenced by the severity of the relative-extinction rate.

#### Exploring threshold effects on Type I error rates

Given that the high Type I error rates revealed in the previous simulations are real—they are not artifacts of Monte Carlo error or the specific parameter values that we used—we might ask whether these errors reflect an excess of “near misses”, where there is only a *slight* preference for the (incorrect) multi-rate model over the (true) constant-rate model. To explore this possibility, we examined the ∆AIC values computed for the parameter space explored in the first simulation (Figure 3).

Surfaces of computed ∆AIC values for the original algorithm, `oMEDUSA`, are both extreme and extremely complex. For smaller and intermediate trees, with *N* = 100 and *N* = 1, 000 species, the computed ∆AIC scores—reflecting the difference between the AIC scores for the (correct) rate-constant and the (incorrect) multi-rate models—increases as a complex function of the relative-extinction rate, *∈*, and the number of terminal unresolved lineages, *k*. Moreover, the preference for the (incorrect) multi-rate model is typically very strong; the computed ∆AIC values are up to 2500 times the fixed critical threshold (where ∆AIC_crit_ = 4). Accordingly, the high Type I error rates for the `oMEDUSA` algorithm reflect a preponderance of “wild guesses” rather than “near misses”. The pathological behavior of the `oMEDUSA` algorithm is compounded by the qualitatively different (but equally complex) surface of computed ∆AIC values for the largest trees (with *N* = 10, 000 species). Clearly, the fixed threshold used by `oMEDUSA` is not a viable approach for assessing the relative fit of (and selecting among) diversification models.

Surfaces of computed ∆AIC values for the newer `tMEDUSA` and `nMEDUSA` algorithms contrast sharply with the erratic behavior of the `oMEDUSA` algorithm. For small trees, with *N* = 100 species, the computed ∆AIC surfaces conform more uniformly to the corresponding critical thresholds. Accordingly, many of the false-positive cases based on `tMEDUSA` and `nMEDUSA` analyses of the smallest trees correspond to “near-ish misses”, where the computed ∆AIC *moderately* exceeds the ∆AIC_crit_. By contrast, for larger trees, with *N* = 1, 000 and 10, 000 species, surfaces of computed ∆AIC values for both algorithms greatly exceed the corresponding surface of critical thresholds, ∆AIC_crit_. As was generally the case with the `oMEDUSA` algorithm, false-positive cases based on `tMEDUSA` and `nMEDUSA` analyses of larger trees represent “wild guesses”, where the (true) constant-rate model is decisively rejected in favor of the (incorrect) multi-rate model.

In summary, support for the incorrect multi-rate model generally increases with tree size, *N*. This observation suggests two possible explanations. The first attributes the correlation between tree size and Type I error rate to power: constant-rate models may be rejected less frequently in small trees owing to a simple lack of statistical power to detect (spurious) diversification-rate shifts. The alternative explanation focuses on the relative proportion of the tree included in terminal unresolved lineages. Consider plots of the computed ∆AIC surfaces for `tMEDUSA` and `MEDUSA` (Figure 3). For smaller trees (with *N* = 100 species), the degree to which the computed ∆AIC surface exceeds the ∆AIC_crit_ threshold is a function of the number of terminal unresolved lineages, *k*. Note that we use the same set of terminal unresolved lineages in all simulations; *k* = {10, 15, 20, 25, 30, 40, 100}. The proportion of the tree relegated to terminal unresolved lineages therefore depends on tree size. For example, when *k* = 100 and *N* = 100, the tree is 100% resolved (each terminal lineage includes a single species), but when *k* = 100 and *N* = 1, 000, the tree is only 10% resolved (900 species are apportioned among the 100 terminal unresolved lineages). These observations suggest that the high Type I error rate for the `MEDUSA` algorithms may be related to terminal unresolved lineages. We evaluate this possible explanation in the next section by estimating the Type I error rates for `MEDUSA` based on analyses of completely resolved trees.

#### Exploring Type I error rates for complete species trees

In order to control for the apparently complex effects of terminally unresolved lineages on Type I error rate, we performed a series of simulations focused on complete species trees (*i.e.*, trees in which each tip represents a single species). Specifically, we simulated complete trees of seven sizes, *N* = {50, 100, 150, 200, 250, 500, 1, 000}, under a constant-rate birth-death process where the speciation rate, *λ*, was held at 0.01, and the extinction rate, *µ*, was incremented over a range of ten values, *µ* = {0.000, 0.001, 0.002,…, 0.009}. We simulated 1, 000 replicates for each value of *N* and *µ*, generating a total of 70, 000 complete species trees. We then analyzed each tree using `tMEDUSA` and `nMEDUSA`, and, as in previous simulations, repeated the entire series of analyses evaluating ‘stem’ and ‘crown’ diversification-rate shifts (we chose to omit `oMEDUSA` from these analyses because it proved to be prohibitively slow for completely sampled trees of these sizes).

Results for simulations based on completely sampled trees are summarized in Figure 4 (see also Table S.7). The overall Type I error rates were substantially lower in these simulations: `tMEDUSA` and `nMEDUSA` exhibited Type I error rates of 4% and 26%, respectively. As in the previous simulations, the disparity in the behavior of the two newer `MEDUSA` algorithms is unexpected; we will return to this issue in the Discussion section. Importantly, the marked decrease in the overall Type I error rate for this set of simulations—which control for the effect of unresolved terminal clades— provides further evidence that the pathological behavior of the `MEDUSA` methods stems either (1) from a problem computing the likelihood for terminal unresolved clades, and/or (2) from a problem combining the likelihoods for the terminal unresolved clades and the backbone tree. We explore the former possibility immediately below, and return to the latter possibility in the Discussion.

#### Assessing the impact of terminal unresolved lineages

Could errors in calculating the likelihood of terminal unresolved clades be adversely impacting the performance of the `MEDUSA` algorithms? We explored this possibility by simulating trees with *N* = {1, 2,…, 10, 20,…, 100, 200,…, 1000} species under a constant-rate birth-death process where the speciation rate was fixed to *λ* = 0.01 and the extinction rate was incremented over a range of values, *µ* = {0.000, 0.001, 0.002,…, 0.009}. For each value of *µ* and *N*, we simulated 1,000 trees and recorded the stem age of each tree, *t*. We then used the stem age, *t*, and the size, *N*, of each tree—effectively treating it as a unresolved terminal lineage—to obtain maximum-likelihood estimates of *λ* and *µ* (using equation 2, above, from Rabosky et al. 2007).

Results for simulations based on terminally unresolved lineages reveal that maximum-likelihood estimates of net-diversification rates and relative-extinction rates are strongly biased for very small lineages (Figure S.7, Tables S.8 and S.9). Not surprisingly, estimates become extremely biased when the terminal lineage includes a single species. Regardless of the true rates used to simulate these ‘singleton’ lineages (*N* = 1), the estimated net-diversification and relative-extinction rates must be zero, as there are no observed speciation (or extinction) events over the duration of the lineage, *t*. To assess whether these biased parameter estimates could impact Type I error rate, we explored the associated likelihood scores.

For each simulated tree, we computed the difference between the estimated log-likelihood of the data (ln *𝓛*_est_, based on the estimated parameter values, 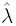 and 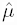), and the actual log-likelihood of the data (ln *𝓛*_true_, based on the generating parameter values, *λ* and *µ*):

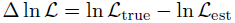

More negative values of ∆ ln *𝓛* correspond to stronger support for a model that differs from the true model (note that ∆ ln *𝓛* can never be greater than zero). Echoing the bias in parameter estimates, singleton lineages exhibit strongly biased log-likelihood estimates. On average, the ∆ ln *𝓛* values for singleton lineages were nearly twice those of other lineages 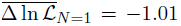 compared to 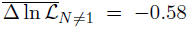; Figure 5). This ‘singleton effect’ stems from instability in the likelihood function: for a tree with a single extant species, the maximum-likelihood estimate of the net-diversification rate converges to zero, and the likelihood of the data becomes 1.

To assess whether this singleton effect contributes to the detection of spurious diversification-rate shifts, we explored the relationship between presence of singletons and patterns of Type I error in our simulated trees. For the initial constant-rate simulation, trees that were (incorrectly) identified by `tMEDUSA` and `nMEDUSA` to include a diversification-rate shift were significantly enriched for singletons (*χ*^2^-test *p* = 0.012 and *p* ≤ 10^−16^ for `tMEDUSA` and `nMEDUSA`, respectively); by contrast, `oMEDUSA` was significantly more likely to identify diversification-rate shifts in trees without singletons (*p* ≤ 10^−16^). The relationship between tree size, resolution, relative extinction, and the enrichment of singletons among false-positive cases is complex (Figure S.8, Table S.10), but we note two general patterns: 1) in large regions of parameter space—especially when *N* = 10, 000 and *k* is small—`tMEDUSA` and `nMEDUSA` *always* identify a spurious diversification-rate shift in the presence of a singleton, and; 2) the singleton effect decreases as phylogenetic resolution increases.

### Parameter Estimation

The ability of `MEDUSA` to estimate parameters of the birth-death stochastic-branching process represents an important advantage over competing methods. This allows us to not only identify the number and phylogenetic distribution of diversification-rate shifts, but also to estimate the magnitude of corresponding changes in the net-diversification rate and/or relative-extinction rate. Although the intended focus of `MEDUSA` is estimating the location and number of diversification-rate shifts, it may also be used with a focus on estimating (absolute or relative) diversification rates. Most methods for estimating diversification rates assume that rates are constant through time and/or across lineages. However, estimates based on these methods are apt to be biased if the assumption of rate homogeneity is violated by shifts in diversification rate across lineages. In principle, `MEDUSA` can provide more reliable diversification-rate estimates by accommodating diversification-rate shifts as nuisance parameters. Accordingly, even if—as we have demonstrated above—`MEDUSA` does not provide reliable estimates of the location and number of diversification-rate shifts, it may nevertheless prove useful for estimating parameters of the birth-death process. In this section, we explore the ability of `MEDUSA` to provide reliable estimates of diversification-rate parameters.

#### Reliability of parameter estimates when the correct diversification model is selected

How reliable are rate parameter estimates when `MEDUSA` selects the correct (constant-rate) diversification model? To explore this question, we examined parameter estimates for the cases in which `MEDUSA` identified the correct diversification model for trees simulated under a constant-rate birth-death process. On average, net-diversification rates estimated by `oMEDUSA`, `tMEDUSA`, and `nMEDUSA` were 479.7%, 113.0%, and 106.5% their true values, respectively (Figures 6, S.21). Estimates of relative-extinction rate were similarly biased: parameter values inferred by `oMEDUSA`, `tMEDUSA`, and `nMEDUSA` were on average 6.7%, 20.2%, and 27.8% their true values, respectively (Figures S.9, S.22). In summary, even when `MEDUSA` selects the correct diversification model, parameter estimates are severely biased.

#### Reliability of estimated magnitude of diversification-rate shifts

Imagine that we have detected a significant diversification-rate shift using `MEDUSA`. Given the extremely high Type I error rate, we should be concerned that this diversification-rate shift is actually spurious. Imagine further, however, that the magnitude of the inferred diversification-rate shift is estimated to be very large. We might hope that the estimated magnitude could be used to help us distinguish between spurious and *bona fide* diversification-rate shifts. This would require that—despite severe bias in estimates of the *absolute* net-diversification and relative-extinction rates—estimates of the *relative* difference between these biased rate estimates prove to be reliable.

To explore this possibility, we plotted the magnitude of spurious diversification-rate shifts for the cases where `MEDUSA` incorrectly identified a multi-rate model in the constant-rate simulations. Spurious shifts in net-diversification rate were inferred to involve large average magnitudes (and broad 95% quantiles): 2.3 (4.7), 2.7 (6.7), and 2.4 (4.8) for the `oMEDUSA`, `tMEDUSA`, and `nMEDUSA` algorithms, respectively (Figures 7, S.25, S.12, S.26). Similarly, spurious shifts in relative-extinction rate were inferred to have large average magnitudes (and wide 95% quantiles): 2.7 (5.7), 3.1 (7.7), and 3.4 (7.9) for `oMEDUSA`, `tMEDUSA`, and `nMEDUSA`, respectively (Figures S.13, S.27, S.14, S.28). Accordingly, the magnitude of diversification-rate shifts estimated using `MEDUSA` are unreliable, and so cannot be used to distinguish ‘real’ from ‘spurious’ results when the method is applied to empirical data.

## Discussion

Biologists are clearly interested in identifying shifts in diversification rate along lineages, and have enthusiastically embraced `MEDUSA` as an approach for addressing this problem (Table S.1). `MEDUSA` has been used more frequently in empirical studies than any other statistical phylogenetic method for identifying lineage-specific diversification rates (Figure S.3), and, in fact, has been used more frequently than *any* statistical phylogenetic method for studying lineage diversification. Nevertheless, our simulation study reveals that `MEDUSA` is deeply flawed. In this section, we first summarize our inferences regarding the statistical behavior of `MEDUSA`, then consider the various pathologies that render this method unreliable, and conclude with a discussion of the more general implications of our findings.

### Characterizing the statistical (mis)behavior of MEDUSA

For almost half of the trees that we simulated in our study—where rates were strictly constant across lineages of the tree—`MEDUSA` identified strong support for one or more diversification-rate shifts. This deeply troubling result holds for all versions of the algorithm (`oMEDUSA`, `tMEDUSA`, and `nMEDUSA`), applies to estimates derived using available options for these methods (whether rate shifts are assumed to occur along ‘stem’ or ‘crown’ nodes; see Supplemental Material), and holds over a wide range of absolute parameter values (Figures 2, S.5, S.6, S.17, S.18) that broadly encompass the conditions encountered in the analyses of actual empirical datasets (Figures S.1, S.2; Table S.1). The selection of (incorrect) multi-rate models by `MEDUSA` does not represent a slight bias. Rather, the (correct) constant-rate model is decisively rejected. Recall that the conventional threshold indicating a significant difference between two competing models is ∆AIC_crit_ = 4. By contrast, the computed ∆AIC values in favor of the incorrect diversification model exceeded the critical threshold of significance by an average of 522.7 AIC units (the `oMEDUSA`, `tMEDUSA`, and `nMEDUSA` algorithms had average values of 1035.5, 3.5, and 6.4, respectively; Figures 3, S.19).

We emphasize that these extreme false discovery rates are not artifacts of our simulation study. Our findings cannot be attributed to the optimization routine being trapped on local optima far away from global optima, as we initiated analyses of simulated trees from the true parameter values (those used to simulate the data). Our characterization of the statistical behavior of `MEDUSA` is therefore somewhat optimistic; the method will fare worse when applied to real datasets, where the true parameter values are unknown. Additionally, our conclusions are not manifestations of Monte Carlo error: we simulated more than twice the number of replicates required for precise estimates of the level and patterns of Type I error (Figures S.4, S.16, Tables S.3, S.20).

`MEDUSA` also provides severely biased estimates of diversification-rate parameters (Figures 6, 7). Even when `MEDUSA` correctly identified the (true) constant-rate model, the net-diversification and relative-extinction rate estimates were on average 183% and 20% of their true values, respectively (Figures 6, S.9, S.21, S.22). Similarly, when `MEDUSA` identified the (incorrect) multi-rate model, the spurious diversification-rate shifts were inferred to involve rate changes of large magnitude. On average, spurious diversification-rate shifts were estimated to involve a 2.5-fold change in net-diversification rate (Figures 7, S.25), with a large spread of values (the upper 95% quantile was an average 5.5-fold shift in net-diversification rate; Figures S.12, S.26). Similarly, these illusory diversification-rate shifts were estimated to entail an average 3.1-fold change in relative-extinction rate (Figures S.13, S.27), with a correspondingly large spread of values (the upper 95% quantile was an average 7.4-fold shift in net-diversification rate; Figures S.14, S.28).

In summary, many of the putative advances that make `MEDUSA` appealing seem to be problematic. For example, the ability to accommodate incompletely sampled trees with `MEDUSA` is an attractive feature of the method; in fact, most applications (82.5%) of this method involve incomplete trees. Nevertheless, the pathological behavior of `MEDUSA` is most extreme under these conditions (Figures 4, S.20). Likewise, the adoption of a birth-death branching process is presented as an important advantage of `MEDUSA` over competing methods that do not accommodate extinction. However, (relative) extinction-rate estimates obtained using `MEDUSA` are effectively meaningless (Figures S.9, S.11, S.22, S.24). This finding is relevant to the general debate regarding whether (relative) extinction rates can (*e.g.*, Rabosky et al. 2007; Rabosky 2014) or cannot (*e.g.*, Rabosky 2010) be reliably estimated from phylogenies. Our sense is that procuring reliable estimates of (relative) extinction rates from phylogenies of extant species is generally difficult when diversification rates are constant, and becomes extremely difficult when diversification rates experience tree-wide changes through time, and is effectively impossible (or at least inadvisable) when diversification rates change along lineages.

### Diagnosing the pathologies afflicting MEDUSA

The pathological behavior of `MEDUSA` stems from two primary causes: the likelihood functions are incorrect, and the thresholds for evaluating the fit of candidate diversification models are unreliable. The likelihood function is critical to likelihood-based (maximum-likelihood and Bayesian inference) methods, as it is the vehicle that conveys information in the data to estimate the model parameters. Accordingly, even a minor error in the likelihood function of a method is likely to render it unreliable. The likelihood functions of `MEDUSA` contain two major flaws. First, the likelihood function used to compute the probability of terminal lineages (equation 2) is unstable. As we have demonstrated, this instability causes severely biased estimates of parameters (Figure S.7) and the corresponding log-likelihoods (Figure 5) for ‘singleton’ terminal lineages (*i.e.*, those comprising a single species). Consequently, when `MEDUSA` is applied to a tree containing one or more terminal lineages with a single species, this ‘singleton effect’ creates the illusion of rate variation within the tree, which may cause overfitting of the diversification models. That is, the singleton effect contributes to the selection of diversification models that entail one or more spurious rate shifts.

Second, and more profoundly, `MEDUSA` is based on a composite likelihood function (equation 3; Rabosky et al. 2007) that is invalid for trees in which diversification rates change across lineages. The expression for the composite likelihood (*i.e.*, for the backbone, *𝓘*, and the terminal unresolved lineages, *𝒯*, of the tree) is only correct when rates of diversification are stochastically constant. The probabilities for the internal branches, *P* (*τ*_*i*_ | ⋅), and terminal unresolved polytomies, *P* (*n*_*i*_ | ⋅), are both conditional on a homogeneous rate of speciation and extinction, *λ*_*i*_, *µ*_*i*_, across the tree. Variation in diversification rates across branches and/or terminal lineages—the process that `MEDUSA` was designed to detect—clearly violates this condition and results in biased parameter estimates and incorrect likelihoods.

The inability of `MEDUSA` to correctly estimate the likelihood of a given diversification model is a fatal problem. However, `MEDUSA` is further confounded by problems associated with selecting among candidate diversification models. Recall that `MEDUSA` uses the AIC to select among competing diversification models: we first estimate the maximum-likelihood score, ln *𝓛*, for two adjacent diversification models—*M*_*i*_ and *M*_*i*+1_ with *i* and *i* + 1 diversification-rate categories—then calculate the AIC score for each candidate model—AIC = 2*p* − 2 ln *𝓛* with *p* free parameters—and finally compute the difference in the AIC scores for the two competing models—∆AIC = AIC_*i*_− AIC_*i*+1_. Finally, we compare the computed ∆AIC score to the critical threshold, ∆AIC_crit_; we reject the simpler model if the improvement in the AIC score conferred by the more complex model exceeds this threshold. The original implementation, `oMEDUSA`, used a conventional, fixed ∆AIC_crit_ threshold. As we have shown (Figures 3, S.19), a fixed critical ∆AIC_crit_ threshold is not a viable solution: the surface of computed ∆AIC values varies considerably depending upon the tree size, *N*, the number of terminal unresolved lineages, *k*, and the relative-extinction rate, *∈*. When the computed ∆AIC values exceed the fixed ∆AIC_crit_ threshold, the `MEDUSA` will select the incorrect (overly complex) diversification model, and in so doing, identify spurious diversification-rate shifts.

Developers of `MEDUSA` also appear to have realized that a conventional, fixed ∆AIC_crit_ threshold is not appropriate for selecting among diversification models (Luke Harmon, personal communication). As we understand it, they developed the following solution to this problem. First, they performed a simulation study that was similar in some respects to the current study: they simulated trees of various sizes, *N*, under a constant-rate birth-death process. Next, they generated the distribution of ∆AIC values computed for one-versus two-rate diversification models for the various tree sizes. Finally, they devised a function that approximated the resulting distribution of computed ∆AIC values, which they use to specify the ∆AIC_crit_ threshold values in the newer versions of the algorithm. This is the basis for the surface of AIC_crit_ values depicted in Figures 3 and S.19.

Although clearly an improvement over the conventional, fixed ∆AIC_crit_ critical threshold, this solution is still inadequate. The critical ∆AIC_crit_ threshold used in the two newer algorithms, `tMEDUSA` and `nMEDUSA`, is strictly a static function of tree size, *N*, for completely sampled trees. As we have seen, however, computed ∆AIC values are also strongly impacted by the degree of phylogenetic resolution (*i.e.*, the number of terminal unresolved linages, *k*), the relative extinction rate, *∈*, and the absolute diversification rate. Importantly, the static function implemented in `tMEDUSA` and `nMEDUSA` does not capture the effect of these factors on the computed ∆AIC values. Accordingly, when the data depart from the precise conditions used to devise the static function, the critical ∆AIC_crit_ threshold based on that static threshold becomes increasingly incorrect, and Type I error rate becomes increasingly inflated.

This insight suggests a possible solution: we simply need to devise a more complex static function that captures the impact of these additional factors on the computed ∆AIC values. However, we believe that such an extension will not provide a viable solution. The static function used to describe the critical ∆AIC_crit_ threshold in `tMEDUSA` and `nMEDUSA` is based on constant-rate simulations. Specifically, the function is attempting to describe the distribution of ∆AIC values computed for the (correct) constant-rate versus the (incorrect) two-rate models, ∆AIC = AIC_one-rate_ − AIC_two-rate_. Our current study focusses exclusively on simulations under a constant-rate birth-death process. However, our work on a related project involves an extensive simulation study of birth-death processes that involve one or more diversification-rate shifts (May, Höhna, and Moore, in preparation). We have discovered that the computed ∆AIC values depend on the absolute degree of the models being compared. That is, all else being equal, a critical ∆AIC_crit_ threshold that is correct for selecting between diversification models with one-versus two-rate categories is not reliable for selecting between diversification models with two-versus three-rate categories.

Accordingly, providing a correct and general solution for robust selection of diversification models may require that we dynamically generate critical ∆AIC_crit_ thresholds that are precisely tailored for the specific comparison at hand. We are optimistic that such a solution could be devised using Monte Carlo simulation; this is an area of current work (May, Höhna, and Moore, in preparation).

### General implications of the current study

The problematic likelihood function used in `MEDUSA` (equation 3; Rabosky et al. 2007) is also used in—and may similarly compromise the reliability of—several other methods developed for estimating diversification rates from incompletely sampled phylogenies, including those in LASER (Rabosky et al. 2007) and MECCA (Slater et al. 2012). Similarly, many statistical phylogenetic methods are based on selecting among candidate models using the Akaike Information Criterion. These include methods for studying lineage diversification (Rabosky 2006; Rabosky and Lovette 2008; Rabosky and Glor 2010), and approaches for exploring the evolution of continuous traits under either Brownian motion (Thomas and Freckleton 2012) or Ornstein-Uhlenbeck (Ingram and Mahler 2013) process models. The AIC is also used to assess the relative fit of candidate models of nucleotide substitution (*e.g.*, Posada 2008; Darriba et al. 2012), to select among partition schemes (*e.g.*, Lanfear et al. 2012), and to choose among models of lineage-specific natural-selection pressure (*e.g*, Pond and Frost 2005). The AIC provides a seemingly attractive solution for choosing among models—it is fast and easy to compute, and can be flexibly applied to select among nested or non-nested models. However, our results accord with those of previous studies (*e.g.*, Alfaro and Huelsenbeck 2006; Boettiger et al. 2012), suggesting that the AIC is not a reliable means for choosing among phylogenetic models.

Owing to the inherent complexity of the inference problems, the vast majority of comparative phylogenetic methods cannot be solved analytically, and so resort to numerical approximations (*i.e.*, hill-climbing algorithms used for maximum-likelihood estimates of parameters or MCMC algorithms used in Bayesian methods to approximate the joint posterior probability density of parameters). Careful simulation is therefore required to validate these methods: this approach can uniquely assess the ability of the inference methods to recover the true (known) parameters and/or model used to generate the data. This is particularly true of *ad hoc* methods (*i.e.*, those not developed within a formal modeling framework), which are becoming increasingly popular in the field of comparative phylogenetics^1^.

Accordingly, the design and execution of simulation studies deserve our careful consideration. A simulation study will provide meaningful insights into the statistical behavior of a given method only if it adequately explores the relevant parameter space. Conceptually, this entails circumscribing an *N*-dimensional hypervolume (for methods impacted by *N* variables, such as tree size, diversification rate, etc.) that is likely to encompass the conditions encountered in the course of applying the method to real data. This hypervolume is discretized into cells, each representing a unique combination or parameter values. The granularity of the discretization required to faithfully characterize the statistical behavior of a method depends on its details. Methods developed in a formal modeling framework generally require a less granular discretization of parameter space, as their direct connection to established statistical theory provides a basis for interpolating their behavior between sampled points in parameter space. By contrast, *ad hoc* methods generally require a more finely discretized (and therefore more demanding) exploration of parameter space, as their behavior is apt to be relatively unpredictable. This behavioral inscrutability is manifest in the present study: *e.g.*, Type I error rates and bias in parameter estimates varied wildly over parameter space (Figures 2, 3). Finally, each discrete cell of parameter space must be explored with sufficient intensity. Practically, this means simulating an adequate number of outcomes for each unique combination of parameter values to provide ‘acceptable’ error variance on the corresponding estimates. The degree of Monte Carlo error associated with a finite number of replicates can (and should) be assessed (*c.f.*, Figure S.4). Careful simulation studies represent a non-trivial enterprise: our study of the `MEDUSA` algorithm involved ∼ 7, 000, 000 analyses that consumed ∼ 35, 000 hours (∼ 4 years) of CPU time.

From this perspective, our experience emphasizes that the increasingly popular use of `R` for developing comparative phylogenetic methods is a double-edged sword. The primary advantage is the very low threshold required to quickly implement a method, often by drawing upon existing `R` functions and packages. The downside, however, is that the extremely slow speed of methods implemented purely in `R` (which is typically ∼ 30 − 60 times slower than low-level programming languages, such as `C` and `C++`) is a major impediment to the necessary validation of these methods via simulation. For this reason, the statistical behavior of many recent comparative phylogenetic methods is largely (or completely) unknown. We believe this is a reason for serious concern.

The primary conclusion of our simulation study—that the `MEDUSA` methods are deeply flawed— might be viewed as a ‘negative’ result. After all, our findings cast considerable doubt on the conclusions of the many empirical studies that have used `MEDUSA`, and argue strongly against the application of this approach in future empirical studies. Nevertheless, we believe that it is far better to be aware of the limitations of a method (even to the extent where it is deemed unusable) than to (unwittingly) continue using a method that is likely to provide spurious results. Moreover, beyond demonstrating that `MEDUSA` is unreliable, the results of our simulation study provide insights into the reasons *why* these methods are unreliable. Understanding the nature of these problems can focus theoretical efforts to develop more reliable methods. Accordingly, far from discouraging, we view the findings of this study as cause for optimism. We are hopeful that future efforts will resolve issues afflicting the `MEDUSA` framework for identifying shifts in diversification rates that—coupled with rigorous evaluation of these new methods—will continue to enhance our ability to explore a broad range of fundamental evolutionary processes.

**Figure 1.**
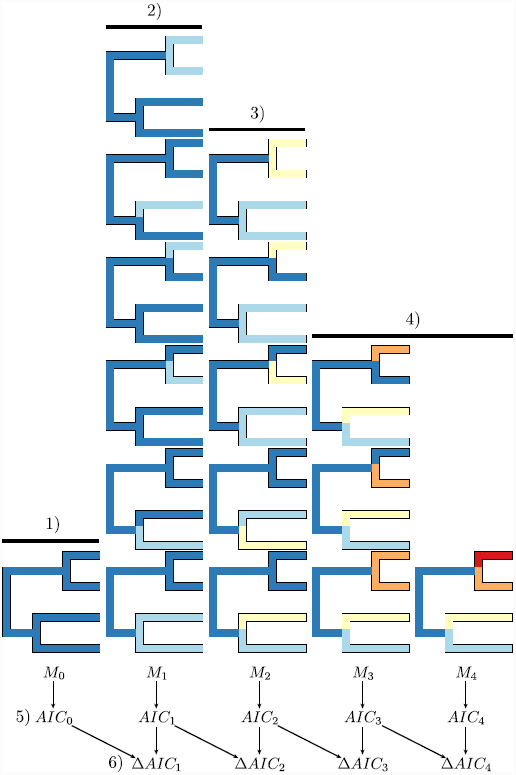
The MEDUSA algorithm. 1) The one-rate model, *M*_1_, is fit to the data using maximum likelihood. 2) Every possible two-rate model is then fit to the data, and the models are ranked by their likelihood score (with the best model, *M*_2_, at the bottom). 3) Every possible three-rate model that is consistent with the best two-rate model is then fit to the data, and the models are then ranked as in the previous step. 4) Step 3 is repeated for increasingly complex models until the most complex model has been evaluated. 5) The AIC score is computed for the best model in each index (*i.e.*, with 0, 1,…rate shifts). 6) The ∆AIC value is calculated for each pair of adjacent models (as the difference in their AIC scores). Each of the ∆*AIC_i_* comparisons are evaluated in succession, where the more complex model is selected if the computed ∆AIC is greater than some pre-specified threshold, ∆AIC_crit_. This process continues until either the most complex model is accepted or the improvement in the AIC score is too small to exceed the ∆*AIC_crit_* threshold.

**Figure 2.**
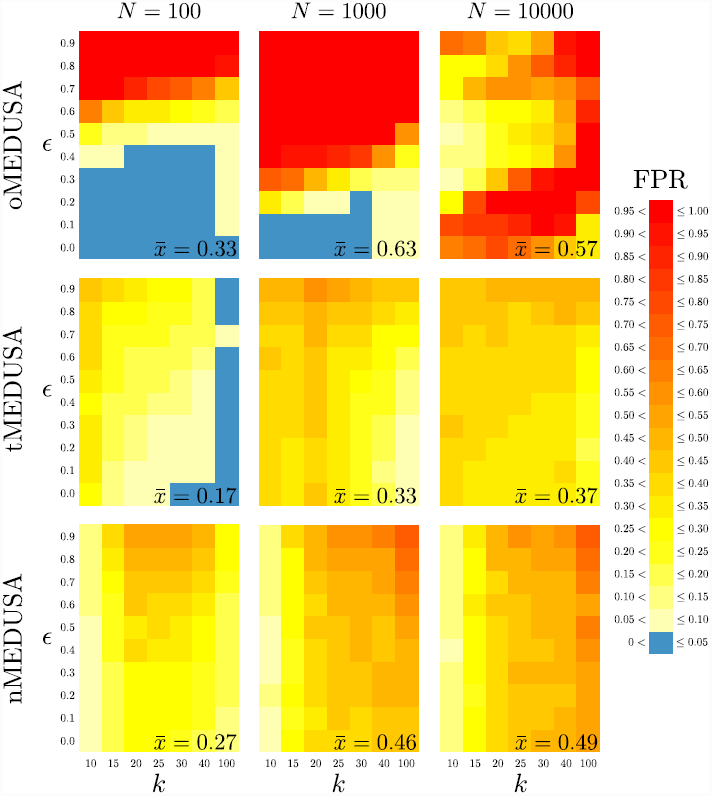
Type I error rates for the three MEDUSA algorithms. The three panels in each row summarize results for one algorithm (`oMEDUSA`, `tMEDUSA`, and `nMEDUSA`, from top to bottom), the three panels in each column summarize results for one tree size (with *N* = 100, 1, 000, 10, 000 species, from left to right). Within each panel, we plot the number of terminal lineages, *k*, against the relative-extinction rate used in the simulation. Trees were simulated under a constant-rate birth-death process using the parameters summarized in Table S.2. The cells within each panel are colored as a heat map reflecting the frequency with which spurious diversification-rate shifts were inferred (see legend). The average Type I error rate is summarized in the lower right of each panel.

**Figure 3.**
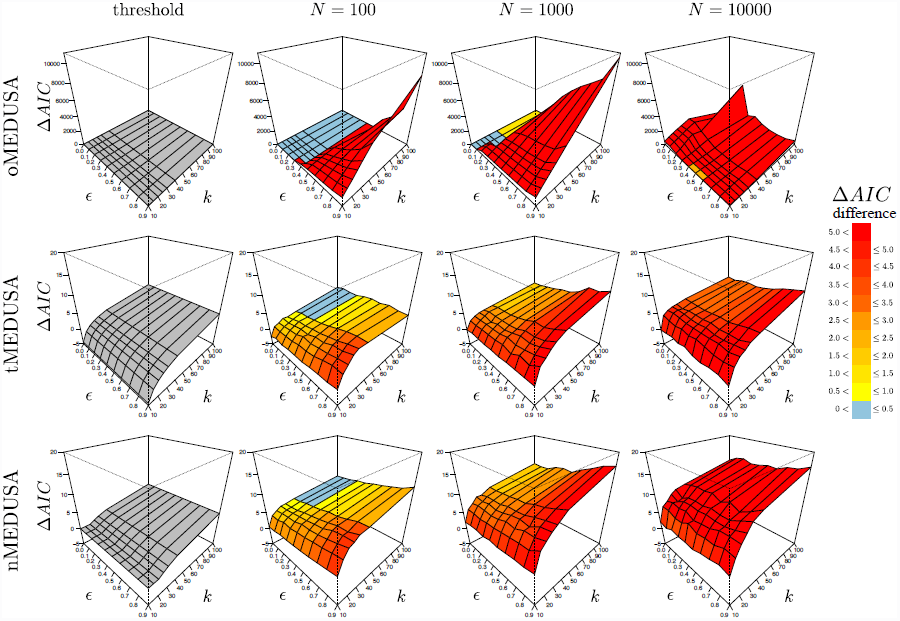
Critical and computed ∆AIC surfaces for the three MEDUSA algorithms. The four panels in each row correspond to one algorithm (`oMEDUSA`, `tMEDUSA`, and `nMEDUSA`, from top to bottom). The first column depicts the surface of ∆AIC_crit_ threshold values for each algorithm; when the difference between the AIC scores of two competing models exceeds this threshold, the more complex model is selected. Columns 2 − 4 summarize the ∆AIC_crit_ values computed from simulated trees of each size (*N* = 100, 1, 000, 10, 000). The computed ∆AIC surfaces are colored to reflect the degree to which they exceed the corresponding ∆AIC_crit_ threshold (see legend). Surfaces are computed from the trees used in the initial constant-rate simulation (*c.f.*, Figure 2) under parameters described in Table S.2. Note the difference in the scale of ∆AIC_crit_ values for the `oMEDUSA` algorithm. For detailed values, see Table S.6.

**Figure 4.**
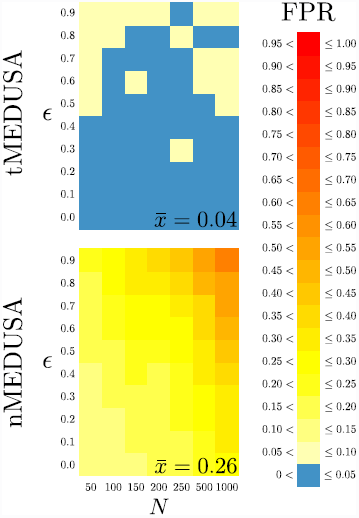
Type I error rates for completely sampled trees. Each row summarizes results for one algorithm (`tMEDUSA` and `nMEDUSA` on the top and bottom, respectively). Within each panel, we plot the number of species in the tree, *N*, against the relative-extinction rate used in the simulation. Trees were simulated under a constant-rate birth-death process using the parameters summarized in Table S.7. The cells within each panel are colored as a heat map reflecting the frequency with which spurious diversification-rate shifts were inferred (see FPR legend). The average Type I error rate is summarized in the lower right of each panel.

**Figure 5.**
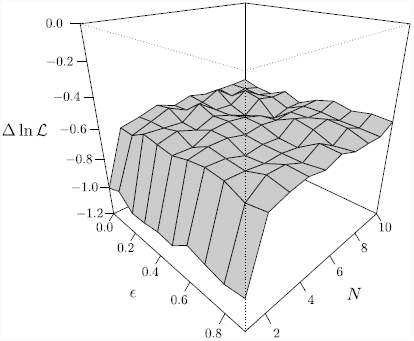
Model overfitting for terminal unresolved lineages. Trees of size *N* ranging from 1 to 1000 were simulated under a constant-rate birth-death process using parameters summarized in Table S.8. The stem age of each tree was then used to estimate parameters of a constant-rate birth-death process using equation 2. We plot the difference between the likelihood under the true parameters and the maximum-likelihood estimates of the parameters, ∆ ln *𝓛*, against the size of the tree, *N* (truncated to 10 for visibility) and the relative extinction rate, *∈*. Smaller values of ∆ ln *𝓛* indicate a stronger preference for the maximum-likelihood parameters against the true parameters.

**Figure 6.**
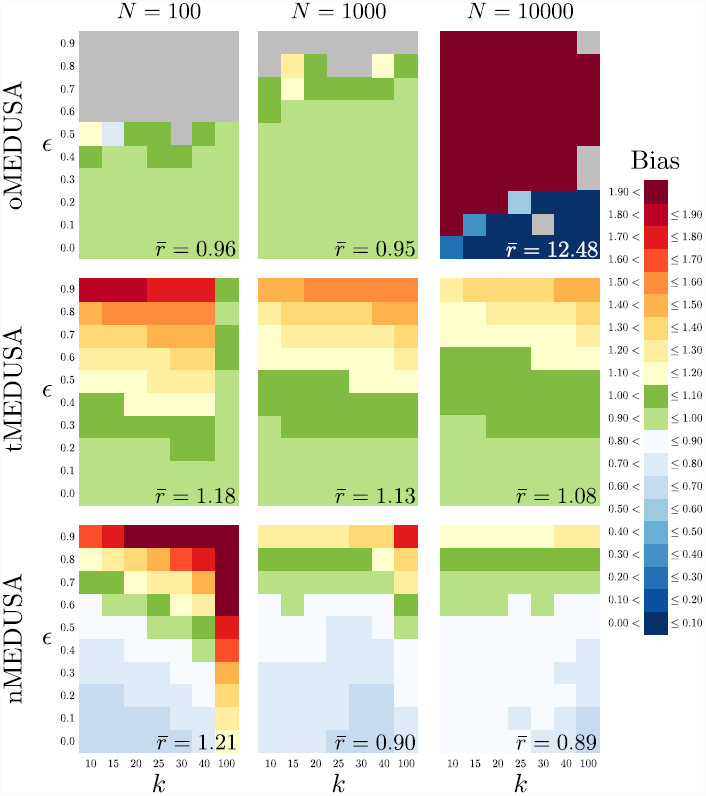
Bias in net-diversification rate estimates when the constant-rate model is selected. The three panels in each row summarize results for one algorithm (`oMEDUSA`, `tMEDUSA`, and `nMEDUSA`, from top to bottom), the three panels in each column summarize results for one tree size (with *N* = 100, 1, 000, 10, 000 species, from left to right). Within each panel, we plot the number of terminal lineages, *k*, against the relative-extinction rate used in the simulation. Trees were simulated under a constant-rate birth-death process using the parameters summarized in Table S.11. The cells within each panel are colored as a heat map reflecting the average difference between the estimated and true net-diversification rates when the one-rate model is correctly identified (see Bias legend). The average bias in the inferred net-diversification rate is summarized in the lower right of each panel.

**Figure 7.**
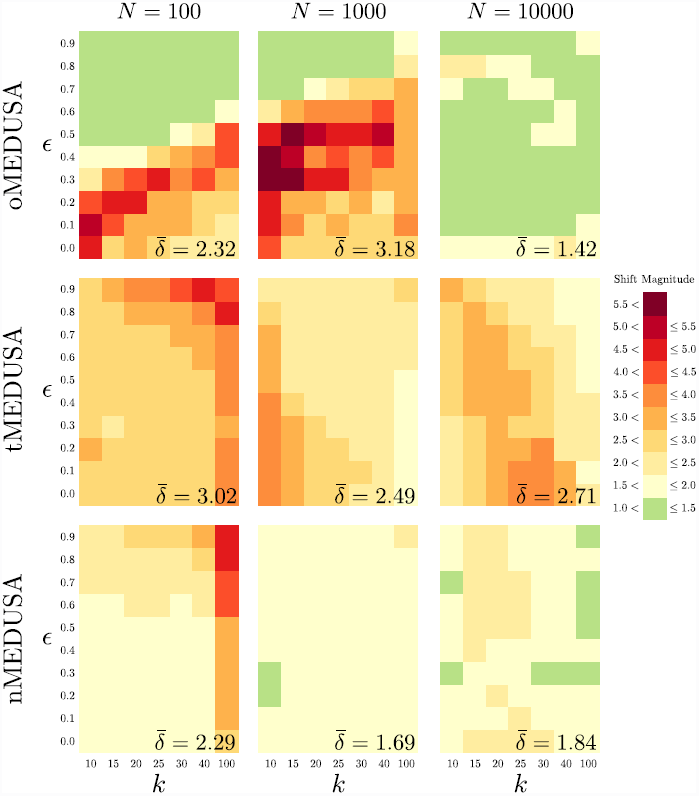
Magnitude of shifts in net-diversification rate when a two-rate model is chosen. The three panels in each row summarize results for one algorithm (`oMEDUSA`, `tMEDUSA`, and `nMEDUSA`, from top to bottom), the three panels in each column summarize results for one tree size (with *N* = 100, 1, 000, 10, 000 species, from left to right). Within each panel, we plot the number of terminal lineages, *k*, against the relative-extinction rate used in the simulation. Trees were simulated under a constant-rate birth-death process using the parameters summarized in Table S.15. The cells within each panel are colored as a heat map reflecting the average magnitude of shifts in net-diversification rate when a two-rate model is incorrectly identified (see Shift Magnitude legend). In order to accommodate the bimodal distribution of rate shifts, the magnitude of each net-diversification-rate shift was computed as the ratio of the higher rate to the lower rate, regardless of which rate corresponded to the background-rate category. The average magnitude of inferred diversification-rate shifts is summarized in the lower right of each panel.

## Acknowledgments

We are grateful to Michael Turelli and Peter Wainwright for encouraging us to pursue this project, to Yaniv Brandvain, Tracy Heath, Sebastian Höhna, John Huelsenbeck, Michael Landis, Bruce Rannala, and Tanja Stadler for thoughtful discussion, and to Andrew Magee for assistance with the survey of empirical studies. We are particularly grateful to Mike Alfaro and Luke Harmon for sharing their insights on this work. This research was supported by NSF grants DEB-0842181, DEB-0919529, and DBI-1356737 awarded to BRM. Computational resources for this work were provided by an NSF XSEDE grant (DEB-120031) to BRM.

## Footnotes

1 `MEDUSA` is classified as an *ad hoc* method because it does not explicitly model changes in diversification rate over the tree as a stochastic process, but instead heuristically evaluates the assignment of diversification rates to branches of the tree according to an arbitrarily algorithm.

## Supplementary Material

### S.1 A More Detailed Description of the MEDUSA Algorithm

There are (2*k* − 2) clades in the tree, *k* of which are terminal branches and (*k* − 2) of which are nodes. Label the terminal branches in the tree 1 through *k*, and all the non-root nodes in the tree (*k* + 1) through (2*k* − 2) in arbitrary order.

1. Fit a single-rate diversification model to Ψ by maximum likelihood using equation (3); compute *AIC*_one-rate_.
2. Set *i* = 2. This is an index of the number of rate classes under consideration.
3. Set *j* = 1. This is an index of the clade under consideration for a rate shift.
4. Visit clade *j*.
  a. If clade *j* already has a rate shift, go to step 5.
  b. Define a partition of branches into categories where the descendant branches of clade *j* are in one rate category, all the descendant branches of any other rate shifts are in their own rate categories, and the rest of the branches are in a “background” rate category. When a branch might belong to multiple rate categories, the branch is placed in the category defined by the youngest clade.
  c. Fit a multiple-rate diversification model defined by the partition to Ψ by maximum likelihood using equation (3); compute 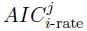.
5. If *j* = (2*k* − 2), go to step 8.
6. Set *j* = *j* + 1.
7. Return to step 4.
8. Determine the best (lowest) 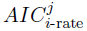. Call it *AIC*_*i*-rate_.
9. Compute ∆*AIC* = *AIC*_(*i*−1)-rate_ − *AIC*_*i*-rate_.
10. If ∆*AIC* ≥ ∆*AIC*_crit_, accept the shift at the clade with the best 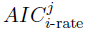. Otherwise, go to step 14.
11. If *i* = (2*k* − 2), go to step 14.
12. Set *i* = *i* + 1.
13. Return to step 3.
14. Terminate the algorithm.

### S.2 Survey of empirical studies using MEDUSA

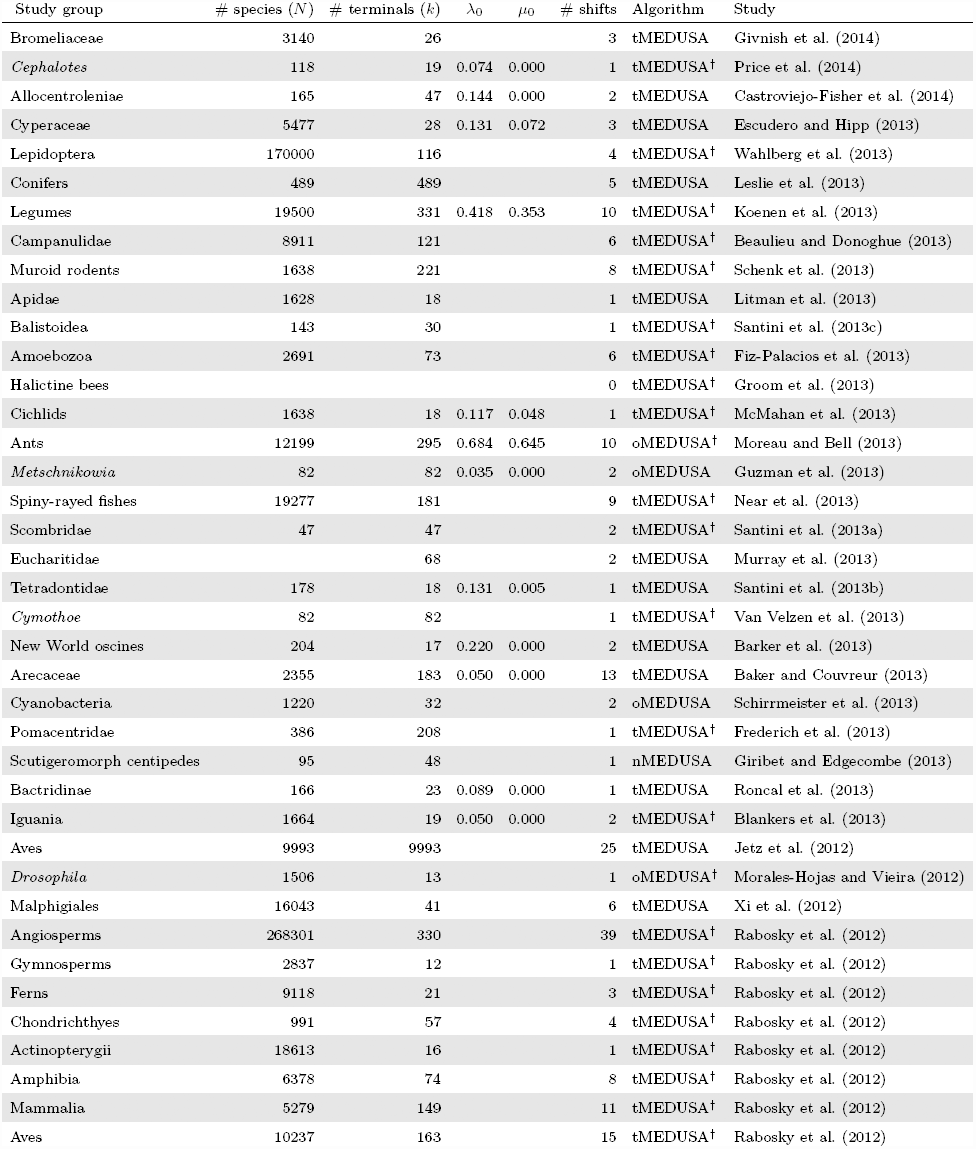
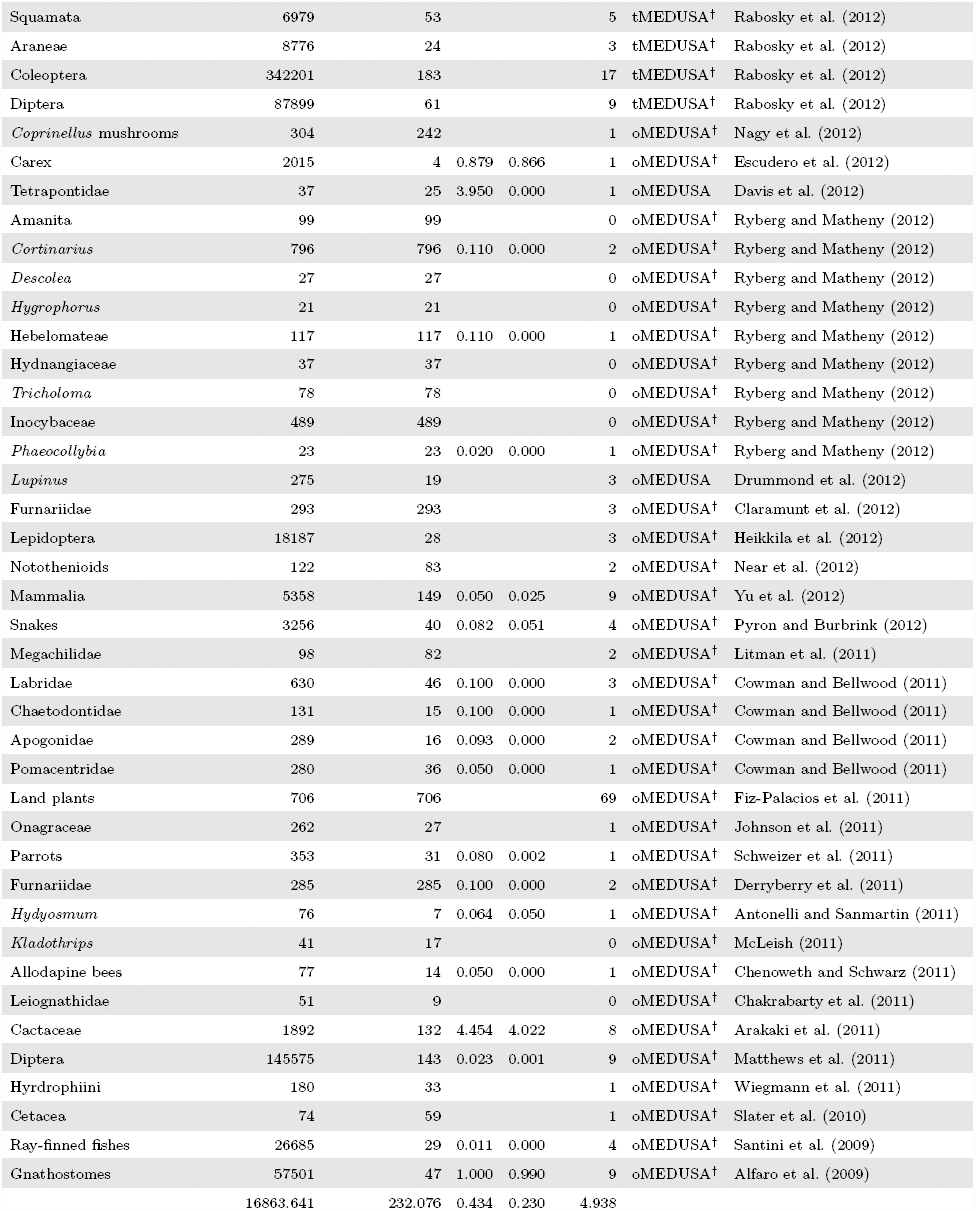
N.B. The total number of species (*N*), the number of incompletely resolved terminal lineages (*k*), the estimated background speciation (*λ*_0_) and extinction (*µ*_0_) rates in species per species per million years, and the inferred number of diversification-rate shifts (Shifts) inferred by published studies using various `MEDUSA` algorithms. ^†^: These studies did not provide sufficient information to unambiguously determine the particular algorithm used; in these cases, we assigned `oMEDUSA` if the study was published before the release of `GEIGER 1.99` and `tMEDUSA` if it was published after.

**Figure S.1:**
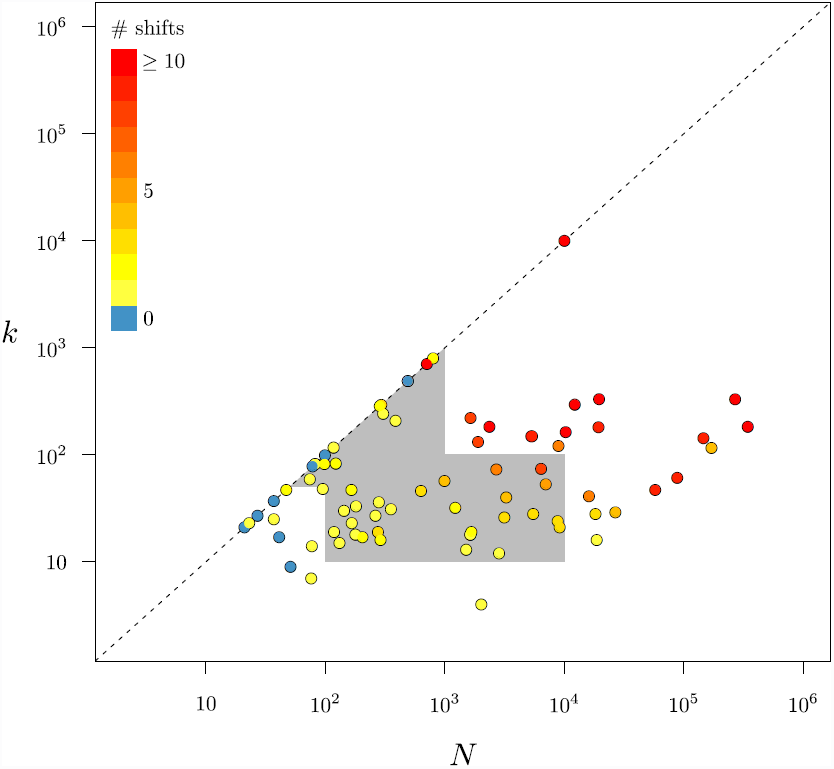
Parameter space of empirical datasets that have been analyzed with MEDUSA: tree size and phylogenetic resolution. A log-log plot of the number of species in each dataset, *N*, against the number of terminal lineages in each tree, *k*. Each dataset is colored by the number of shifts inferred by `MEDUSA`. Datasets with complete species sampling fall on the dashed line, while those with incomplete sampling fall below the dashed line. The grey area contains the values of *N* and *k* used in our simulation study.

**Figure S.2:**
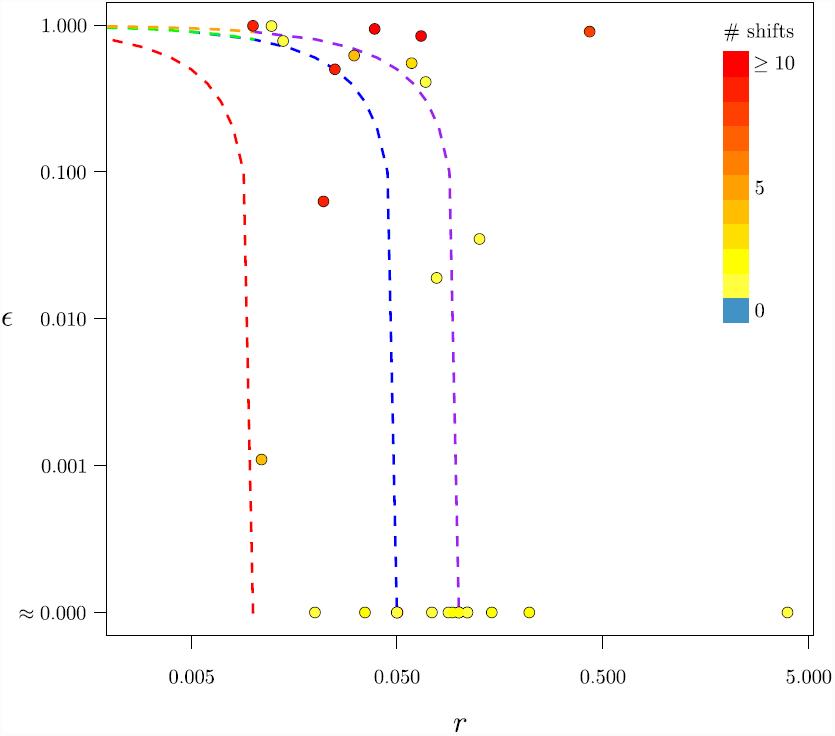
Parameter space of empirical datasets that have been analyzed with MEDUSA: diversification rates. A log-log plot of the net-diversification rate, *r*, against the relative-extinction rate, *∈*, for the background rate category of empirical datasets analyzed with `MEDUSA`. Each dataset is colored by the number of shifts inferred by `MEDUSA`. Dashed lines correspond to parameter settings used in our simulation study. Red: Table S.2; Blue: Table S.4, left; Purple: Table S.4, right; Green: Table S.5, left; Orange: Table S.5, right.

**Figure S.3:**
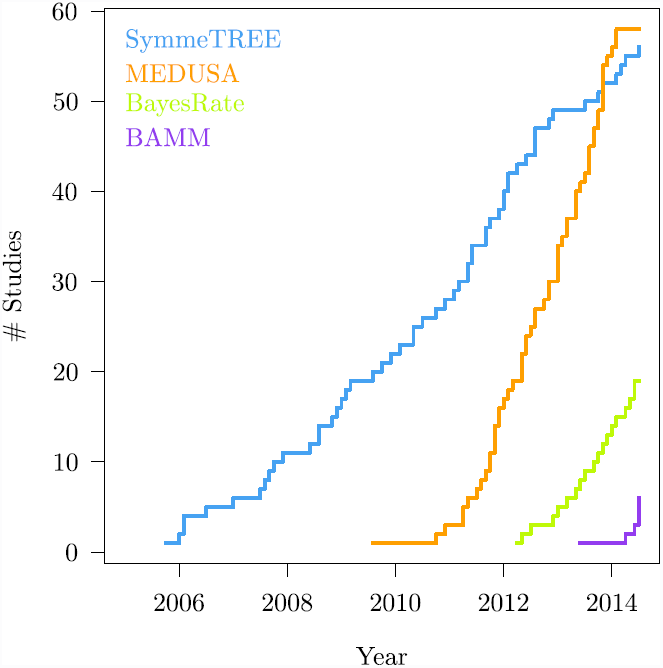
Usage of alternative methods for detecting lineage-specific diversification-rates in empirical studies.

### S.3 The effect of relative-extinction rate, tree size, and resolution (stem shifts)

**Table S.2:**
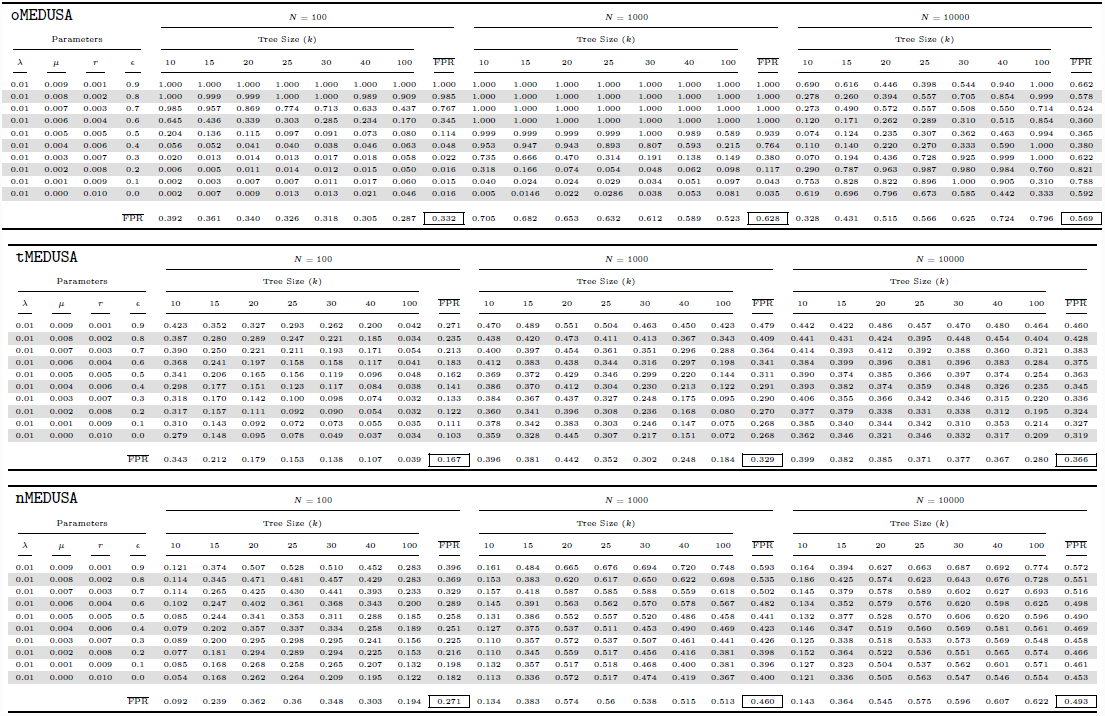
Parameters and estimated Type 1 error rates for the initial constant-rate simulations (stem shifts)

### S.4 Exploring the impact of Monte Carlo error on Type I error rates (stem shifts)

**Figure S.4:**
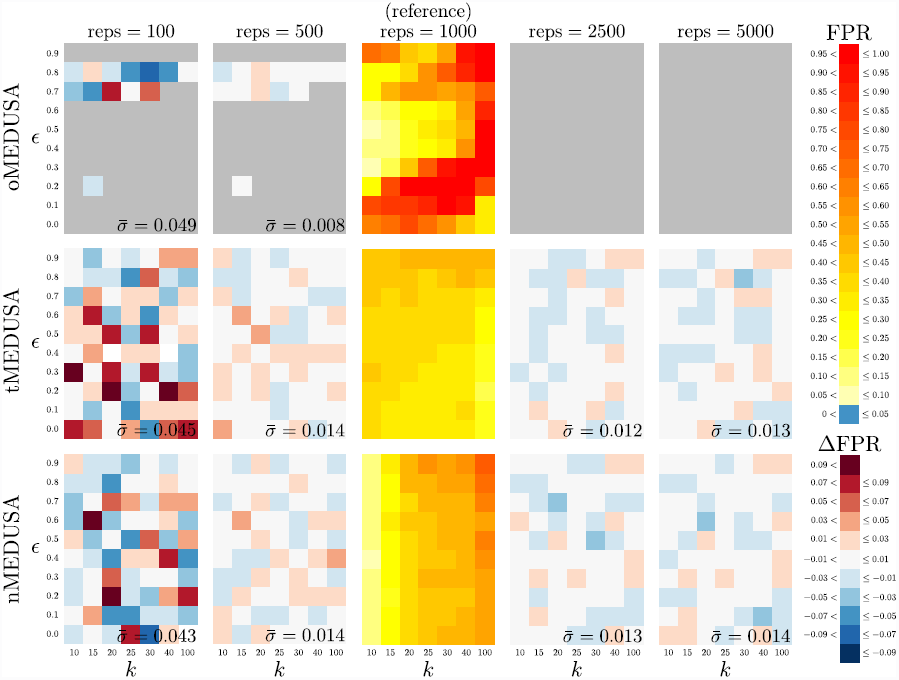
Exploring the impact of Monte Carlo error on the complexity and intensity of estimated Type I error rates. The five panels in each row summarize results for one algorithm (`oMEDUSA`, `tMEDUSA`, and `nMEDUSA`, from top to bottom), the three panels in each column summarize results based on the number of replicates (with 100 to 5, 000 simulated trees, from left to right). The center column—which is reproduced from Figure 2 (the right-most column, with *N* = 10, 000 species)—summarizes the Type I error rates based on 1, 000 simulated trees, where the cells of each panel are colored to reflect the frequency with which spurious diversification-rate shifts were inferred (see FPR legend). These results serve as a reference to explore the impact of the number of simulated replicates on the inferred patterns and overall rates of Type I error. In the other columns, the cells within each panel are colored to reflect the *difference* in the estimated Type I error rate relative to that of its corresponding reference cell (see ∆FPR legend). The standard deviation, *σ*, of the Type I error rate is summarized in the lower right of each panel. These experiments suggest that ~500 replicates are sufficient for precise estimates of Type I error rates. For detailed results, see Table S.3.

**Table S.3:**
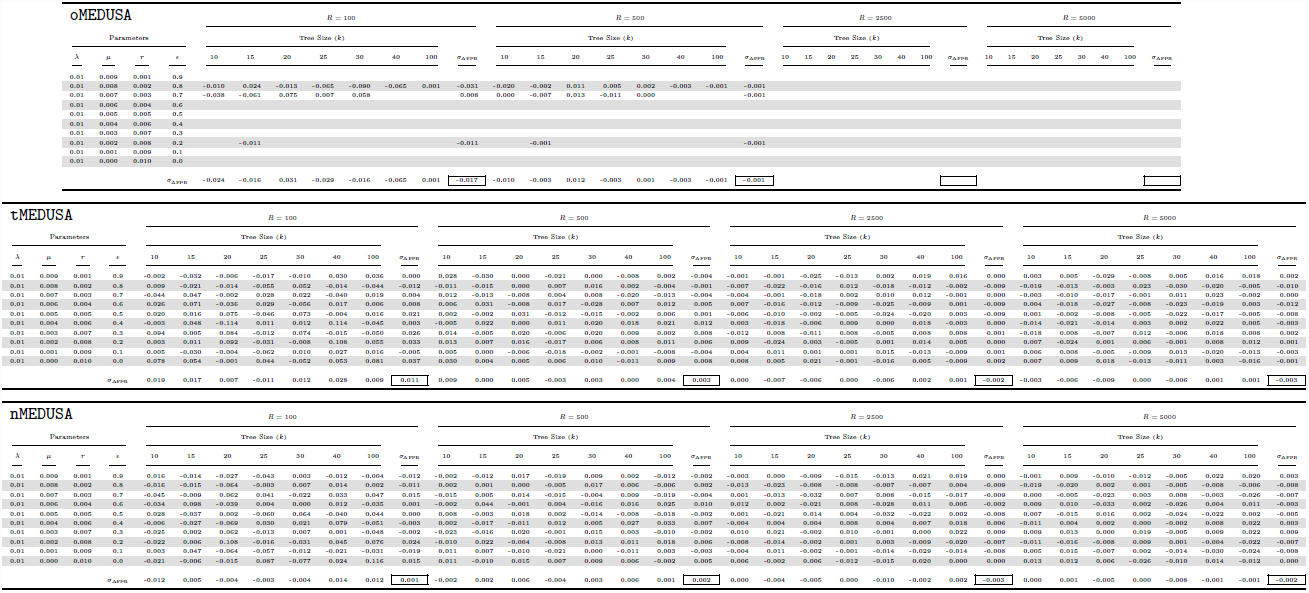
Results of the Monte Carlo Error experiment.

### S.5 The effect of absolute diversification rates on Type I error (stem shifts)

**Figure S.5:**
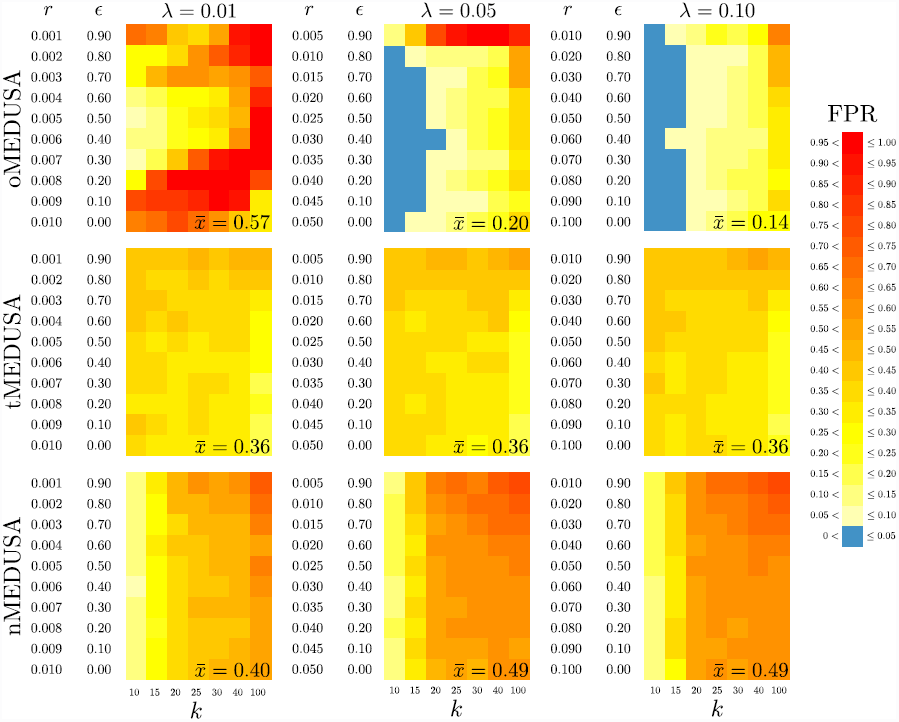
The effect of increasing the absolute diversification rate on Type I error: maintaining consistent relative-extinction rates. The three panels in each row summarize results for one algorithm (`oMEDUSA`, `tMEDUSA`, and `nMEDUSA` from top to bottom), the panels in each column summarize results for one absolute speciation rate (*λ* = 0.01, 0.05, 0.10, from left to right). Trees with *N* = 10, 000 species were simulated under a constant-rate birth-death process using the parameters described in Table S.4. For the absolute rates explored here, we specified a set of extinction rates, *µ*, such that the set of relative-extinction rates, *∈* = 0.00, 0.01,…, 0.90 are consistent with those used in the initial simulation (Figure 2, reproduced here in the left column). Within each panel, we plot the number of terminal lineages, *k*, against the diversification rate, *r*. The cells within each panel are colored to reflect the frequency with which spurious diversification-rate shifts were inferred (see legend). The average Type I error rate is summarized in the lower right of each panel.

**Table S.4:**
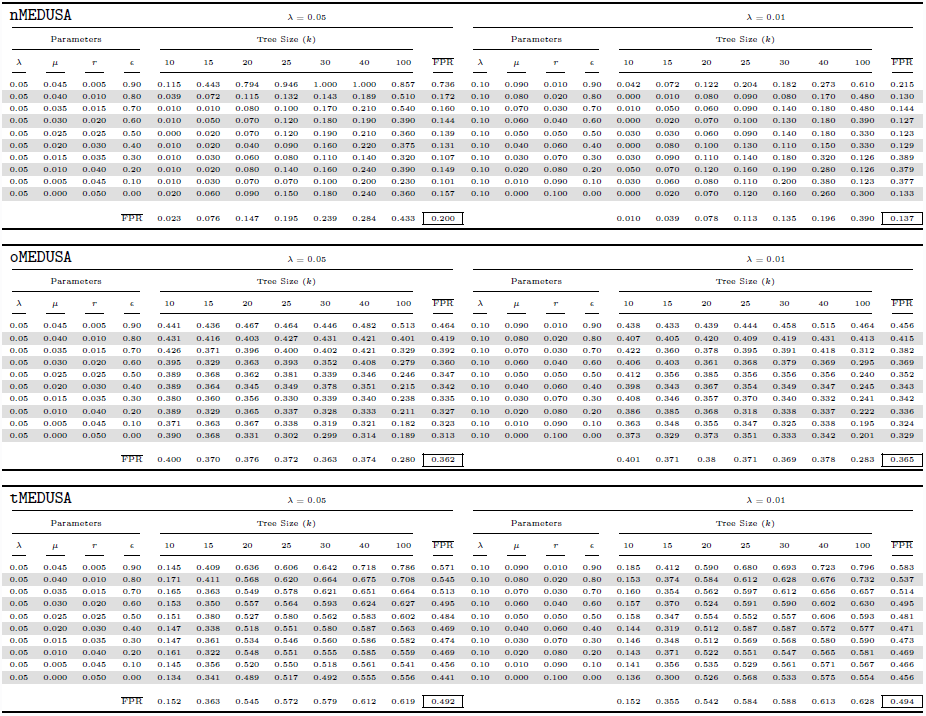
Parameters and Type I error rates for constant-rate simulations with 5-and 10-fold increases in absolute diversification rate.

**Figure S.6:**
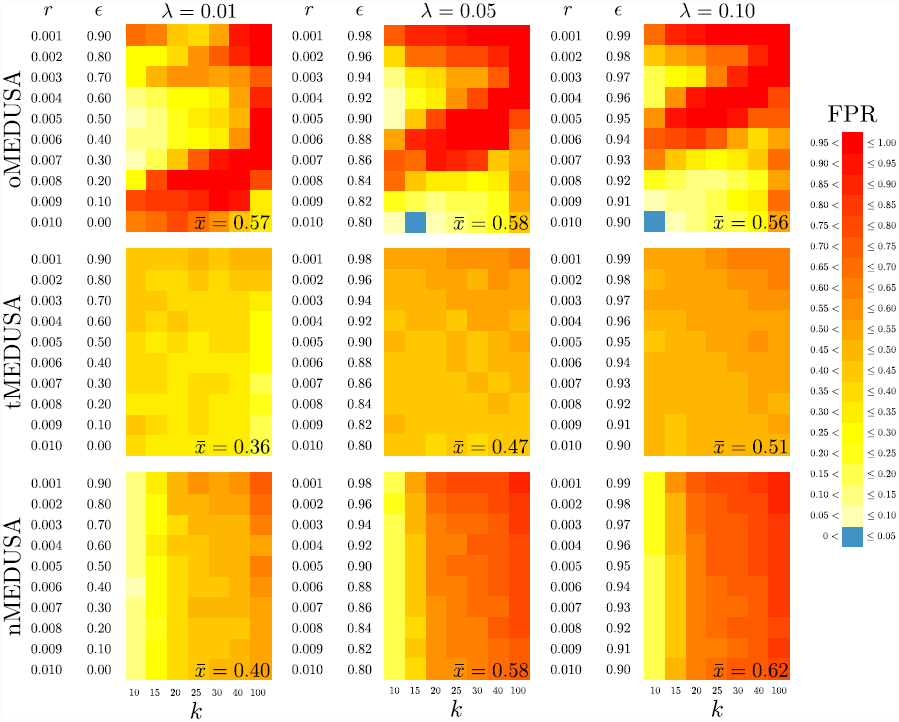
The effect of increasing the absolute diversification rate on Type I error: maintaining consistent net-diversification rates. The three panels in each row summarize results for one algorithm (`oMEDUSA`, `tMEDUSA`, and `nMEDUSA` from top to bottom), the panels in each column summarize results for one absolute speciation rate (*λ* = 0.01, 0.05, 0.10, from left to right). Trees with *N* = 10, 000 species were simulated under a constant-rate birth-death process using the parameters described in Table S.5. For the absolute rates explored here, we specified a set of extinction rates, *µ*, such that the set of net-diversification rates, *r* = 0.001, 0.002,…, 0.010 are consistent with those used in the initial simulation (Figure 2, reproduced here in the left column). Within each panel, we plot the number of terminal lineages, *k*, against the diversification rate, *r*. The cells within each panel are colored to reflect the frequency with which spurious diversification-rate shifts were inferred (see legend). The average Type I error rate is summarized in the lower right of each panel.

**Table S.5:**
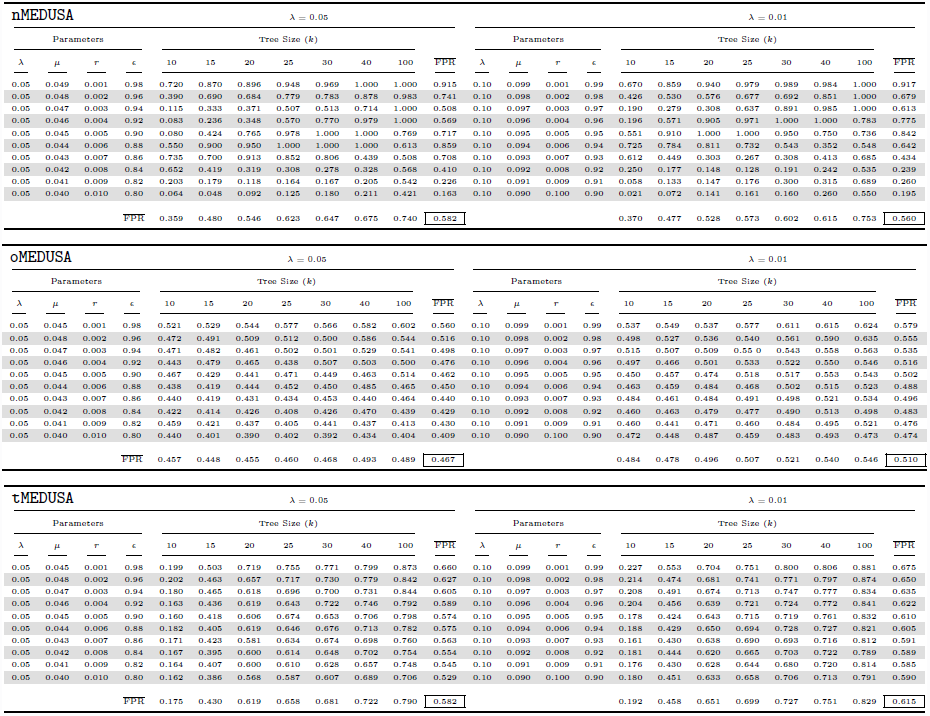
Parameters and Type I error rates for constant-rate simulations with 5-and 10-fold increases in absolute diversification rate.

### S.6 Exploring threshold effects on Type I error rates (stem shifts)

**Table S.6:**
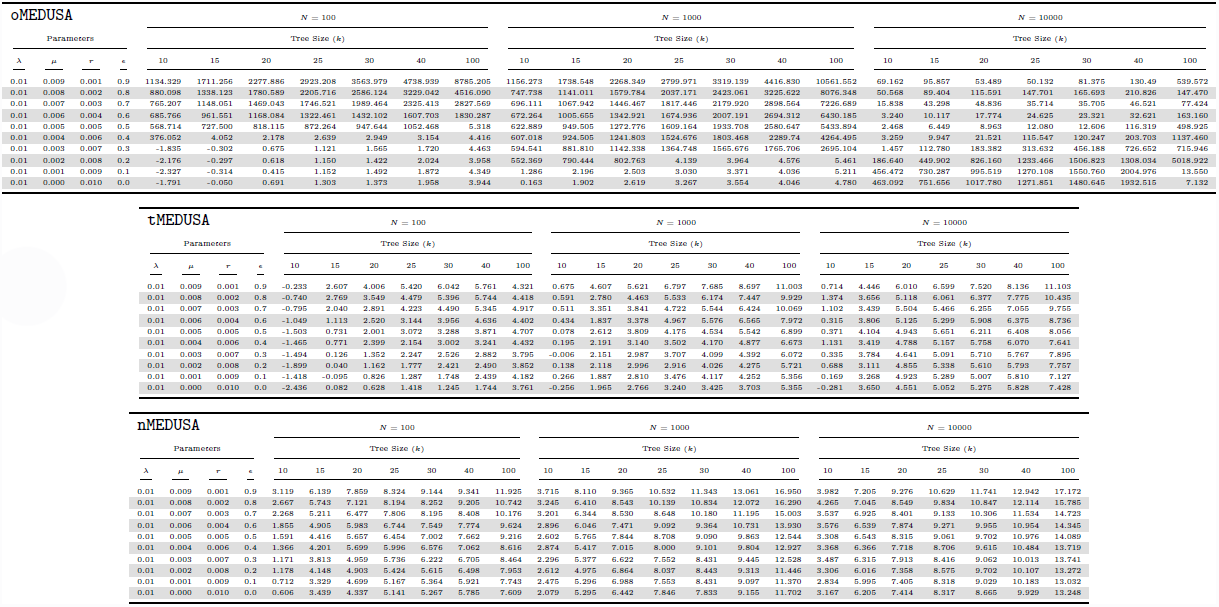
Empirical quantiles for ∆AICc (stem shifts).

### S.7 Exploring Type I error rates for complete species trees (stem shifts)

**Table S.7:** Results of the constant-rate simulation with completely sampled trees.

| Parameters |  |  |  | tMEDUSA |  |  |  |  |  |  |  |  |  | nMEDUSA |  |  |  |  |  |  |  |  |  |
| --- | --- | --- | --- | --- | --- | --- | --- | --- | --- | --- | --- | --- | --- | --- | --- | --- | --- | --- | --- | --- | --- | --- | --- |
| | | | | Tree Size ($k = N$) | | | | | | | | | | Tree Size ($k = N$) | | | | | | | | | |
| $\lambda$ | $\mu$ | $r$ | $\epsilon$ | 50 | 100 | 150 | 200 | 250 | 500 | 1000 | FPR | 50 | 100 | 150 | 200 | 250 | 500 | 1000 | FPR | | | | |
| 0.10 | 0.090 | 0.01 | 0.9 | 0.052 | 0.057 | 0.058 | 0.052 | 0.045 | 0.055 | 0.057 | 0.054 | 0.213 | 0.297 | 0.329 | 0.379 | 0.404 | 0.516 | 0.645 | 0.398 |  |  |  |  |
| 0.10 | 0.080 | 0.02 | 0.8 | 0.056 | 0.054 | 0.050 | 0.046 | 0.058 | 0.045 | 0.046 | 0.051 | 0.193 | 0.271 | 0.303 | 0.325 | 0.390 | 0.451 | 0.541 | 0.353 |  |  |  |  |
| 0.10 | 0.070 | 0.03 | 0.7 | 0.052 | 0.037 | 0.050 | 0.046 | 0.050 | 0.064 | 0.057 | 0.051 | 0.170 | 0.243 | 0.260 | 0.279 | 0.315 | 0.396 | 0.541 | 0.315 |  |  |  |  |
| 0.10 | 0.060 | 0.04 | 0.6 | 0.051 | 0.044 | 0.051 | 0.047 | 0.042 | 0.060 | 0.060 | 0.051 | 0.171 | 0.215 | 0.254 | 0.251 | 0.284 | 0.377 | 0.491 | 0.292 |  |  |  |  |
| 0.10 | 0.050 | 0.05 | 0.5 | 0.062 | 0.044 | 0.036 | 0.050 | 0.039 | 0.045 | 0.065 | 0.049 | 0.167 | 0.178 | 0.206 | 0.229 | 0.262 | 0.350 | 0.435 | 0.261 |  |  |  |  |
| 0.10 | 0.040 | 0.06 | 0.4 | 0.038 | 0.045 | 0.044 | 0.041 | 0.039 | 0.038 | 0.045 | 0.041 | 0.141 | 0.174 | 0.206 | 0.197 | 0.230 | 0.314 | 0.354 | 0.231 |  |  |  |  |
| 0.10 | 0.030 | 0.07 | 0.3 | 0.046 | 0.045 | 0.037 | 0.048 | 0.053 | 0.040 | 0.035 | 0.043 | 0.129 | 0.173 | 0.199 | 0.188 | 0.229 | 0.271 | 0.328 | 0.217 |  |  |  |  |
| 0.10 | 0.020 | 0.08 | 0.2 | 0.041 | 0.038 | 0.040 | 0.041 | 0.032 | 0.034 | 0.050 | 0.039 | 0.124 | 0.131 | 0.168 | 0.194 | 0.210 | 0.255 | 0.331 | 0.202 |  |  |  |  |
| 0.10 | 0.010 | 0.09 | 0.1 | 0.030 | 0.032 | 0.032 | 0.035 | 0.036 | 0.041 | 0.049 | 0.036 | 0.126 | 0.129 | 0.168 | 0.165 | 0.196 | 0.242 | 0.311 | 0.191 |  |  |  |  |
| 0.10 | 0.000 | 0.10 | 0.0 | 0.029 | 0.032 | 0.028 | 0.031 | 0.042 | 0.032 | 0.030 | 0.032 | 0.115 | 0.124 | 0.150 | 0.151 | 0.167 | 0.207 | 0.251 | 0.166 |  |  |  |  |
| FPR |  |  |  | 0.046 | 0.043 | 0.043 | 0.044 | 0.044 | 0.045 | 0.049 | 0.045 | 0.155 | 0.194 | 0.224 | 0.236 | 0.269 | 0.338 | 0.423 | 0.263 |  |  |  |  |

### S.8 Assessing the impact of terminal unresolved clades (stem shifts)

**Figure S.7:**
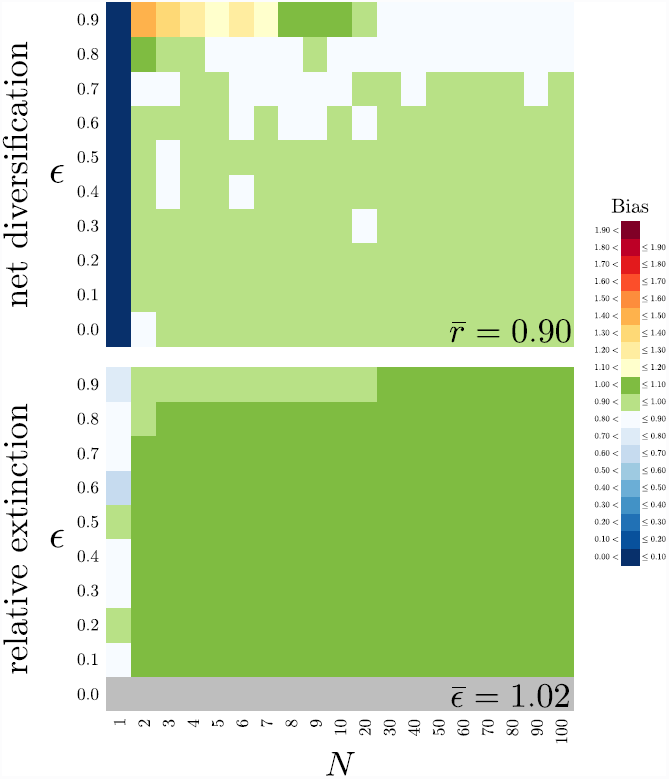
The size of terminal unresolved lineages effects bias in parameter estimates. Trees of size *N* ranging from 1 to 1000 were simulated under a constant-rate birth-death process using parameters summarized in Tables S.8 and S.9. For each simulated tree, we estimated the net-diversification rate (upper panel) and relative-extinction rate (lower panel) using equation 2. Each panel plots lineage size, *N*, against the relative-extinction rate, *∈*, used in the simulation. The cells within each panel are colored as a heat map reflecting the difference between the estimated and true rates (see Bias legend). The average bias in the parameter estimates is summarized in the lower right of each panel.

**Table S.8:**
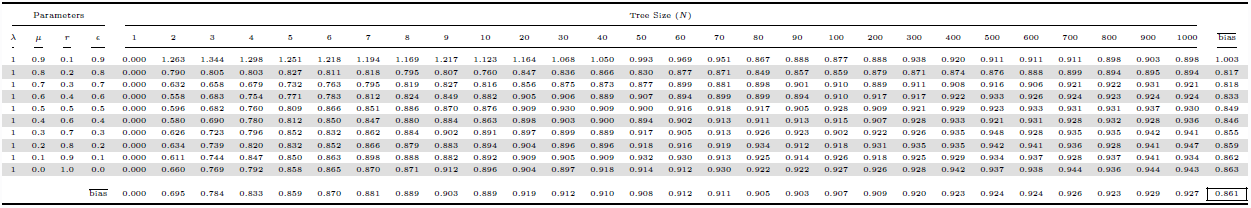
Estimated net-diversification rates of unresolved clades simulated under a constant-rate birth-death process.

**Table S.9:**
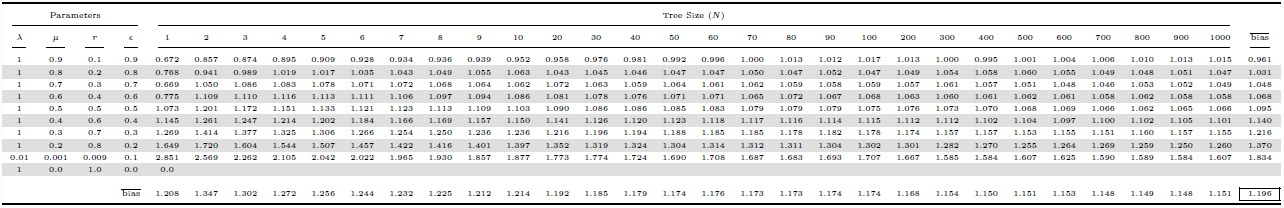
Estimated relative-extinction rates of unresolved clades simulated under a constant-rate birth-death process.

**Figure S.8:**
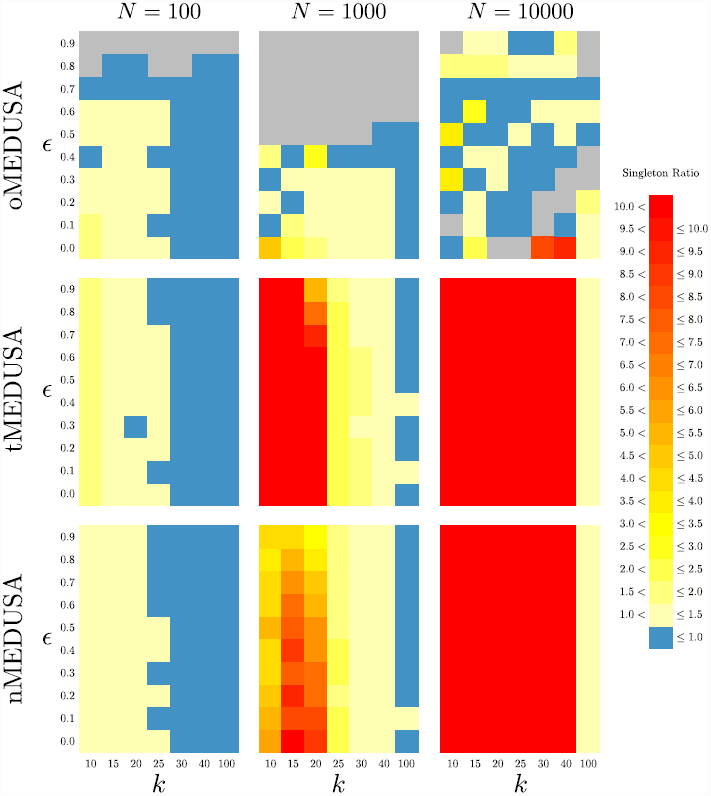
Relative frequency of singleton terminal linaeages in trees with spurious rate shifts versus trees without rate shifts. The three panels in each row summarize results for one algorithm (`oMEDUSA`, `tMEDUSA`, and `nMEDUSA`, from top to bottom), the three panels in each column summarize results for one tree size (with *N* = 100, 1, 000, 10, 000 species, from left to right). Within each panel, we plot the number of terminal lineages, *k*, against the relative-extinction rate used in the simulation. Trees were simulated under a constant-rate birth-death process using the parameters summarized in Table S.2. The cells within each panel are colored as a heat map reflecting the ratio of the frequency of ‘singletons’ (terminal lineages with a single species) in trees where `MEDUSA` incorrectly identified a rate shift versus the singleton frequency in trees where it correctly identified no rate shift (see Singleton Ratio legend).

**Table S.10:**
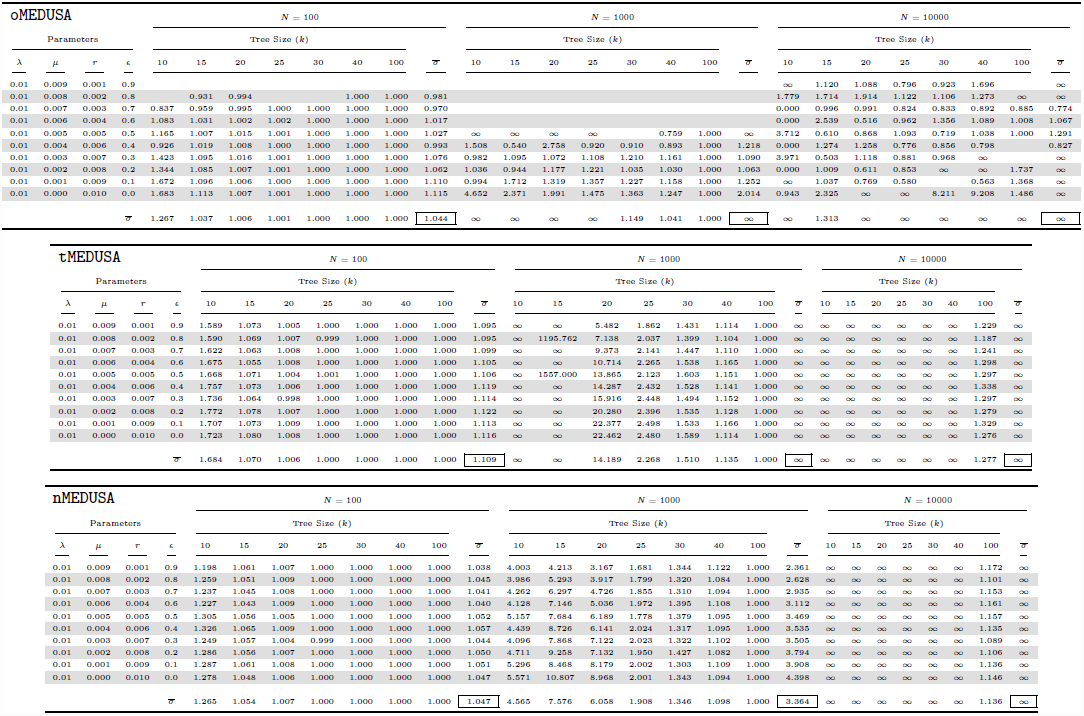
Relative frequency of singletons in trees with rate shifts againts trees without rate shifts.

### S.9 Parameter estimation (stem shifts)

**Table S.11:**
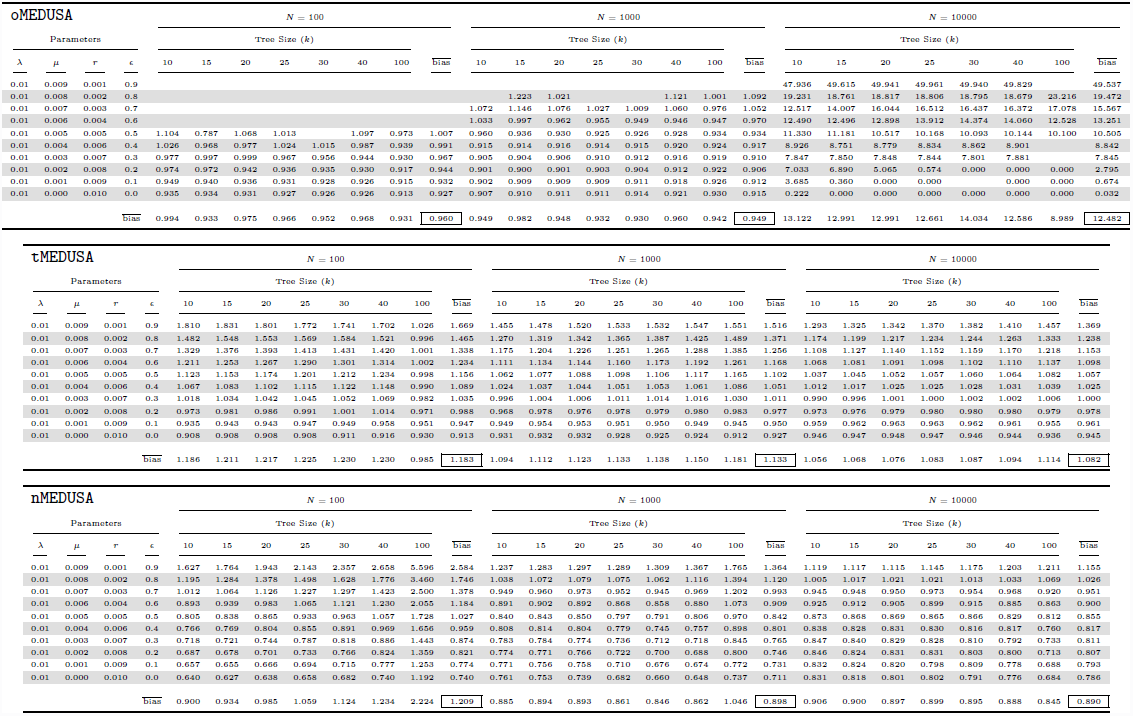
Estimated net-diversification rates for the initial constant-rate simulations when a single-rate model is selected.

**Figure S.9:**
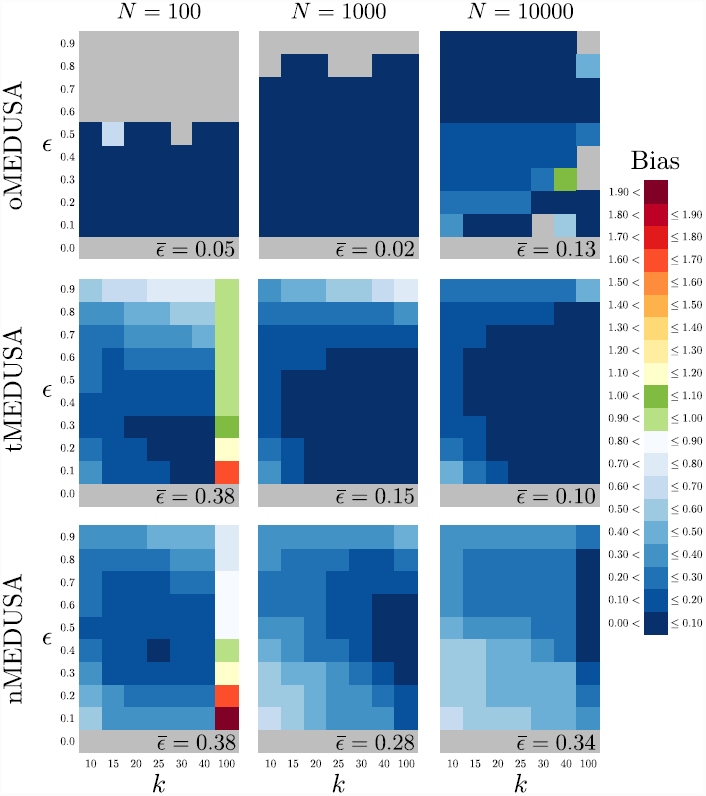
Bias in estimated relative-extinction rate when a single-rate model is selected. The three panels in each row summarize results for one algorithm (`oMEDUSA`, `tMEDUSA`, and `nMEDUSA`, from top to bottom), the three panels in each column summarize results for one tree size (with *N* = 100, 1, 000, 10, 000 species, from left to right). Within each panel, we plot the number of terminal lineages, *k*, against the relative-extinction rate used in the simulation. Trees were simulated under a constant-rate birth-death process using the parameters summarized in Table S.12. The cells within each panel are colored to reflect the average difference between the estimated and true relative-extinction rates when a one-rate model is correctly identified (see Bias legend). The average bias in estimates of relative extinction-rate is summarized in the lower right of each panel.

**Table S.12:**
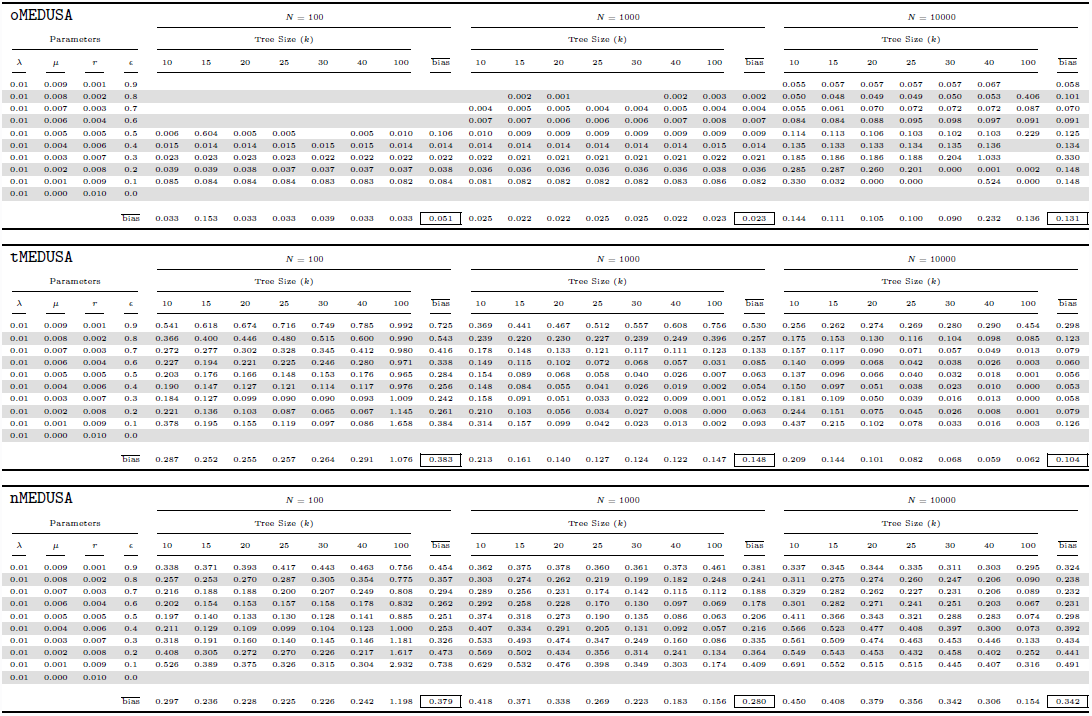
Estimated relative-extinction rates for the initial constant-rate simulations when a single-rate model is selected.

**Figure S.10:**
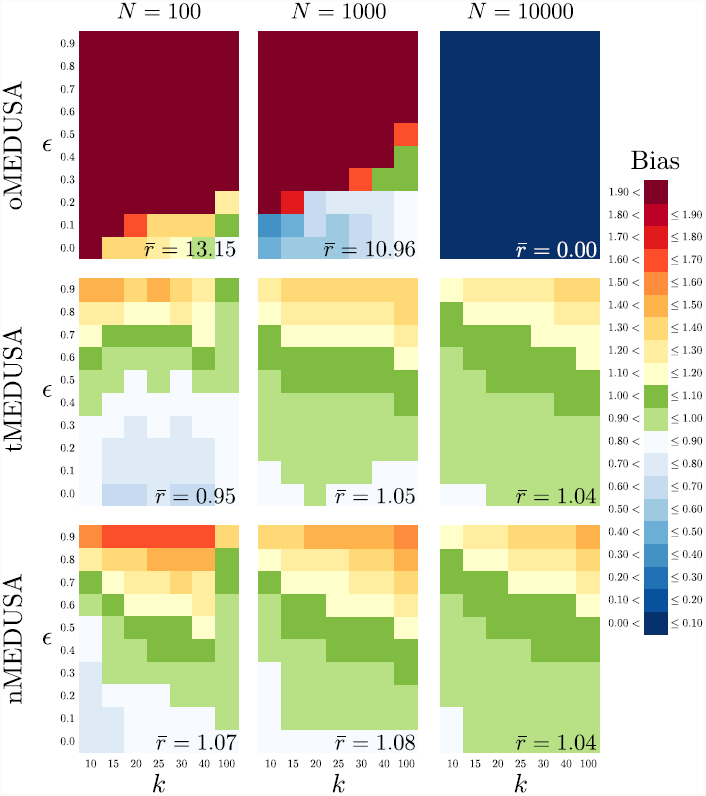
Bias in estimated background net-diversification rates when a two-rate model is selected. The three panels in each row summarize results for one algorithm (`oMEDUSA`, `tMEDUSA`, and `nMEDUSA`, from top to bottom), the three panels in each column summarize results for one tree size (with *N* = 100, 1, 000, 10, 000 species, from left to right). Within each panel, we plot the number of terminal lineages, *k*, against the relative-extinction rate used in the simulation. Trees were simulated under a constant-rate birth-death process using the parameters summarized in Table S.13. The cells within each panel are colored to reflect the average ratio of the estimated *vs.* true background net-diversification rate when a two-rate model is incorrectly identified (see Bias legend). The average bias in the inferred background net-diversification rate is summarized in the lower right of each panel.

**Table S.13:**
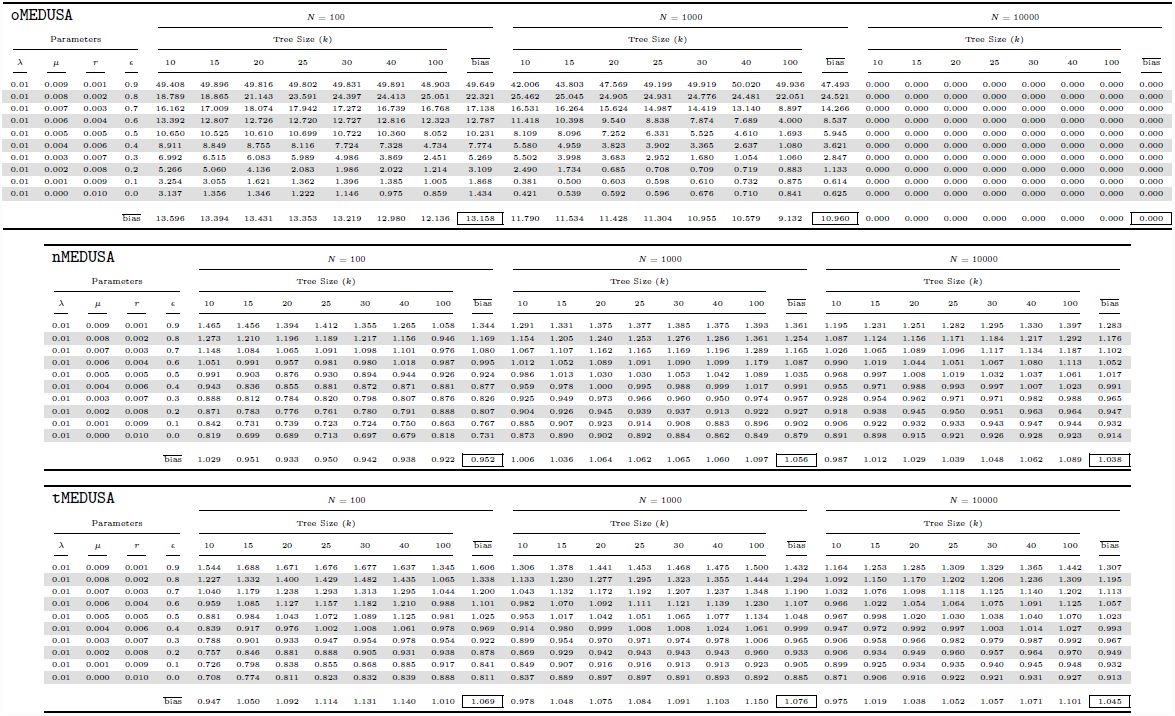
Estimated background net-diversification rates for the initial constant-rate simulations when a two-rate model is selected.

**Figure S.11:**
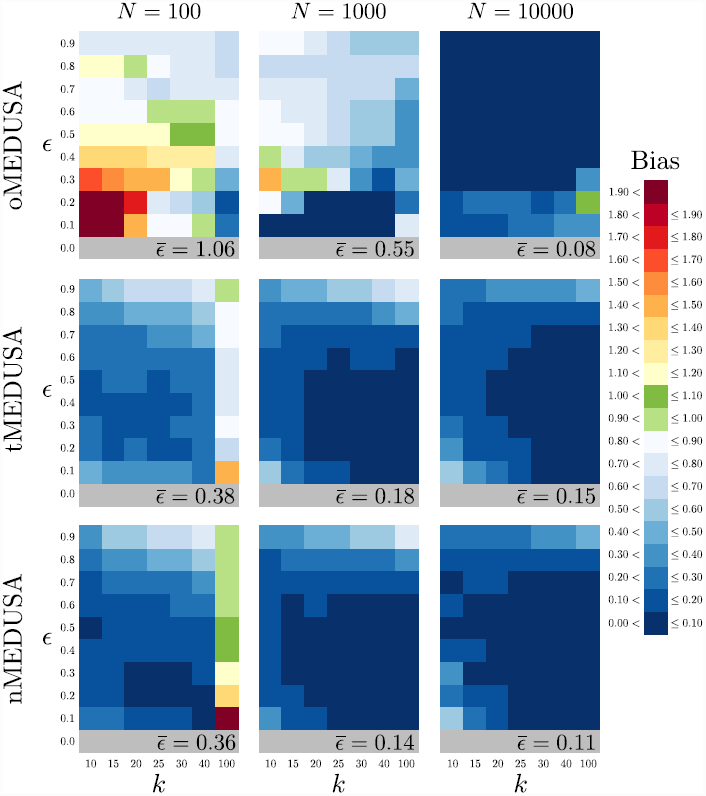
Bias in estimated background relative-extinction rates when a two-rate model is selected. The three panels in each row summarize results for one algorithm (`oMEDUSA`, `tMEDUSA`, and `nMEDUSA`, from top to bottom), the three panels in each column summarize results for one tree size (with *N* = 100, 1, 000, 10, 000 species, from left to right). Within each panel, we plot the number of terminal lineages, *k*, against the relative-extinction rate used in the simulation. Trees were simulated under a constant-rate birth-death process using the parameters summarized in Table S.14. The cells within each panel are colored to reflect the average ratio of the estimated *vs.* true background relative-extinction rate when a two-rate model is incorrectly identified (see Bias legend). The average bias in the inferred background relative-extinction rate is summarized in the lower right of each panel.

**Table S.14:**
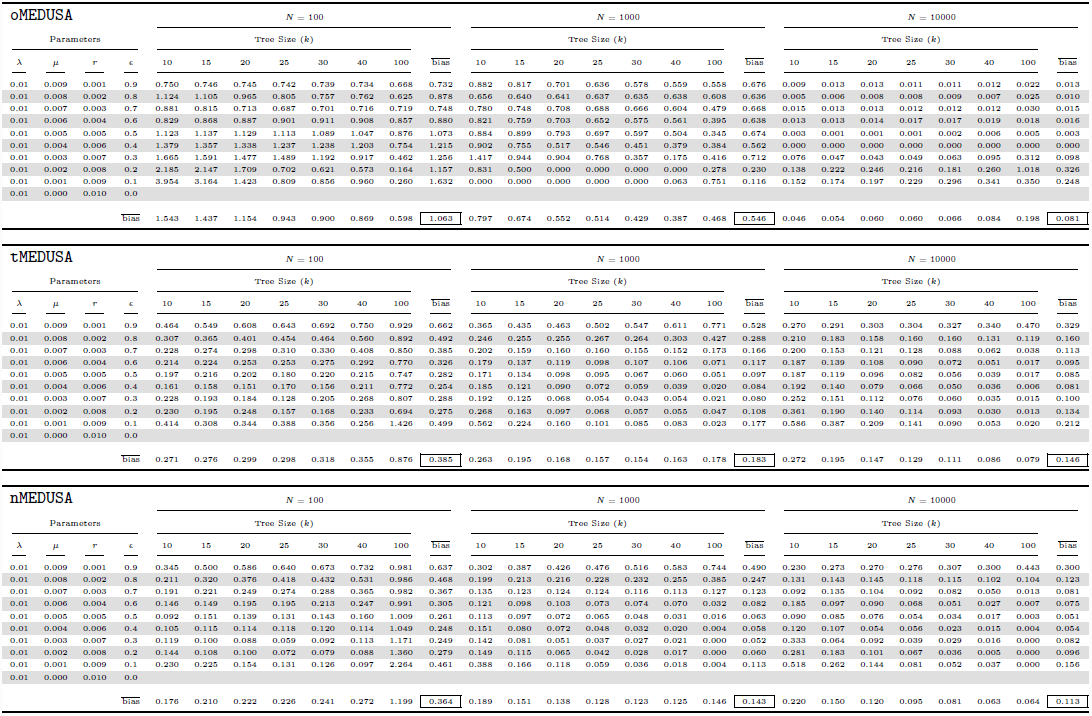
Estimated backgound relative-extinction rates for the initial constant-rate simulations when a two-rate model is selected.

**Table S.15:**
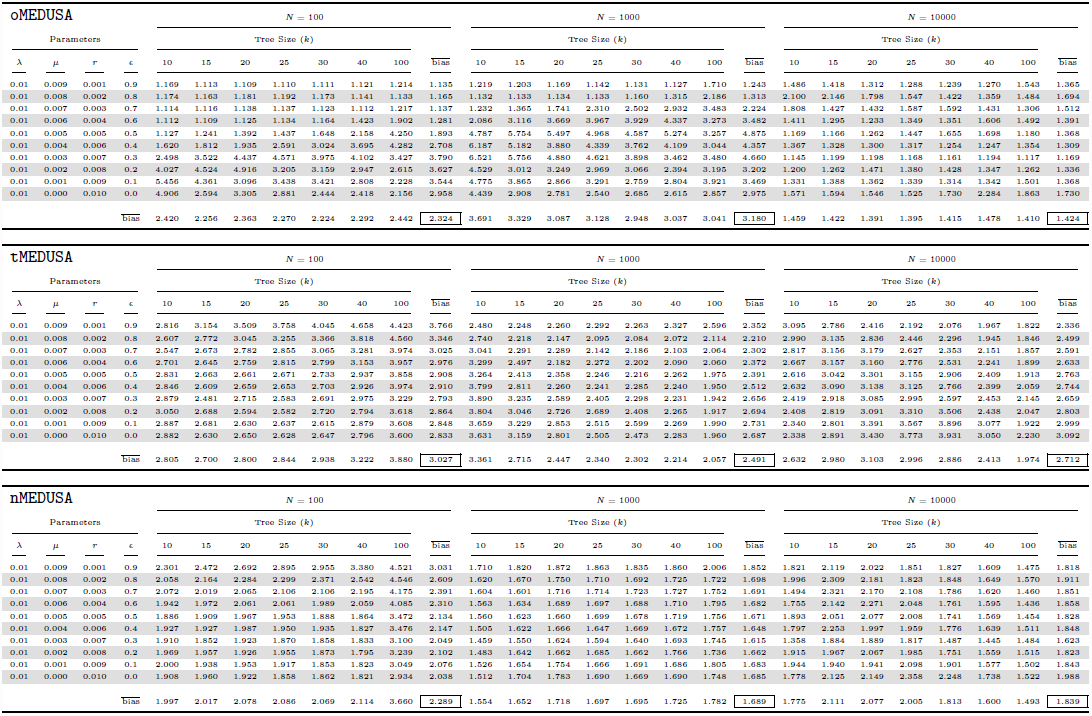
Estimated magnitude of net diversification-rate shifts for the initial constant-rate simulations when a two-rate model is selected.

**Figure S.12:**
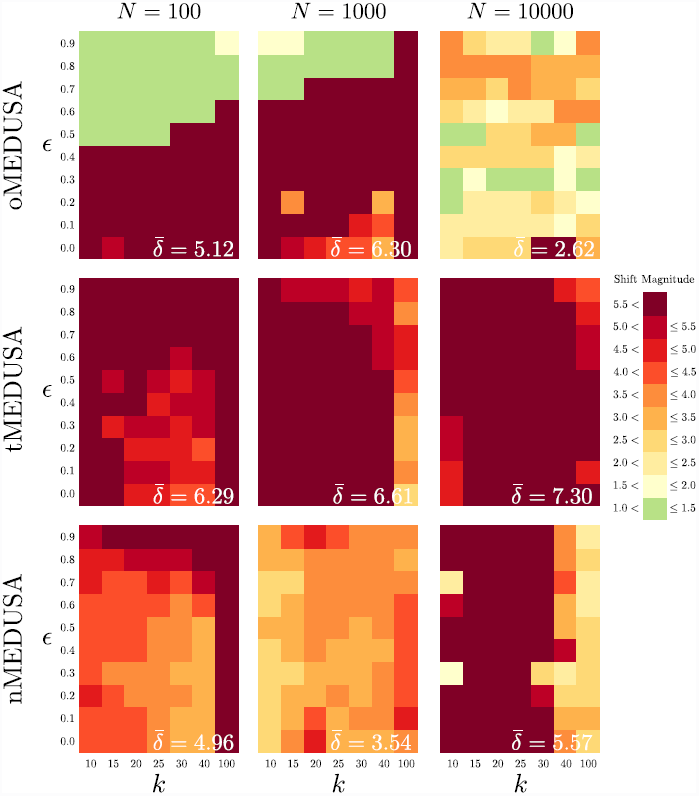
95% quantile of shift magnitude in net-diversification rate when a two-rate model is selected. The three panels in each row summarize results for one algorithm (`oMEDUSA`, `tMEDUSA`, and `nMEDUSA`, from top to bottom), the three panels in each column summarize results for one tree size (with *N* = 100, 1, 000, 10, 000 species, from left to right). Within each panel, we plot the number of terminal lineages, *k*, against the relative-extinction rate used in the simulation. Trees were simulated under a constant-rate birth-death process using the parameters summarized in Table S.16. The cells within each panel are colored to reflect the 95% quantile of the magnitude of the inferred shift in net-diversification rate when a two-rate model is incorrectly identified (see Shift Magnitude legend). In order to accommodate the bimodal distribution of rate shifts, the magnitude of each net-diversification-rate shift was computed as the ratio of the higher rate to the lower rate, regardless of which rate corresponded to the background-rate category. The average 95% quantile of the magnitude of shifts in net-diversification rate shift is summarized in the lower right of each panel.

**Table S.16:**
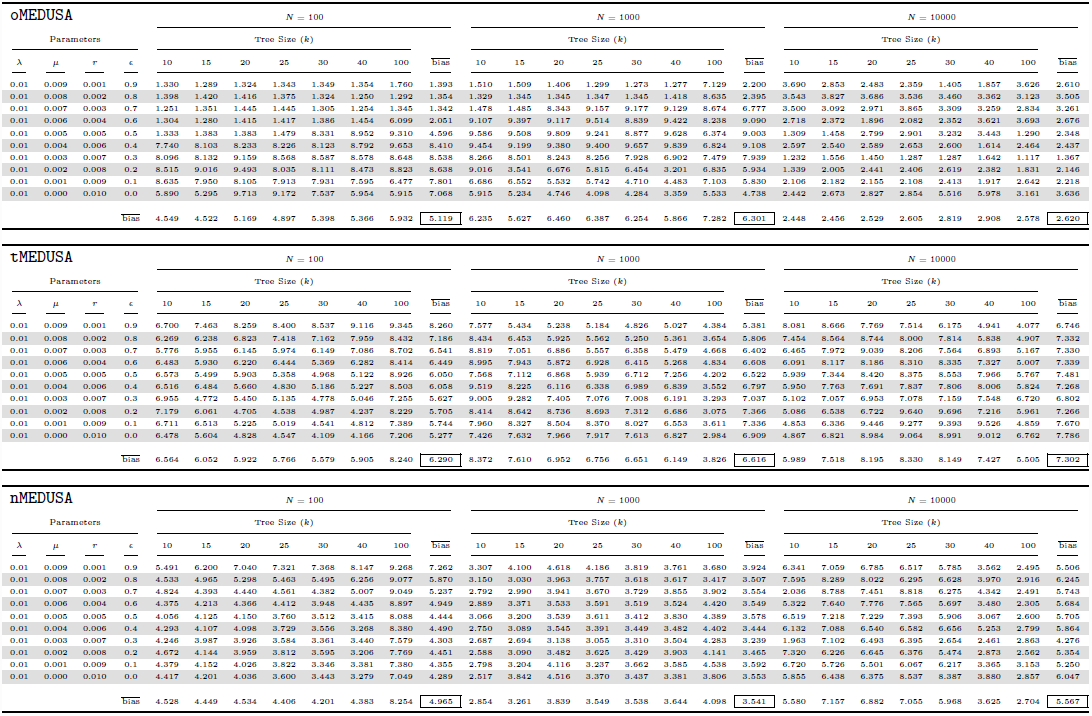
95% quantile of estimated magnitude of net diversification-rate shifts for the initial constant-rate simulations when a two-rate model is selected.

**Figure S.13:**
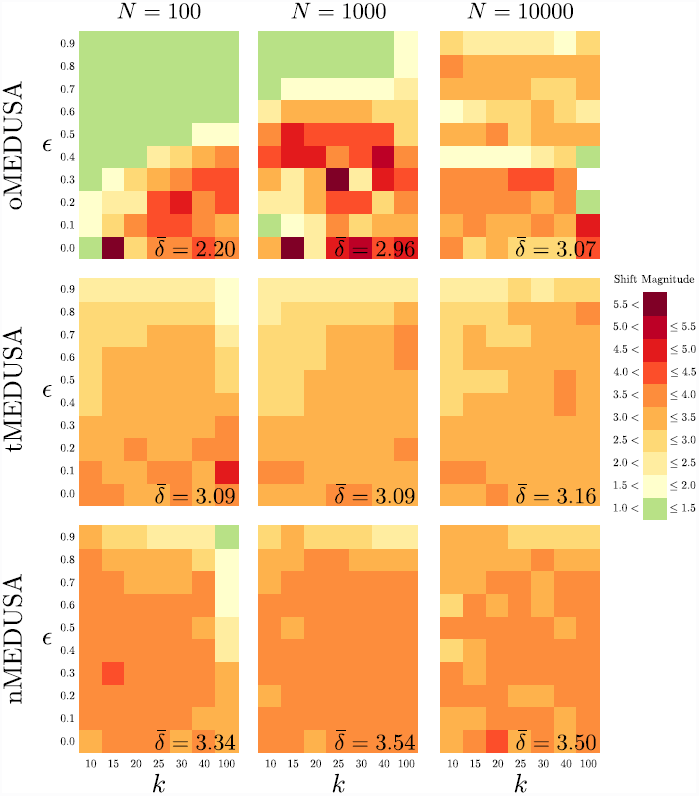
Magnitude of shift in relative-extinction rate when a two-rate model is selected. The three panels in each row summarize results for one algorithm (`oMEDUSA`, `tMEDUSA`, and `nMEDUSA`, from top to bottom), the three panels in each column summarize results for one tree size (with *N* = 100, 1, 000, 10, 000 species, from left to right). Within each panel, we plot the number of terminal lineages, *k*, against the relative-extinction rate used in the simulation. Trees were simulated under a constant-rate birth-death process using the parameters summarized in Table S.17. The cells within each panel are colored to reflect the average magnitude of the inferred shift in relative-extinction rate when a two-rate model is incorrectly identified (see Shift Magnitude legend). In order to accommodate the bimodal distribution of rate shifts, the magnitude of each relative-extinction-rate shift was computed as the ratio of the higher rate to the lower rate, regardless of which rate corresponded to the background-rate category. The average magnitude of shifts in relative-extinction rate shift is summarized in the lower right of each panel.

**Table S.17:**
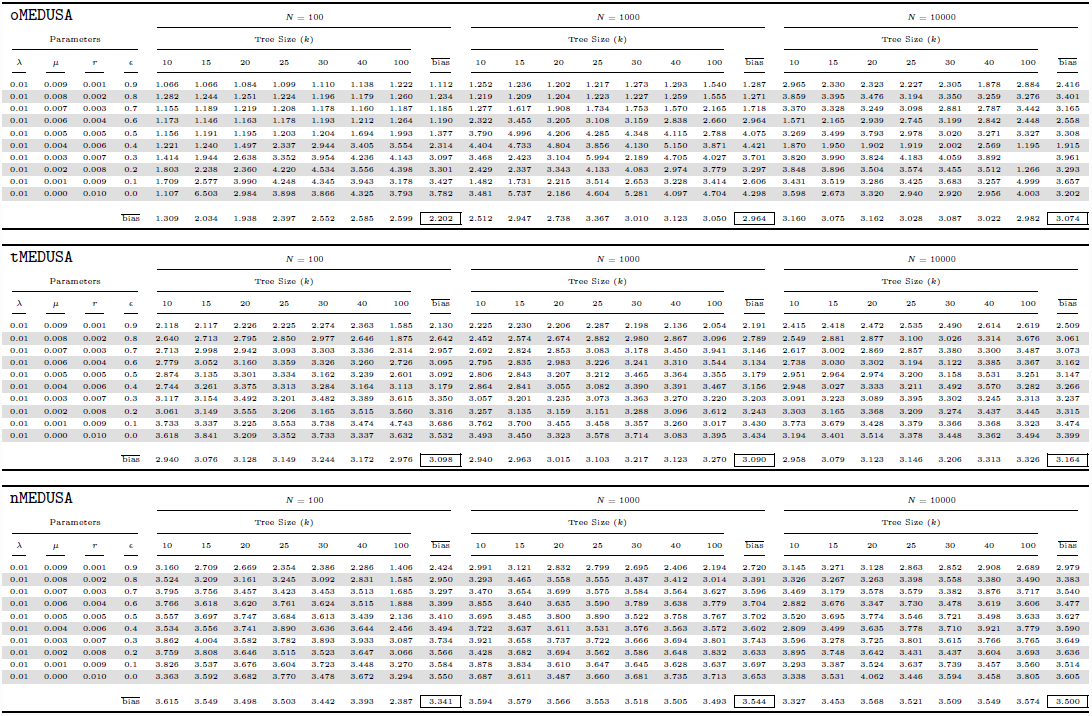
Estimated magnitude of relative-extinction rate shifts for the initial constant-rate simulations when a two-rate model is selected.

**Figure S.14:**
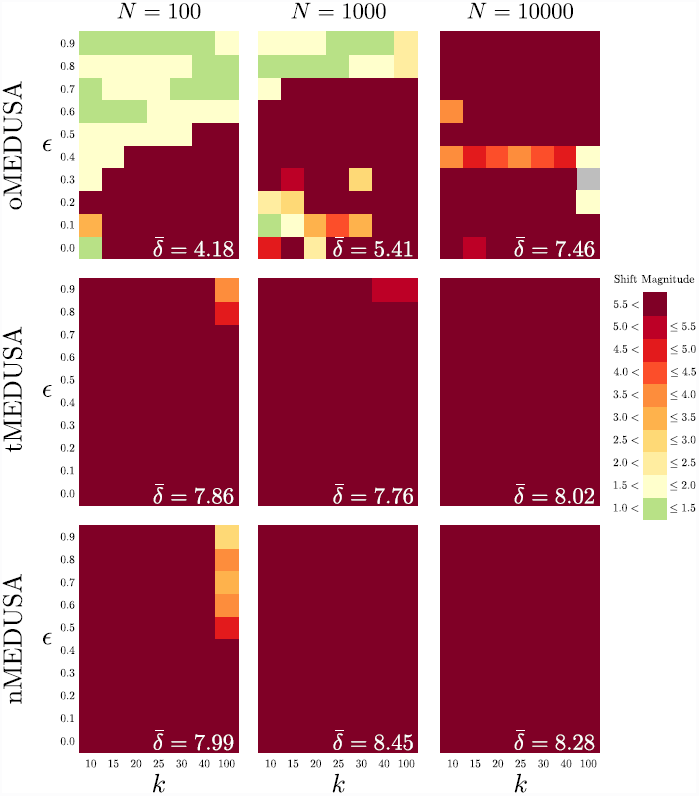
95% quantile of shift magnitude in relative-extinction rate when a two-rate model is selected. The three panels in each row summarize results for one algorithm (`oMEDUSA`, `tMEDUSA`, and `nMEDUSA`, from top to bottom), the three panels in each column summarize results for one tree size (with *N* = 100, 1, 000, 10, 000 species, from left to right). Within each panel, we plot the number of terminal lineages, *k*, against the relative-extinction rate used in the simulation. Trees were simulated under a constant-rate birth-death process using the parameters summarized in Table S.18. The cells within each panel are colored to reflect the 95% quantile of the magnitude of the inferred shift in relative-extinction rate when a two-rate model is incorrectly identified (see Shift Magnitude legend). In order to accommodate the bimodal distribution of rate shifts, the magnitude of each relative-extinction-rate shift was computed as the ratio of the higher rate to the lower rate, regardless of which rate corresponded to the background-rate category. The average 95% quantile of the magnitude of shifts in relative-extinction rate shift is summarized in the lower right of each panel.

**Table S.18:**
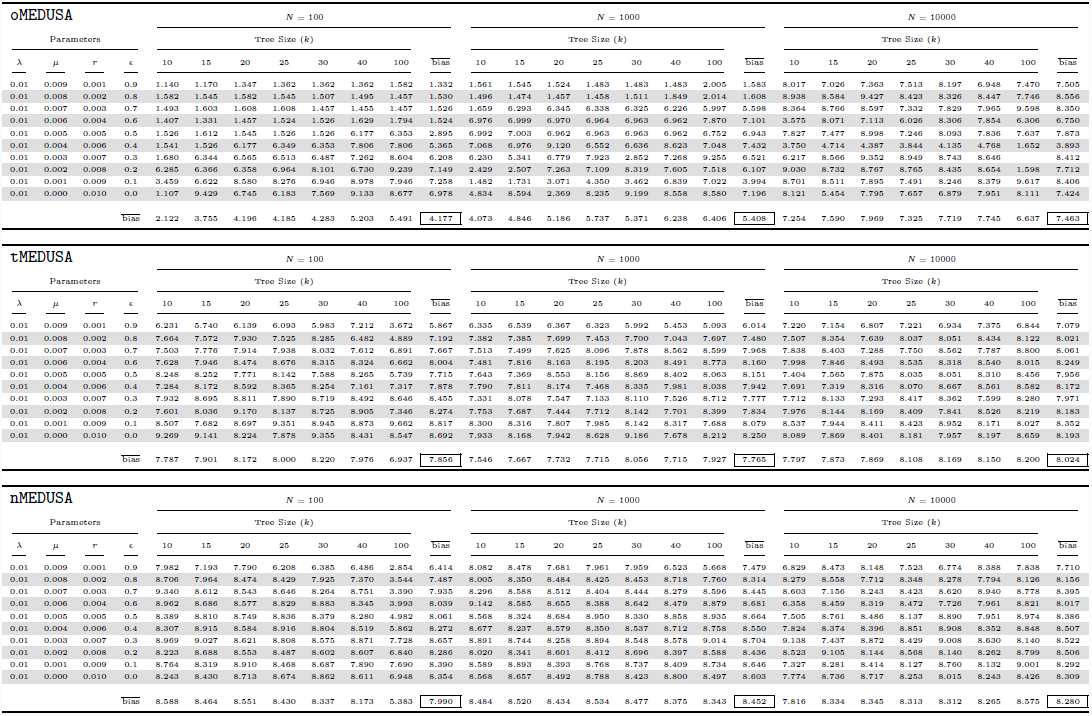
95% quantile of estimated magnitude of relative-extinction-rate shifts for the initial constant-rate simulations when a two-rate model is selected.

### S.10 The effect of relative-extinction rate, tree size, and phylogenetic resolution (crown shifts).

**Figure S.15:**
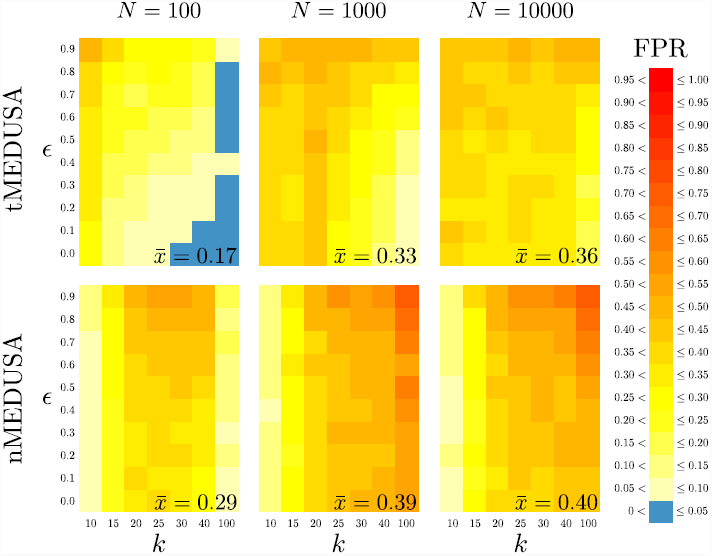
Type I error rates for the three MEDUSA algorithms (crown shifts). The three panels in each row summarize results for one algorithm (`oMEDUSA`, `tMEDUSA`, and `nMEDUSA`, from top to bottom), the three panels in each column summarize results for one tree size (with *N* = 100, 1, 000, 10, 000 species, from left to right). Within each panel, we plot the number of terminal lineages, *k*, against the relative-extinction rate used in the simulation. Trees were simulated under a constant-rate birth-death process using the parameters summarized in Table S.19. The cells within each panel are colored as a heat map reflecting the frequency with which spurious diversification-rate shifts were inferred (see legend). The average Type I error rate is summarized in the lower right of each panel.

**Table S.19:**
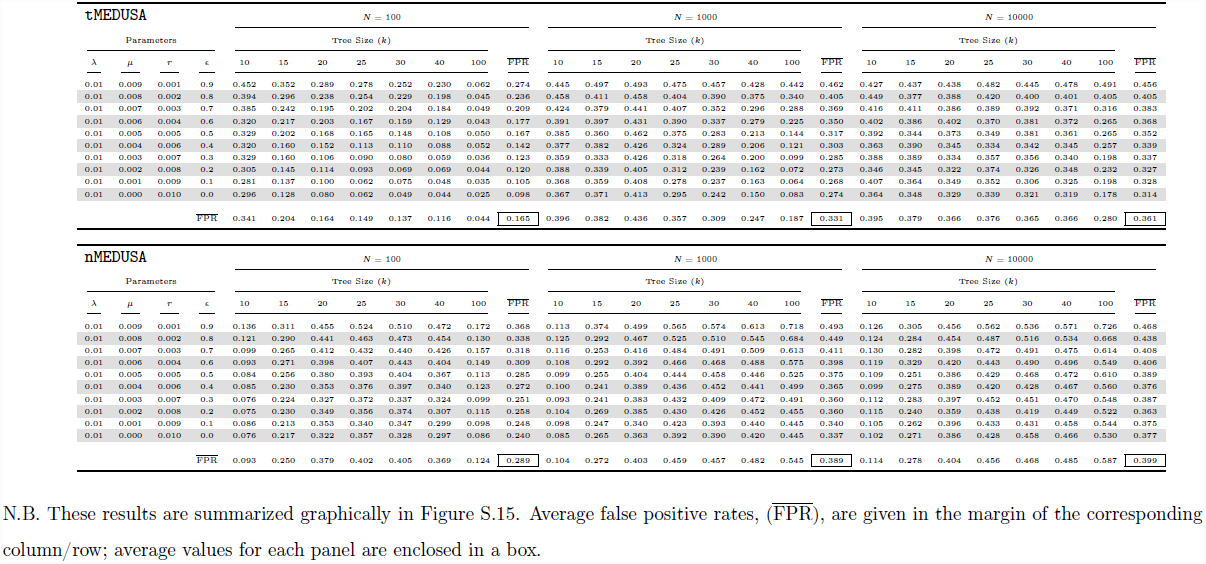
Parameters and estimated Type 1 error rates for the initial constant-rate simulations (crown shifts)

### S.11 Exploring the impact of Monte Carlo error on Type I error rates (crown shifts)

**Figure S.16:**
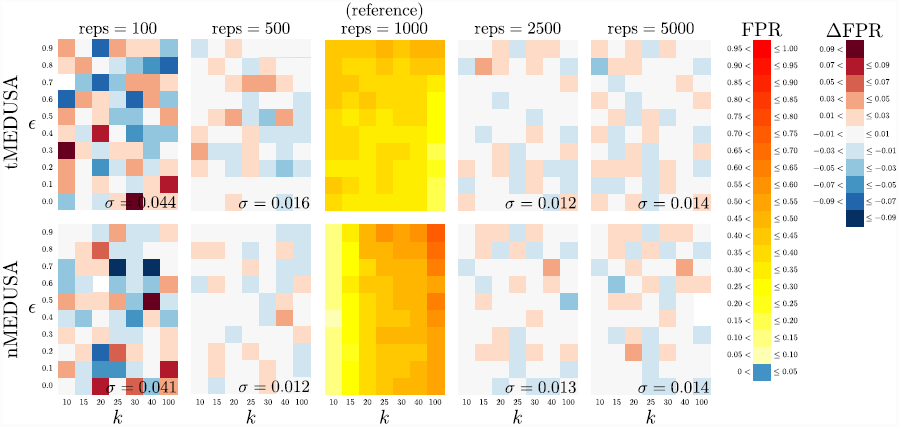
Exploring the impact of Monte Carlo error on the complexity and intensity of estimated Type I error rates. The five panels in each row summarize results for one algorithm (`tMEDUSA` and `nMEDUSA`, from top to bottom), the three panels in each column summarize results based on the number of replicates (with 100 to 5, 000 simulated trees, from left to right). The center column—which is reproduced from Figure 2 (the right-most column, with *N* = 10, 000 species)—summarizes the Type I error rates based on 1, 000 simulated trees, where the cells of each panel are colored to reflect the frequency with which spurious diversification-rate shifts were inferred (see FPR legend). These results serve as a reference to explore the impact of the number of simulated replicates on the inferred patterns and overall rates of Type I error. In the other columns, the cells within each panel are colored to reflect the *difference* in the estimated Type I error rate relative to that of its corresponding reference cell (see ∆FPR legend). The standard deviation, *σ*, of the Type I error rate is summarized in the lower right of each panel. These experiments suggest that ~500 replicates are sufficient for precise estimates of Type I error rates. For detailed results, see Table S.20.

**Table S.20:**
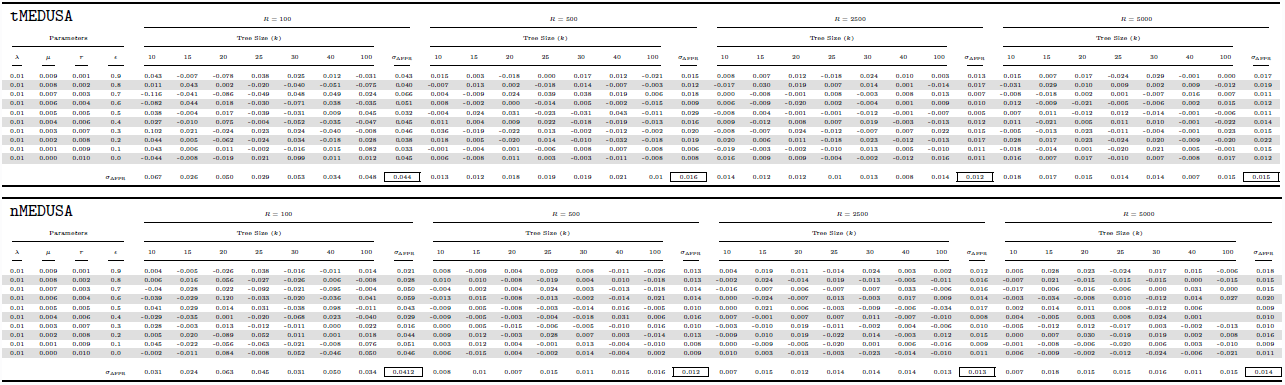
Results of the Monte Carlo Error experiment.

### S.12 The effect of absolute diversification rates on Type I error (crown shifts)

**Figure S.17:**
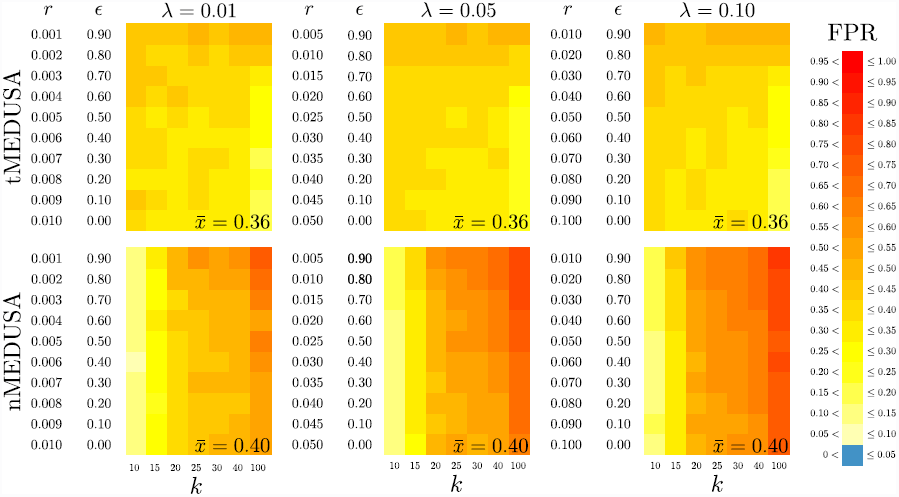
The effect of increasing the absolute diversification rate on Type I error: maintaining consistent relative-extinction rates. The three panels in each row summarize results for one algorithm (`tMEDUSA`, and `nMEDUSA` from top to bottom), the panels in each column summarize results for one absolute speciation rate (*λ* = 0.01, 0.05, 0.10, from left to right). Trees with *N* = 10, 000 species were simulated under a constant-rate birth-death process using the parameters described in Table S.21. For the absolute rates explored here, we specified a set of extinction rates, *µ*, such that the set of relative-extinction rates, *∈* = 0.00, 0.01,…, 0.90 are consistent with those used in the initial simulation (Figure 2, reproduced here in the left column). Within each panel, we plot the number of terminal lineages, *k*, against the diversification rate, *r*. The cells within each panel are colored to reflect the frequency with which spurious diversification-rate shifts were inferred (see legend). The average Type I error rate is summarized in the lower right of each panel.

**Table S.21:**
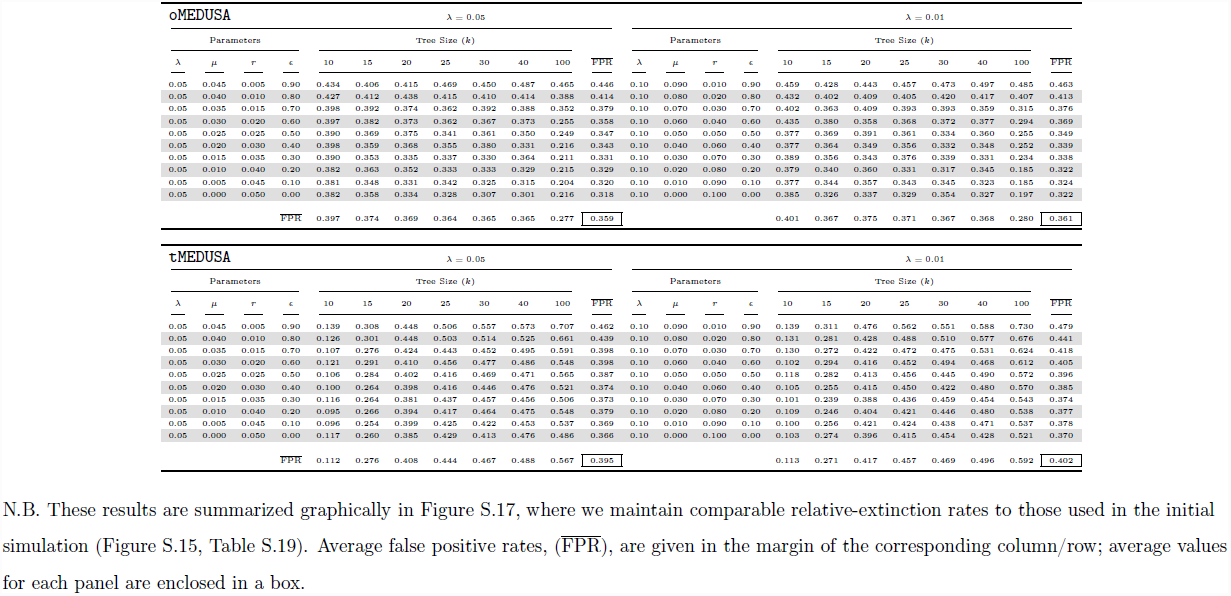
Parameters and Type I error rates for constant-rate simulations with 5-and 10-fold increases in absolute diversification rate.

**Figure S.18:**
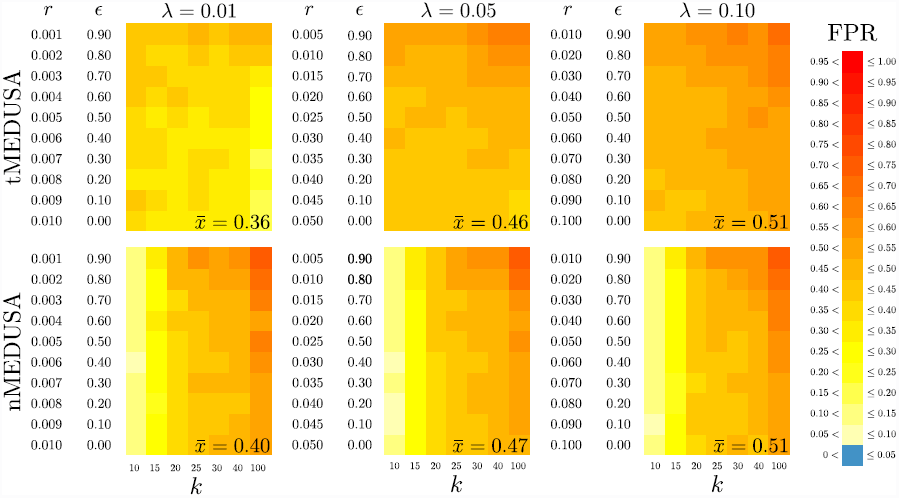
The effect of increasing the absolute diversification rate on Type I error: maintaining consistent net-diversification rates. The three panels in each row summarize results for one algorithm (`tMEDUSA`, and `nMEDUSA` from top to bottom), the panels in each column summarize results for one absolute speciation rate (*λ* = 0.01, 0.05, 0.10, from left to right). Trees with *N* = 10, 000 species were simulated under a constant-rate birth-death process using the parameters described in Table S.22. For the absolute rates explored here, we specified a set of extinction rates, *µ*, such that the set of net-diversification rates, *r* = 0.001, 0.002,…, 0.010 are consistent with those used in the initial simulation (Figure 2, reproduced here in the left column). Within each panel, we plot the number of terminal lineages, *k*, against the diversification rate, *r*. The cells within each panel are colored to reflect the frequency with which spurious diversification-rate shifts were inferred (see legend). The average Type I error rate is summarized in the lower right of each panel.

**Table S.22:**
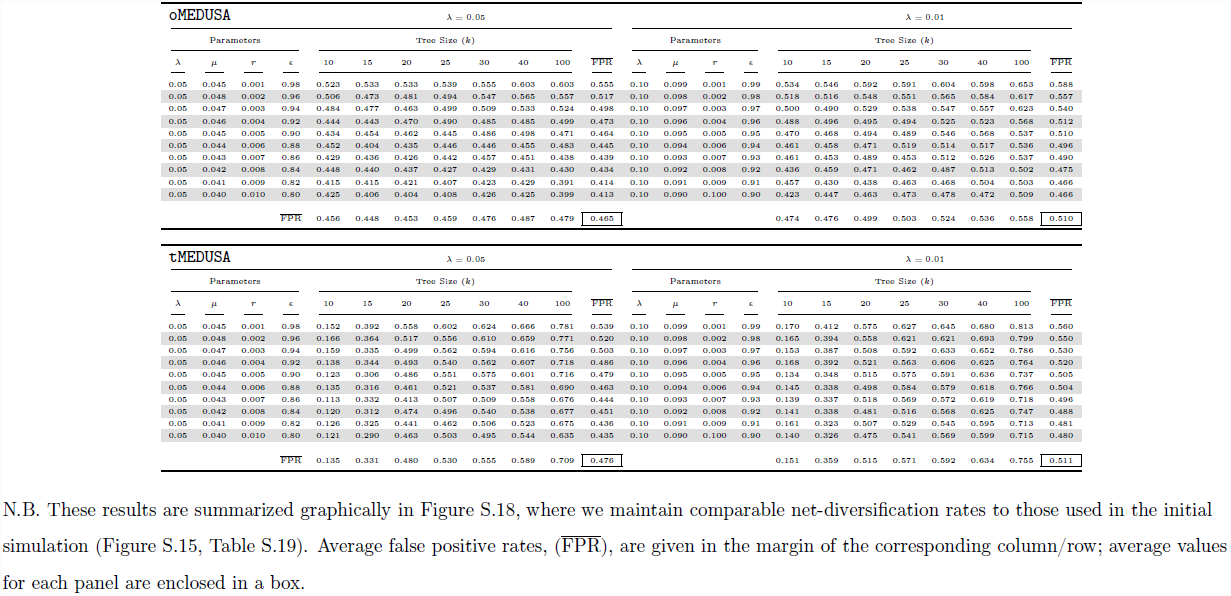
Parameters and Type I error rates for constant-rate simulations with 5-and 10-fold increases in absolute diversification rate.

### S.13 Exploring threshold effects on Type I error rates (crown shifts)

**Figure S.19:**
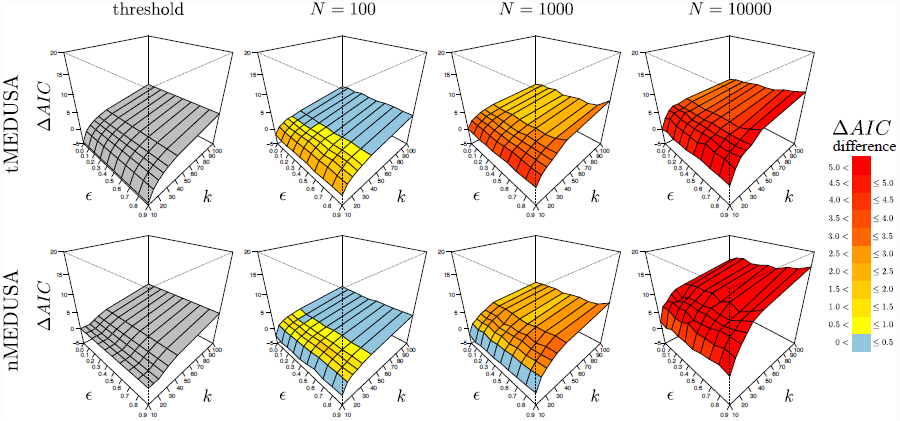
Critical and computed ∆AIC surfaces for two MEDUSA algorithms. The four panels in each row correspond to one algorithm (`tMEDUSA` and `nMEDUSA`, from top to bottom). The first column depicts the surface of ∆AIC_crit_ threshold values for each algorithm; when the difference between the AIC scores of two competing models exceeds this threshold, the more complex model is selected. Columns 2 − 4 summarize the ∆AIC_crit_ values computed from simulated trees of each size (*N* = 100, 1, 000, 10, 000). The computed ∆AIC surfaces are colored to reflect the degree to which they exceed the corresponding ∆AIC_crit_ threshold (see legend). Surfaces are computed from the trees used in the initial constant-rate simulation (*c.f.*, Figure S.15) under parameters described in Table S.19. For detailed values, see Table S.23

**Table S.23:**
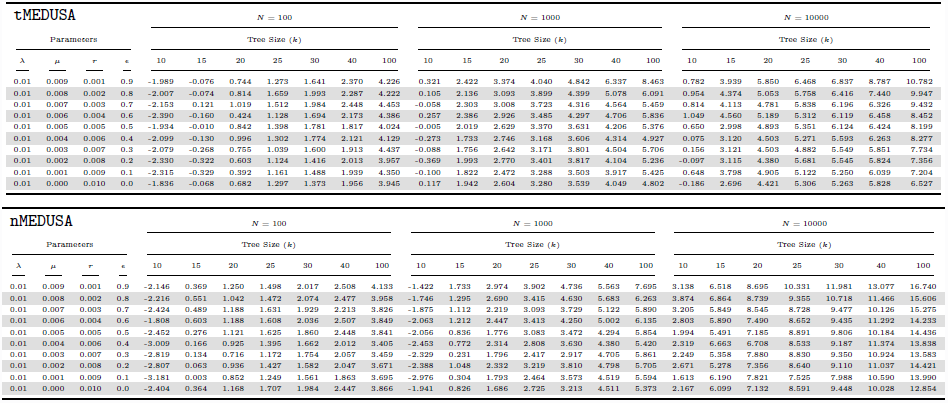
Empirical quantiles for ∆AICc (crown shifts).

### S.14 Exploring Type I error rates for complete species trees (crown shifts)

**Figure S.20:**
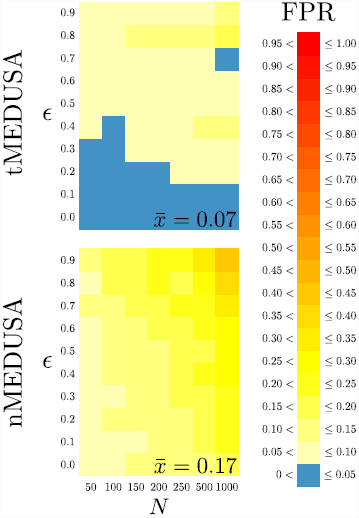
Type I error rates for completely sampled trees. Each row summarizes results for one algorithm (`tMEDUSA` and `nMEDUSA` on the top and bottom, respectively). Within each panel, we plot the number of species in the tree, *N*, against the relative-extinction rate used in the simulation. Trees were simulated under a constant-rate birth-death process using the parameters summarized in Table S.24. The cells within each panel are colored as a heat map reflecting the frequency with which spurious diversification-rate shifts were inferred (see FPR legend). The average Type I error rate is summarized in the lower right of each panel.

**Table S.24:**
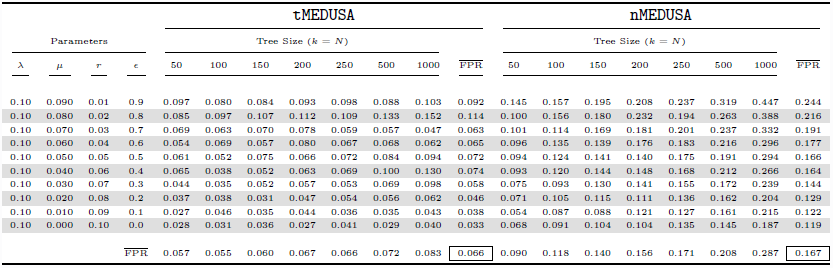
Results of the constant-rate simulation with completely sampled trees.

### S.15 Parameter estimation (crown shifts)

**Figure S.21:**
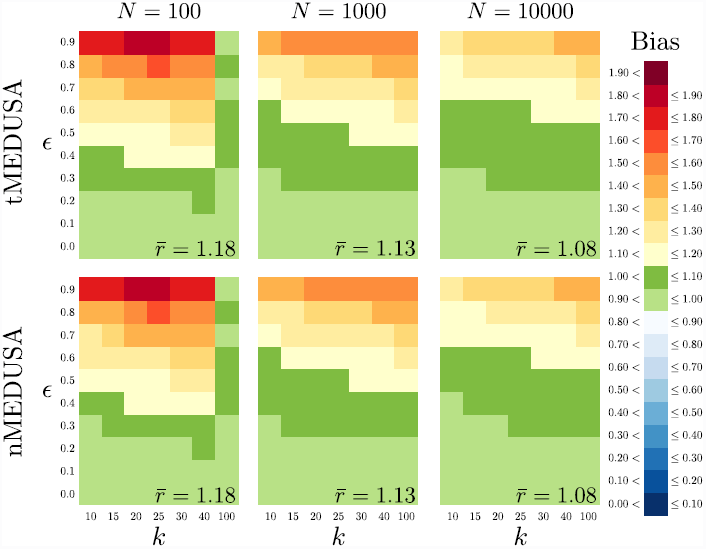
Bias in estimated net-diversification rate when a single-rate model is selected. The three panels in each row summarize results for one algorithm (`tMEDUSA` and `nMEDUSA`, from top to bottom), the three panels in each column summarize results for one tree size (with *N* = 100, 1, 000, 10, 000 species, from left to right). Within each panel, we plot the number of terminal lineages, *k*, against the relative-extinction rate used in the simulation. Trees were simulated under a constant-rate birth-death process using the parameters summarized in Table S.25. The cells within each panel are colored to reflect the average difference between the estimated and true net-diversification rates when a one-rate model is correctly identified (see Bias legend). The average bias in estimates of relative extinction-rate is summarized in the lower right of each panel.

**Table S.25:**
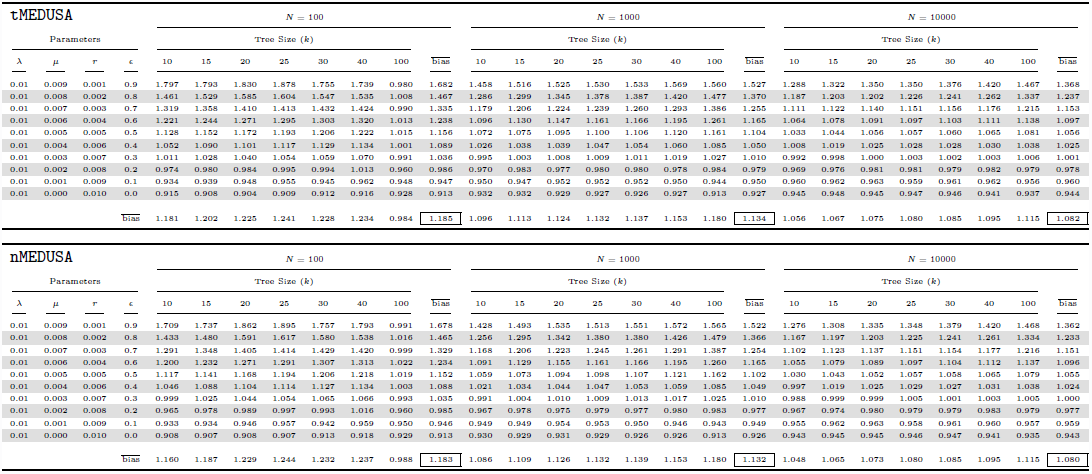
Estimated net-diversification rates for the initial constant-rate simulations when a single-rate model is selected.

**Figure S.22:**
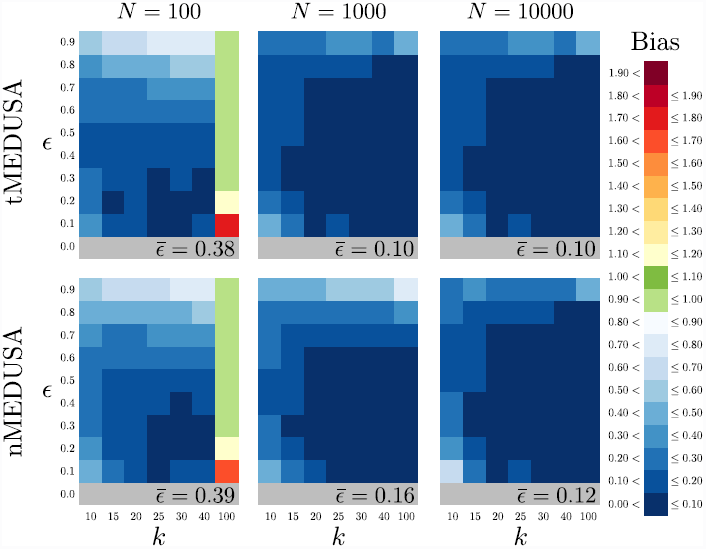
Bias in estimated relative-extinction rate when a single-rate model is selected. The three panels in each row summarize results for one algorithm (`tMEDUSA` and `nMEDUSA`, from top to bottom), the three panels in each column summarize results for one tree size (with *N* = 100, 1, 000, 10, 000 species, from left to right). Within each panel, we plot the number of terminal lineages, *k*, against the relative-extinction rate used in the simulation. Trees were simulated under a constant-rate birth-death process using the parameters summarized in Table S.12. The cells within each panel are colored to reflect the average difference between the estimated and true relative-extinction rates when a one-rate model is correctly identified (see Bias legend). The average bias in estimates of relative extinction-rate is summarized in the lower right of each panel.

**Table S.26:**
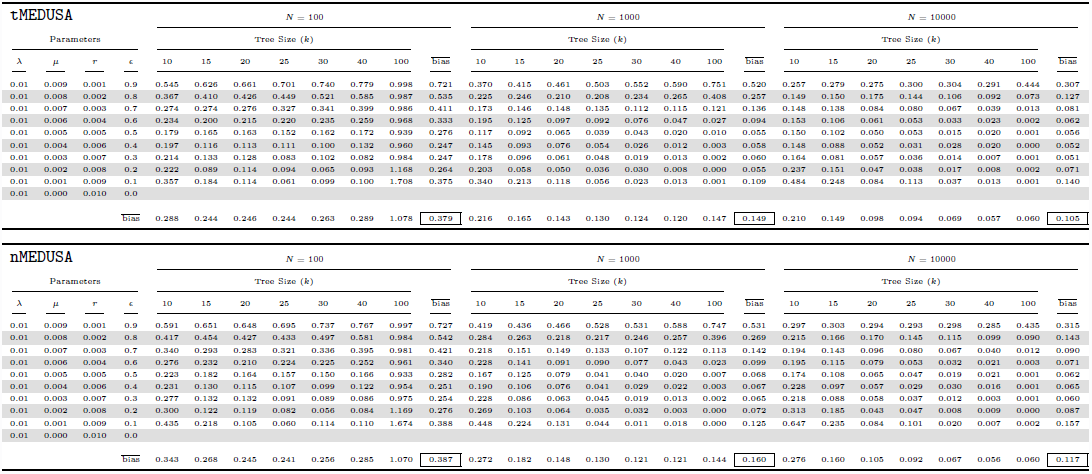
Estimated relative-extinction rates for the initial constant-rate simulations when a single-rate model is selected.

**Figure S.23:**
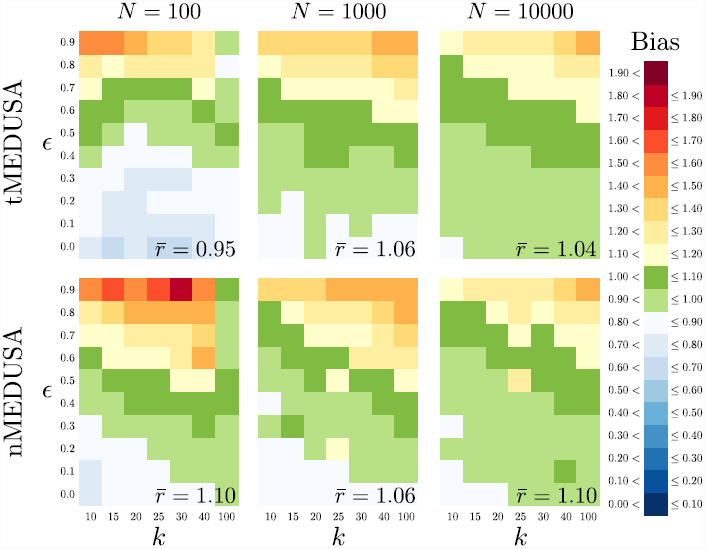
Bias in estimated background net-diversification rates when a two-rate model is selected. The three panels in each row summarize results for one algorithm (texttt`tMEDUSA`, and `nMEDUSA`, from top to bottom), the three panels in each column summarize results for one tree size (with *N* = 100, 1, 000, 10, 000 species, from left to right). Within each panel, we plot the number of terminal lineages, *k*, against the relative-extinction rate used in the simulation. Trees were simulated under a constant-rate birth-death process using the parameters summarized in Table S.27. The cells within each panel are colored to reflect the average difference between the estimated and true background net-diversification rate when a two-rate model is incorrectly identified (see Bias legend). The average bias in the inferred background net-diversification rate is summarized in the lower right of each panel.

**Table S.27:**
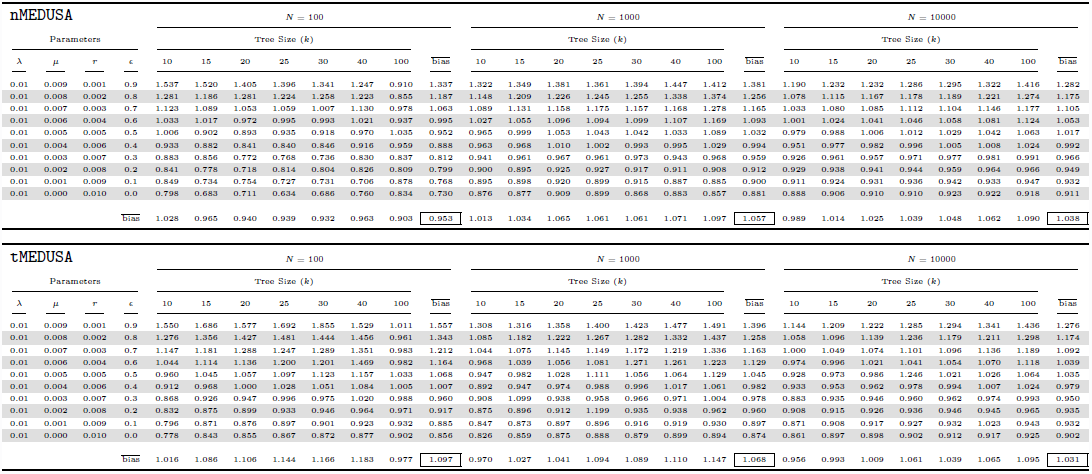
Estimated background net-diversification rates for the initial constant-rate simulations when a two-rate model is selected.

**Figure S.24:**
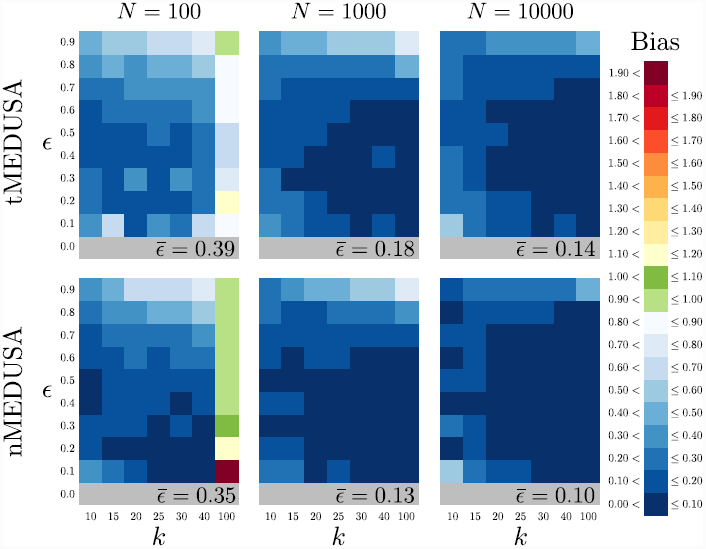
Bias in estimated background relative-extinction rates when a two-rate model is selected. The three panels in each row summarize results for one algorithm (`tMEDUSA`, and `nMEDUSA`, from top to bottom), the three panels in each column summarize results for one tree size (with *N* = 100, 1, 000, 10, 000 species, from left to right). Within each panel, we plot the number of terminal lineages, *k*, against the relative-extinction rate used in the simulation. Trees were simulated under a constant-rate birth-death process using the parameters summarized in Table S.28. The cells within each panel are colored to reflect the average difference between the estimated and true background relative-extinction rate when a two-rate model is incorrectly identified (see Bias legend). The average bias in the inferred background relative-extinction rate is summarized in the lower right of each panel.

**Table S.28:**
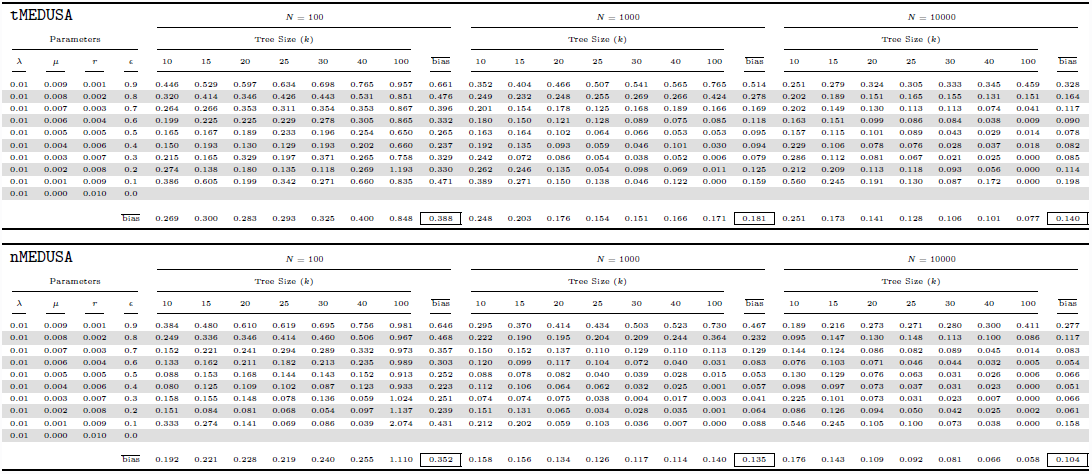
Estimated backgound relative-extinction rates for the initial constant-rate simulations when a two-rate model is selected.

**Figure S.25:**
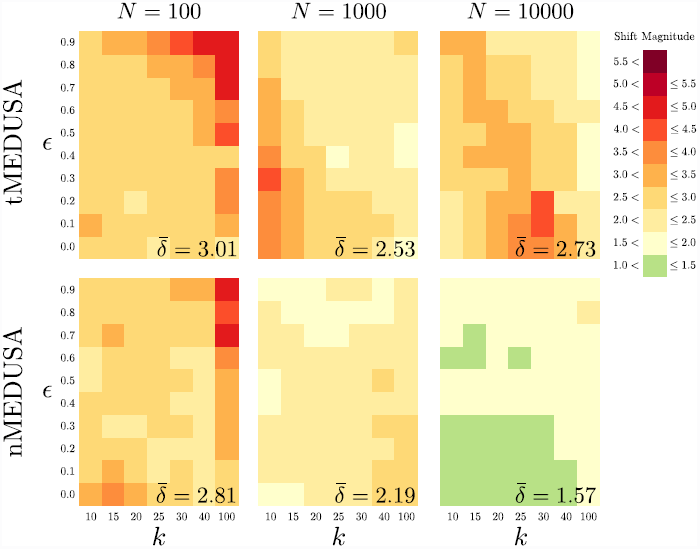
Magnitude of shifts in net-diversification rate when a two-rate model is chosen. The three panels in each row summarize results for one algorithm (`tMEDUSA`, and `nMEDUSA`, from top to bottom), the three panels in each column summarize results for one tree size (with *N* = 100, 1, 000, 10, 000 species, from left to right). Within each panel, we plot the number of terminal lineages, *k*, against the relative-extinction rate used in the simulation. Trees were simulated under a constant-rate birth-death process using the parameters summarized in Table S.29. The cells within each panel are colored as a heat map reflecting the average magnitude of shifts in net-diversification rate when a two-rate model is incorrectly identified (see Shift Magnitude legend). In order to accommodate the bimodal distribution of rate shifts, the magnitude of each net-diversification-rate shift was computed as the ratio of the higher rate to the lower rate, regardless of which rate corresponded to the background-rate category. The average magnitude of inferred diversification-rate shifts is summarized in the lower right of each panel.

**Table S.29:**
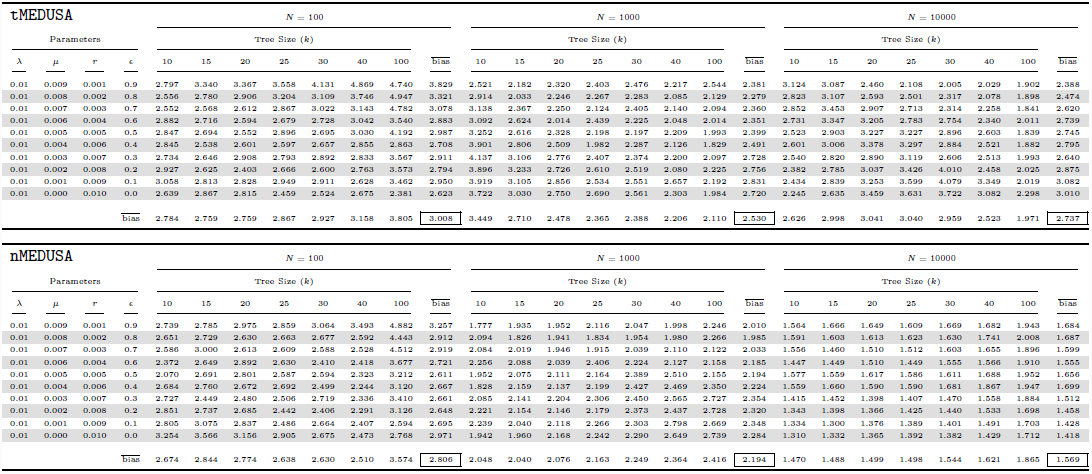
Estimated magnitude of net diversification-rate shifts for the initial constant-rate simulations when a two-rate model is selected.

**Figure S.26:**
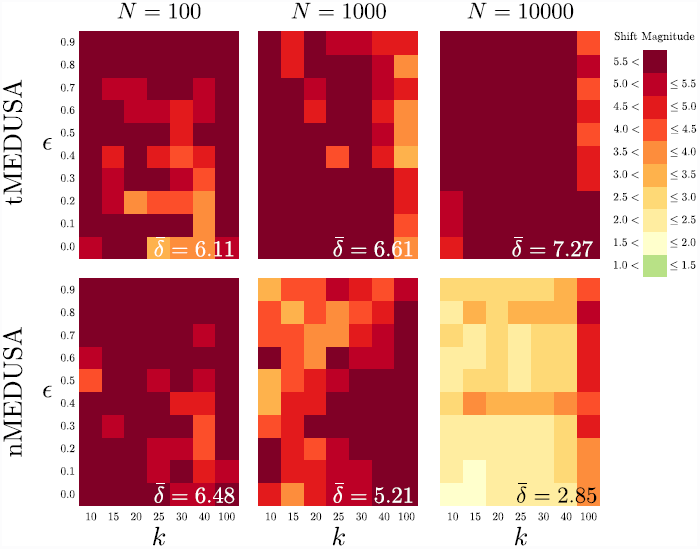
95% quantile of shift magnitude in net-diversification rate when a two-rate model is selected. The three panels in each row summarize results for one algorithm (`tMEDUSA` and `nMEDUSA`, from top to bottom), the three panels in each column summarize results for one tree size (with *N* = 100, 1, 000, 10, 000 species, from left to right). Within each panel, we plot the number of terminal lineages, *k*, against the relative-extinction rate used in the simulation. Trees were simulated under a constant-rate birth-death process using the parameters summarized in Table S.30. The cells within each panel are colored to reflect the 95% quantile of the magnitude of the inferred shift in net-diversification rate when a two-rate model is incorrectly identified (see Shift Magnitude legend). In order to accommodate the bimodal distribution of rate shifts, the magnitude of each net-diversification-rate shift was computed as the ratio of the higher rate to the lower rate, regardless of which rate corresponded to the background-rate category. The average 95% quantile of the magnitude of shifts in net-diversification rate shift is summarized in the lower right of each panel.

**Table S.30:**
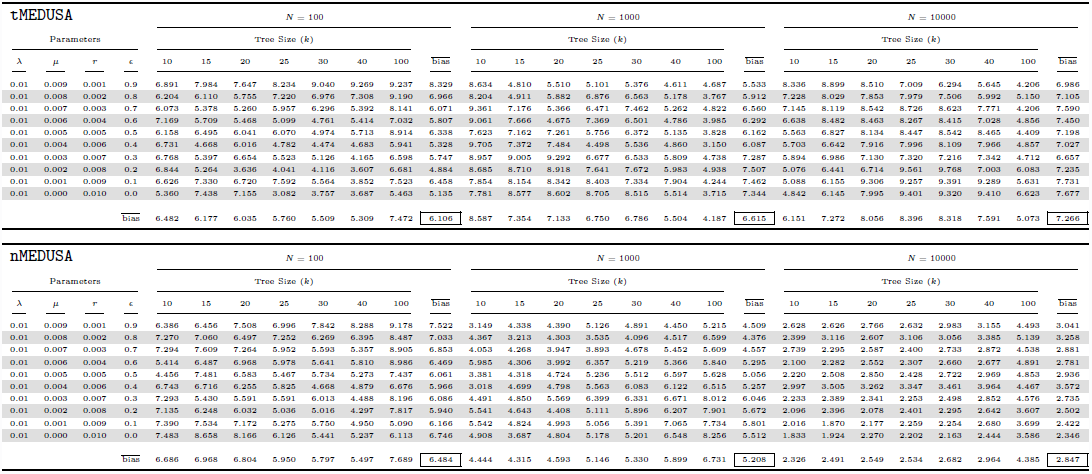
Estimated magnitude of net diversification-rate shifts for the initial constant-rate simulations when a two-rate model is selected.

| tMEDUSA |  |  |  |  |  |  |  |  |  |  |  |  |  |  |  |  |  |  |  |  |  |  |  |  |  |  |  |
| --- | --- | --- | --- | --- | --- | --- | --- | --- | --- | --- | --- | --- | --- | --- | --- | --- | --- | --- | --- | --- | --- | --- | --- | --- | --- | --- | --- |
| Parameters |  |  |  | N = 100 |  |  |  |  |  |  |  | N = 1000 |  |  |  |  |  |  |  |  |  |  |  |  |  |  |  |
|  |  |  |  | Tree Size (k) |  |  |  |  |  |  |  | Tree Size (k) |  |  |  |  |  |  |  |  |  |  |  |  |  |  |  |
| $\lambda$ | $\mu$ | r | $\epsilon$ | 10 | 15 | 20 | 25 | 30 | 40 | 100 | bias | 10 | 15 | 20 | 25 | 30 | 40 | 100 | bias | | | | | | | | |
| 0.01 | 0.009 | 0.001 | 0.9 | 6.891 | 7.984 | 7.647 | 8.234 | 9.040 | 9.269 | 9.237 | 8.329 | 8.634 | 4.810 | 5.510 | 5.101 | 5.376 | 4.611 | 4.687 | 5.533 | 8.336 | 8.899 | 8.510 | 7.009 | 6.294 | 5.645 | 4.206 | 6.986 |
| 0.01 | 0.008 | 0.002 | 0.8 | 6.204 | 6.110 | 5.755 | 7.220 | 6.976 | 7.308 | 9.190 | 6.966 | 8.204 | 4.911 | 5.882 | 6.876 | 6.563 | 5.178 | 3.767 | 5.912 | 7.228 | 8.029 | 7.853 | 7.979 | 7.506 | 5.992 | 5.150 | 7.105 |
| 0.01 | 0.007 | 0.003 | 0.7 | 6.073 | 5.378 | 5.260 | 5.957 | 6.296 | 5.392 | 8.141 | 6.071 | 9.361 | 7.176 | 5.366 | 6.471 | 7.462 | 5.262 | 4.822 | 6.560 | 7.145 | 8.119 | 8.542 | 8.726 | 8.623 | 7.771 | 4.206 | 7.590 |
| 0.01 | 0.006 | 0.004 | 0.6 | 7.169 | 5.709 | 5.468 | 5.099 | 4.761 | 5.414 | 7.032 | 5.807 | 9.061 | 7.666 | 4.675 | 7.369 | 6.501 | 4.786 | 3.985 | 6.292 | 6.638 | 8.482 | 8.463 | 8.267 | 8.415 | 7.028 | 4.856 | 7.450 |
| 0.01 | 0.005 | 0.005 | 0.5 | 6.158 | 6.495 | 6.041 | 6.070 | 4.974 | 5.713 | 8.914 | 6.338 | 7.623 | 7.162 | 7.261 | 5.756 | 6.372 | 5.135 | 3.828 | 6.162 | 5.563 | 6.827 | 8.134 | 8.447 | 8.542 | 8.465 | 4.409 | 7.198 |
| 0.01 | 0.004 | 0.006 | 0.4 | 6.731 | 4.668 | 6.016 | 4.782 | 4.474 | 4.683 | 5.941 | 5.328 | 9.705 | 7.372 | 7.484 | 4.498 | 5.536 | 4.860 | 3.150 | 6.087 | 5.703 | 6.642 | 7.916 | 7.996 | 8.109 | 7.966 | 4.857 | 7.027 |
| 0.01 | 0.003 | 0.007 | 0.3 | 6.768 | 5.397 | 6.654 | 5.523 | 5.126 | 4.165 | 6.598 | 5.747 | 8.957 | 9.005 | 9.292 | 6.677 | 6.533 | 5.809 | 4.738 | 7.287 | 5.894 | 6.986 | 7.130 | 7.320 | 7.216 | 7.342 | 4.712 | 6.657 |
| 0.01 | 0.002 | 0.008 | 0.2 | 6.844 | 5.264 | 3.636 | 4.041 | 4.116 | 3.607 | 6.681 | 4.884 | 8.685 | 8.710 | 8.918 | 7.641 | 7.672 | 5.983 | 4.938 | 7.507 | 5.076 | 6.441 | 6.714 | 9.561 | 9.768 | 7.003 | 6.083 | 7.235 |
| 0.01 | 0.001 | 0.009 | 0.1 | 6.626 | 7.330 | 6.720 | 7.592 | 5.564 | 3.852 | 7.523 | 6.458 | 7.854 | 8.154 | 8.342 | 8.403 | 7.334 | 7.904 | 4.244 | 7.462 | 5.088 | 6.155 | 9.306 | 9.257 | 9.391 | 9.289 | 5.631 | 7.731 |
| 0.01 | 0.000 | 0.010 | 0.0 | 5.360 | 7.438 | 7.155 | 3.982 | 3.757 | 3.687 | 5.463 | 5.135 | 7.781 | 8.577 | 8.602 | 8.705 | 8.515 | 5.514 | 3.715 | 7.344 | 4.842 | 6.145 | 7.995 | 9.401 | 9.320 | 9.410 | 6.623 | 7.677 |
| bias |  |  |  | 6.482 | 6.177 | 6.035 | 5.760 | 5.509 | 5.309 | 7.472 | 6.106 | 8.587 | 7.354 | 7.133 | 6.750 | 6.786 | 5.504 | 4.187 | 6.615 | 6.151 | 7.272 | 8.056 | 8.396 | 8.318 | 7.591 | 5.073 | 7.266 |

| nMEDUSA |  |  |  |  |  |  |  |  |  |  |  |  |  |  |  |  |  |  |  |  |  |  |  |  |  |  |  |
| --- | --- | --- | --- | --- | --- | --- | --- | --- | --- | --- | --- | --- | --- | --- | --- | --- | --- | --- | --- | --- | --- | --- | --- | --- | --- | --- | --- |
| Parameters |  |  |  | N = 100 |  |  |  |  |  |  |  | N = 1000 |  |  |  |  |  |  |  |  |  |  |  |  |  |  |  |
|  |  |  |  | Tree Size (k) |  |  |  |  |  |  |  | Tree Size (k) |  |  |  |  |  |  |  |  |  |  |  |  |  |  |  |
| $\lambda$ | $\mu$ | r | $\epsilon$ | 10 | 15 | 20 | 25 | 30 | 40 | 100 | bias | 10 | 15 | 20 | 25 | 30 | 40 | 100 | bias | | | | | | | | |
| 0.01 | 0.009 | 0.001 | 0.9 | 6.386 | 6.456 | 7.508 | 6.996 | 7.842 | 8.288 | 9.178 | 7.522 | 3.149 | 4.338 | 4.390 | 5.126 | 4.891 | 4.450 | 5.215 | 4.509 | 2.628 | 2.626 | 2.766 | 2.632 | 2.983 | 3.155 | 4.493 | 3.041 |
| 0.01 | 0.008 | 0.002 | 0.8 | 7.270 | 7.060 | 6.497 | 7.262 | 6.269 | 6.395 | 8.487 | 7.033 | 3.867 | 3.213 | 3.303 | 3.535 | 4.096 | 4.517 | 6.599 | 4.376 | 2.399 | 3.116 | 2.607 | 3.106 | 3.056 | 3.385 | 5.139 | 3.258 |
| 0.01 | 0.007 | 0.003 | 0.7 | 7.294 | 7.609 | 7.264 | 5.952 | 5.593 | 5.357 | 8.905 | 6.853 | 4.053 | 4.268 | 3.947 | 3.893 | 4.678 | 5.452 | 5.609 | 4.557 | 2.739 | 2.295 | 2.587 | 2.400 | 2.733 | 2.872 | 4.538 | 2.881 |
| 0.01 | 0.006 | 0.004 | 0.6 | 5.414 | 6.487 | 6.968 | 5.378 | 5.641 | 5.510 | 8.966 | 6.469 | 5.985 | 4.366 | 3.992 | 6.357 | 5.219 | 3.366 | 5.840 | 5.295 | 2.100 | 2.282 | 2.352 | 2.307 | 2.660 | 2.677 | 4.891 | 2.781 |
| 0.01 | 0.005 | 0.005 | 0.5 | 4.456 | 7.481 | 6.583 | 5.467 | 5.734 | 5.273 | 7.437 | 6.061 | 3.381 | 4.318 | 4.724 | 5.236 | 5.512 | 6.597 | 5.628 | 5.056 | 2.220 | 2.508 | 2.850 | 2.428 | 2.722 | 2.969 | 4.853 | 2.936 |
| 0.01 | 0.004 | 0.006 | 0.4 | 6.743 | 6.716 | 6.255 | 5.825 | 4.668 | 4.879 | 6.676 | 5.966 | 3.018 | 4.699 | 4.798 | 5.563 | 6.083 | 6.122 | 6.515 | 5.257 | 2.997 | 3.505 | 3.262 | 3.347 | 3.461 | 3.964 | 4.467 | 3.572 |
| 0.01 | 0.003 | 0.007 | 0.3 | 7.293 | 5.430 | 5.591 | 5.591 | 6.013 | 4.488 | 8.196 | 6.086 | 4.491 | 4.850 | 5.569 | 6.399 | 6.331 | 6.671 | 8.012 | 6.046 | 2.233 | 2.389 | 2.341 | 2.253 | 2.498 | 2.852 | 4.576 | 2.735 |
| 0.01 | 0.002 | 0.008 | 0.2 | 7.135 | 6.248 | 6.032 | 5.036 | 5.016 | 4.297 | 7.817 | 5.940 | 5.541 | 4.643 | 4.408 | 5.111 | 5.896 | 6.207 | 7.901 | 5.672 | 2.096 | 2.396 | 2.078 | 2.401 | 2.295 | 2.642 | 3.607 | 2.502 |
| 0.01 | 0.001 | 0.009 | 0.1 | 7.390 | 7.534 | 7.172 | 5.275 | 5.750 | 4.950 | 5.090 | 6.166 | 5.542 | 4.824 | 4.993 | 5.056 | 5.391 | 7.065 | 7.734 | 5.801 | 2.016 | 1.870 | 2.177 | 2.259 | 2.254 | 2.680 | 3.699 | 2.422 |
| 0.01 | 0.000 | 0.010 | 0.0 | 7.483 | 8.658 | 8.166 | 6.126 | 5.441 | 5.237 | 6.113 | 6.746 | 4.908 | 3.687 | 4.804 | 5.178 | 5.201 | 6.548 | 8.256 | 5.512 | 1.833 | 1.924 | 2.270 | 2.202 | 2.163 | 2.444 | 3.586 | 2.346 |
| bias |  |  |  | 6.686 | 6.968 | 6.804 | 5.950 | 5.797 | 5.497 | 7.689 | 6.484 | 4.444 | 4.315 | 4.593 | 5.146 | 5.330 | 5.899 | 6.731 | 5.208 | 2.326 | 2.491 | 2.549 | 2.534 | 2.682 | 2.964 | 4.385 | 2.847 |

**Figure S.27:**
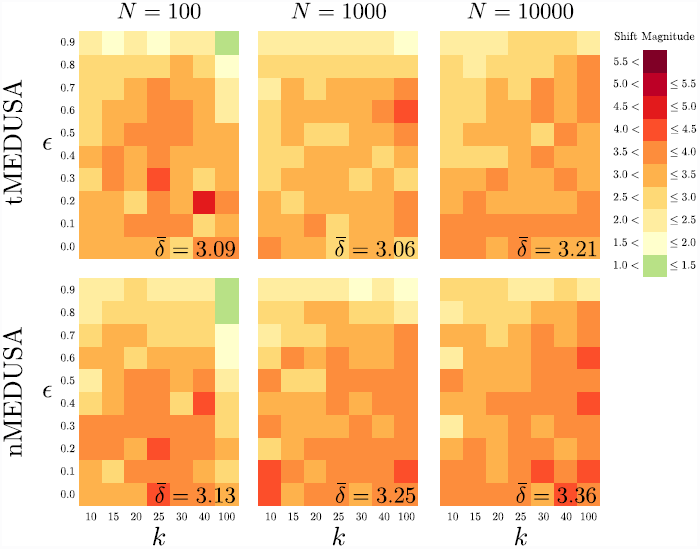
Magnitude of shift in relative-extinction rate when a two-rate model is selected. The three panels in each row summarize results for one algorithm (`tMEDUSA`, and `nMEDUSA`, from top to bottom), the three panels in each column summarize results for one tree size (with *N* = 100, 1, 000, 10, 000 species, from left to right). Within each panel, we plot the number of terminal lineages, *k*, against the relative-extinction rate used in the simulation. Trees were simulated under a constant-rate birth-death process using the parameters summarized in Table S.31. The cells within each panel are colored to reflect the average magnitude of the inferred shift in relative-extinction rate when a two-rate model is incorrectly identified (see Shift Magnitude legend). In order to accommodate the bimodal distribution of rate shifts, the magnitude of each relative-extinction-rate shift was computed as the ratio of the higher rate to the lower rate, regardless of which rate corresponded to the background-rate category. The average magnitude of shifts in relative-extinction rate shift is summarized in the lower right of each panel.

**Table S.31:**
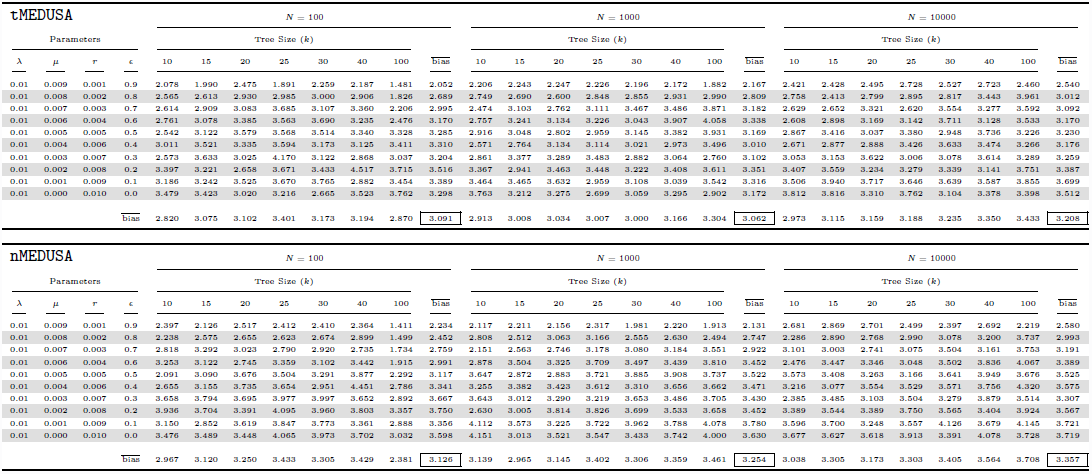
Estimated magnitude of relative-extinction rate shifts for the initial constant-rate simulations when a two-rate model is selected.

**Figure S.28:**
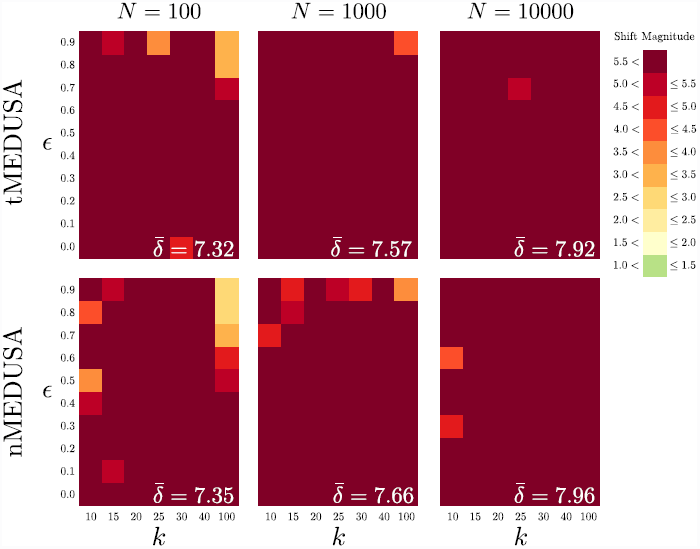
95% quantile of shift magnitude in relative-extinction rate when a two-rate model is selected. The three panels in each row summarize results for one algorithm (`tMEDUSA` and `nMEDUSA`, from top to bottom), the three panels in each column summarize results for one tree size (with *N* = 100, 1, 000, 10, 000 species, from left to right). Within each panel, we plot the number of terminal lineages, *k*, against the relative-extinction rate used in the simulation. Trees were simulated under a constant-rate birth-death process using the parameters summarized in Table S.32. The cells within each panel are colored to reflect the 95% quantile of the magnitude of the inferred shift in relative-extinction rate when a two-rate model is incorrectly identified (see Shift Magnitude legend). In order to accommodate the bimodal distribution of rate shifts, the magnitude of each relative-extinction-rate shift was computed as the ratio of the higher rate to the lower rate, regardless of which rate corresponded to the background-rate category. The average 95% quantile of the magnitude of shifts in relative-extinction rate shift is summarized in the lower right of each panel.

**Table S.32:**
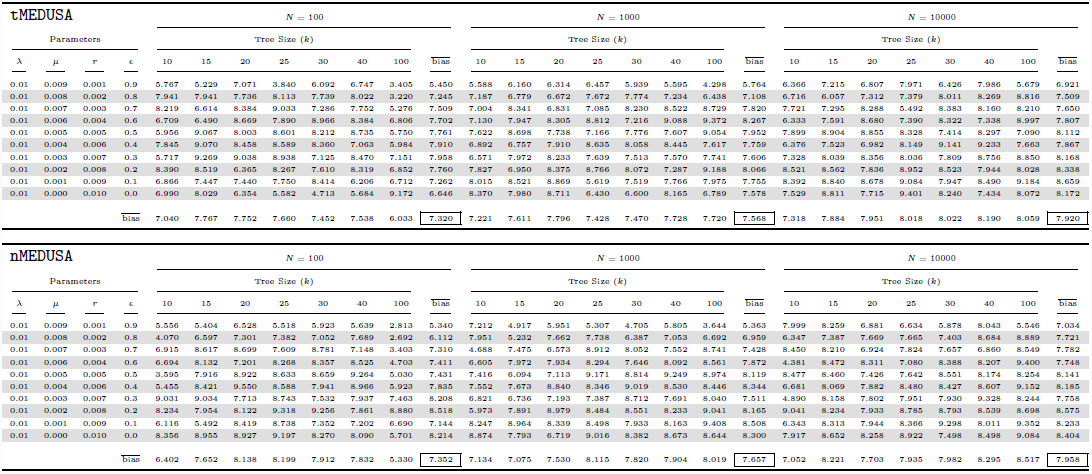
95% quantile of estimated magnitude of relative-extinction-rate shifts for the initial constant-rate simulations when a two-rate model is selected.

